# *ST3GAL3* loss-of-function disrupts synaptic integrity and excitatory/inhibitory cortical dynamics

**DOI:** 10.64898/2026.06.21.733355

**Authors:** David Diouf, Dorothea Lambrini Tsounis, Ehsan Pishva, Tim Vanmierlo, Daniel van den Hove, Klaus-Peter Lesch

## Abstract

This study examined the role of ST3GAL3 as a regulator of excitatory/inhibitory (E/I) synaptic homeostasis using a human iPSC-based model. Neurodevelopmental disorders (NDDs) are increasingly linked to disruptions in the E/I balance, yet the molecular determinants remain poorly defined. ST3GAL3, a sialyltransferase associated with both rare monogenic disorders, including intellectual disability and infantile epilepsy, and complex polygenic conditions, such as ADHD, represents a strong candidate gene for involvement in synaptic regulation. To investigate this, isogenic ST3GAL3 knockout (ST3GAL3 KO) and wildtype (WT) iPSC lines were generated through CRISPR/Cas9 editing and differentiated into cortical neurons using both directed and induced protocols. This dual strategy enabled robust comparisons across cellular contexts and minimised methodological bias. To this end, we conducted functional characterisation using microelectrode array (MEA) technology alongside transcriptomic profiling through RNA sequencing (RNAseq), directly comparing ST3GAL3 KO-derived neurons with their isogenic controls. Functional assays using MEA revealed aberrant bursting patterns, particularly prolonged burst durations and heightened variability in S3GAL3*^KO^* neurons. Complementary transcriptomic profiling performed via RNAseq demonstrated downregulation in ST3GAL3 KO lines of genes involved in cognition, memory, as well as glutamatergic and GABAergic synaptic plasticity and functionality, providing molecular evidence for widespread synaptic dysregulation. Together, these findings establish ST3GAL3 as a key regulator of E/I balance in the cortices, advancing current knowledge on the pathophysiological involvement of ST3GAL3 deficiencies in the development of NDDs.

## Introduction

Neurodevelopmental disorders (NDDs) represent a heterogeneous group of conditions that arise from disrupted development of the central nervous system, typically manifesting in early childhood. These include autism spectrum disorder (ASD), attention-deficit/hyperactivity disorder (ADHD), and intellectual disability (ID), among others [1, 2]. NDDs are highly prevalent, affecting an estimated 18.3% of children and adolescents worldwide, and constitute a major public health challenge [3, 4]. Clinically, NDDs profoundly affect an individual’s autonomy, and quality of life, carrying deficits in cognitive function, motor coordination, emotion regulation, communication, and social interaction [1]. Many NDDs are chronic and persist into adulthood, often accompanied by psychiatric comorbidities such as depression, anxiety and substance use disorders [5]. Despite increasing awareness and ongoing research efforts, current therapeutic options remain largely symptomatic, highly variable, and of poor long-term efficacy.

The aetiology of NDDs is complex and multifactorial, involving a combination of genetic susceptibility and environmental adversity. NDDs present with considerable phenotypic and pathophysiological overlap. However, they also exhibit substantial variability in onset, severity, and progression, making it particularly difficult to categorise them into distinct subgroups or define disorder-specific mechanisms [2, 6–8]. This complexity continues to hinder efforts to identify suitable diagnostic markers and therapeutic targets.

Research has identified an increasing amount of shared neurobiological mechanisms that transcend traditional diagnostic boundaries [9, 10]. Among the most widely recognised mechanisms is the disruption of the excitatory/inhibitory (E/I) balance within neuronal networks [11–22]. This balance refers to the dynamic interplay between excitatory glutamatergic and inhibitory gamma-aminobutyric acid (GABA) neurotransmission [23–25]. Alterations in the E/I equilibrium were first implicated in ASD [26] and have since been reported across a range of NDDs, including ADHD [27–31], ID [11–13], and epilepsy [14–18]. Moreover, growing evidence suggests that E/I imbalance may also contribute to conditions not traditionally classified as developmental, such as psychotic disorders, further underscoring its broad relevance to brain dysfunction [32–35]. Subsequent research has refined the concept of E/I balance into a dynamic and multidimensional framework, ranging across multiple spatiotemporal scales and accompanying diversities that may underlie the broad clinical variability and heterogeneous manifestations observed across NDDs. Given its relevance, it becomes increasingly important to identify and characterise the molecular players that govern and perturb the regulation of the E/I balance.

Among the genes implicated in NDDs, *ST3GAL3* has emerged as a particularly compelling candidate. Loss-of-function mutations in *ST3GAL3* have been associated with non-syndromic autosomal recessive ID (NSARID) and early infantile epileptic encephalopathy (EIEE) [36–39], while genome-wide association studies (GWAS) have identified *ST3GAL3* as the top risk locus for ADHD [40]. Notably, *ST3GAL3* was also found to be associated with general cognitive function outcomes, hence implicating *ST3GAL3* as a contributing factor to the cognitive variation within the normal-range driven by common genetic variation [41]. *ST3GAL3* encodes the homonym sialyltransferase responsible for the sialylation of glycoproteins, a post-translational modification critical for cell adhesion processes [42–44]. ST3GAL3 represents the initiating step in the biosynthetic pathway leading to polysialylated glycoproteins, such as neural cell adhesion molecule 1 (NCAM1) and synaptic cell adhesion molecule 1 (SynCAM1, CADM1), both of which play critical roles in synaptic development and plasticity [45–50]. Despite these associations, remarkably little is known about the precise role of ST3GAL3 in brain development and function, and virtually nothing is known about its contribution to the E/I balance. We, therefore, sought to investigate the role of *ST3GAL3* in this context.

Our work was guided by the central hypothesis that disruption of *ST3GAL3* impairs synaptic regulation in cortical glutamatergic and GABAergic neurons, thereby altering the E/I balance of neuronal networks. To investigate the role of *ST3GAL3* in a human-based context, a *ST3GAL3* knockout (KO) induced pluripotent stem cell (iPSC) line was generated as previously described [51], making use of CRISPR/Cas9 genome editing in a healthy donor-derived iPSC line. The use of a *ST3GAL3* KO an isogenic wild-type iPSC line enables the investigation of *ST3GAL3* inactivation within a genetically uniform background, providing a highly controlled experimental setting. This approach minimises confounding genetic variability and offers a powerful platform to precisely attribute observed phenotypic effects to the loss of ST3GAL3.

In the present work, we leveraged this model to study the impact of ST3GAL3 deficiency on human cortical neurons. We employed both directed and induced differentiation protocols—the two most widely adopted approaches for generating iPSC-derived cortical neurons. This strategy allowed us to compare findings across distinct differentiation paradigms and strengthen the robustness of our conclusions. In doing so, we characterised and compared lines with a combination of molecular, morphological, and functional assays. At the functional level, we employed high-throughput electrophysiological recordings using microelectrode array (MEA) technology to capture network-level activity with high temporal resolution. At the molecular level, we performed transcriptomic profiling via RNA sequencing (RNAseq) to identify gene expression changes associated with *ST3GAL3* deficiency.

Our findings demonstrate that sialylation deficits profoundly impact neuronal network function and gene expression. MEA analyses revealed hyperexcitability in *ST3GAL3 KO* cultures, marked by prolonged and irregular burst activity, absent in isogenic controls. Complementary transcriptomic profiling identified robust and reproducible downregulation of genes implicated in both glutamatergic and GABAergic signalling, suggesting widespread synaptic dysregulation. Collectively, these findings uncover a previously unrecognised role for *ST3GAL3* in maintaining synaptic integrity and E/I balance, thereby offering mechanistic insight into how sialylation deficits may contribute to NND pathophysiology.

## Materials and methods

### Generation and maintenance of human iPSC lines

Human iPSCs (hiPSCs) used in this study were derived from dermal fibroblasts of a neurotypical donor, as previously described. The *ST3GAL3* null mutant line (referred in this thesis as *ST3GAL3*^−/−^; ST3GAL3 KO; UKWMPi002-A-3) was generated via CRISPR/Cas9 genome editing, introducing frameshift mutations within exon 2, leading to complete loss of functional *ST3GAL3* transcripts. The isogenic wild-type (WT) iPSC line, served as the control throughout all experiments. Both lines were routinely cultured on Geltrex (Gibco) or Matrigel (Corning)-coated six-well plates in StemMAC iPSC-Brew XF medium (Miltenyi). iPSCs were maintained at 37°C with 5% CO₂ and passaged at 80-90% confluency (every 3–4 days) using Accutase (Sigma-Aldrich). All iPSC lines were confirmed to express standard pluripotency markers (OCT3/4, NANOG, SSEA4, TRA-1-60) via immunocytochemistry and RT-qPCR and displayed a normal karyotype. Routine checks for mycoplasma contaminations were performed using the LookOut Mycoplasma PCR Detection Kit (Sigma-Aldrich) (data not shown).

In this study, the human iPSC lines employed to investigate the role of ST3GAL3 in regulating the E/I balance in the developing cortex are referred to as: ST3GAL3 knockout (*ST3GAL3*^−/−^; ST3GAL3 KO), and its isogenic wild-type (WT) control (*ST3GAL3*^+/+;^ ST3GAL3 WT).

### Neural differentiation approaches

Two complementary strategies were adopted to derive cortical neurons from iPSCs: an induced differentiation protocol based on the overexpression of lineage-specifying transcription factors neurogenin-2 (Ngn2) and achaete-scute family BHLH transcription factor 1 (Ascl1), and a developmentally guided, directed differentiation protocol involving dual inhibition of the “small mothers against decapentaplegic” (SMAD). The induced protocol relied on lentiviral transduction of transcription factors under doxycycline control, whereas the directed differentiation protocol involved solely the timed exposure of iPSCs to small-molecule inhibitors and growth factors in the culture medium. In the following sections, we first describe the induced protocol, followed by dedicated sections on the directed protocol. Regardless of the protocol adopted, ST3GAL3 KO and WT lines were always differentiated in parallel to ensure minimal technical variability.

### Induced differentiation

#### Lentiviral engineering for inducible neuronal differentiation

To enable inducible neuronal conversion, iPSC lines were stably transduced with lentiviral vectors encoding either Ngn2 or Ascl1 under control of a tetracycline-responsive promoter (TRE3G), alongside the reverse tetracycline-controlled transactivator (rtTA3). Lentiviral production and transduction were performed following the protocol described by Frega et al. (2017). In brief, HEK293T cells were transfected with plasmids encoding the gene of interest (rtTA, Ngn2, or Ascl1) (plasmids kindly provided by research group of Cornelis A. Albers and Nael Nadif Kasri, Radboud University, Nijmegen, the Netherlands), together with the packaging plasmid psPAX2 (Addgene) and the envelope plasmid pMD2.G (Addgene). Transfection was performed using JetPrime reagent (Polyplus) and viral particles were harvested from the supernatant 48 hours post-transfection. The supernatants were filtered through 0.45 µm syringe filters to isolate the lentiviral vectors, which were either used immediately for transduction or stored at –80°C. Lentiviral supernatants containing rtTA, Ngn2, or Ascl1 constructs were subsequently used to transduce iPSCs. Six hours post-transduction, the medium was replaced with fresh StemMACS medium containing 10 μM Y27632 (Miltenyi). Transduced iPSCs were then subjected to dual antibiotic selection with puromycin (Invivogen; 7 μg/ml) and G418 (Sigma;35 μg/ml) over a 5-day period. Transfected hiPSCs were then cultured on matrigel-coated 6-well plates in StemMACS medium supplemented with G418 (50 mg/mL), and puromycin (0.5 mg/mL) and maintained as described above. Stable integration of transgenes was verified by PCR-based genotyping and qPCR assessment of transcript induction dynamics. The absence of lentiviral particles in the cultures was confirmed after four to five passages by genomic PCR on both the adherent cells and the culture supernatants. Through this methodology, both the Ngn2 and Ascl1 constructs are placed under control of a Tet-responsive promoter, activated by the doxycycline-inducible rtTA. Upon doxycycline treatment, rtTA binds the Tet promoter and initiates transcription of Ngn2 or Ascl1, thereby committing the iPSCs to glutamatergic or GABAergic fates, respectively.

#### Co-culture of inducible glutamatergic and GABAergic neurons

On day 0, rtTA/Ngn2– and rtTA/Ascl1-positive cells were brought in suspension with Accutase, mixed in a 65:35 ratio and plated on poly-L-ornithine (15 μg/ml, Sigma) and laminin (10 μg/ml, SIgma)-coated plates or MEA chips at a density of 1,7×10^4^ cells per cm^2^ in StemMACS medium supplemented with 10 μM Y27632. On Day 1, the medium was replaced with DMEM/F12 supplemented with N2 (1:100, ThermoFisher Scientific), non-essential amino acids (NEAA; 1:100, Sigma), primocin (1:500, ThermoFisher Scientific), human recombinant brain-derived neurotrophic factor (BDNF; 10 ng/ml, PeproTech), NT-3 (10 ng/ml, Promocell), laminin (0.2 μg/ml, Biolamina), doxycycline (4 μg/ml, Sigma), and forskolin (10 μM, *Sigma*). On day 3, cytosine b-D-arabinofuranoside hydrochloride (Ara-C) was added to remove non-differentiated hiPSCs left in the culture and inhibit the proliferation. To this extent, medium was refreshed with neurobasal medium (ThermoFisher Scientific) supplemented with Ara-C (2µM, *Sigma*), B27 (1:50, ThermoFisher Scientific), Glutamax (1:100, ThermoFisher Scientific), primocin (1:500), NT3 (10 ng/ml), BDNF (10 ng/ml), forskolin (10 µM), and doxycycline (4µg/ml). From Day 5 onward, ara-C could be removed, and the cultures were maintained in neurobasal medium supplemented with B27 (1:50), glutaMAX (1:100), primocin (1:500), BDNF (10 ng/ml), NT-3 (10 ng/ml), doxycycline (4 µg/ml), and forskolin (10 µM). Half of the medium was refreshed every 2-3 days. Starting from day 13 onwards, doxycycline was removed from the medium. Neuronal cultures were kept through the whole differentiation process at 37°C/ 5% CO_2_.

### Directed differentiation

#### Generation of *neural progenitor cells (*NPC)s via embryoid body-based dual-SMAD inhibition

Unless stated otherwise, all reagents and media used in this protocol were obtained from STEMCELL Technologies. We employed the directed differentiation approach using the STEMdiff neural induction protocol. On day 0, hiPSCs were dissociated into single cells using Gentle Cell Dissociation Reagent and suspended in STEMdiff Neural Induction Medium (NIM) supplemented with SMADi Neural Induction Supplement and Y-27632. A total of 3 x 10^6^ cells per well were seeded in AggreWell-800 plates to generate uniformly sized embryoid bodies (EBs). Between days 1-4, 3/4 of the medium was refreshed daily. On day 5, EBs were collected using a 37 µm Reversible Strainer and replated onto matrigel-coated 6 well-plates. Over the following 6 days, we performed full medium changes with NIM and SMADi. During this phase, radial organisation and rosette formation could be observed, indicative of neuroepithelial differentiation. On day 8, percentage of neural induction was calculated by counting the EBs in which 50% or more of the area of each individual aggregate was filled with neural rosettes. Only replicates with 100% of neural induction were kept, while the rest was discarded. On day 12, neural rosettes were selected using STEMdiff Neural Rosette Selection Reagent and expanded on matrigel-coated plates in NIM to yield stable populations of neural progenitor cells (NPCs). These NPCs were passaged using Accutase (at approximately 80% confluence) and maintained in STEMdiff Neural Progenitor Medium (NPM). NPCs could then be plated for cortical neuron maturation or frozen, using STEMdiff Neural Progenitor Freezing Medium. Typical forebrain-patterned NPC colonies were observed after 2-4 passages.

#### Directed differentiation and maturation into cortical neurons

To achieve cortical neuron differentiation, NPCs were seeded onto poly-L-ornithine/laminin-coated culture plates at a density of 3 x 10⁴ cells/cm². From the next day, cultures were maintained in BrainPhys Neuronal Medium supplemented with NeuroCult SM1 Neuronal Supplement (1:50), N2 Supplement-A (1:100), human recombinant BDNF (20 ng/mL), human recombinant glial cell line-derived neurotrophic factor (GDNF; 20 ng/mL), dibutyryl-cAMP (1 mM), and ascorbic acid (200 nM). Medium was refreshed every 2–3 days via half-medium changes. Neuronal maturation and synaptogenesis became evident from day 14 onward, and cultures were maintained up to 35 days for all downstream analyses. Neuronal cultures were kept through the whole differentiation process at 37°C and 5% CO_2_.

### Immunocytochemistry

For iPSCs and NPCs, cells were rinsed with Dulbecco’s phosphate buffered saline (dPBS, Gibco) and fixed with 4% paraformaldehyde (PFA, RotiHistofix, Roth) for 20 minutes at room temperature (RT). For neurons, to minimise detachment during fixation, half of the culture medium was first removed and replaced with dPBS. This dPBS wash was repeated three times. Subsequently, half of the medium was removed and replaced with 4% PFA for 5 minutes. Following this pre-fixation step, 3/4 of the total volume was replaced with fresh 4% PFA, and neurons were incubated at RT for an additional 15 minutes. The subsequent steps were identical across all cell types. To reduce non-specific antibody binding, cells were incubated with blocking buffer (1 hour, RT), composed of dPBS supplemented with bovine-serum albumin (1%, Sigma-Aldrich), and fetal bovine serum (10%, ThermoFisher). When using antibodies targeting intracellular epitopes, the blocking buffer was supplemented with 0.1% Triton X-100 (Sigma-Aldrich) to permeabilise cell membranes. Following the blocking step, primary antibodies were diluted in blocking buffer, and the antibody solution was added to the dedicated wells. Plates were sealed with parafilm, and incubated overnight at 4°C. The next day, cells were washed thrice with dPBS for 5 minutes at RT and incubated with the secondary antibody dilution, composed of the blocking buffer supplemented with the appropriate Alexa Fluor-conjugated secondary antibodies and DAPI nuclear stain (Invitrogen). Cells were washed three more times with dPBS for 5 minutes at RT, leaving dPBS in each well after the last wash. Plates were then sealed with parafilm and stored at 4°C. Imaging was performed using an Olympus IX81 inverted fluorescence microscope. Images were then processed using software CellSense (Olympus) and Fiji [52].

### RNA extraction

Prior to extraction, neuronal and NPC cultures were washed with dPBS and collected using a cell scraper. We then performed total RNA extraction using the miRNEasy Micro kit (Qiagen), following manufacturer’s instructions. RNA was eluted in RNAse-free water, analysed with a Nanophotometer N60 (Implen), and stored at –80°C for qRT-PCR or RNAseq.

For iPSCs, cells were collected using Accutase, resuspended in dPBS, and cells were mechanically lysed using sterile stainless-steel beads in a TissueLyser II (Qiagen) at 20 Hz for 30 seconds. Total RNA was extracted using the RNeasy Plus Mini Kit (Qiagen). To eliminate genomic DNA contamination, lysates were incubated with DNase dissolved in RDD buffer (Qiagen) within the spin columns for 15 minutes at RT. The remaining steps were carried out according to the manufacturer’s instructions. RNA was eluted in RNase-free water, analysed with the Nanophotometer, and stored at –80 °C for qRT-PCR.

### Genotyping

Genomic DNA from all types of cells was extracted using the PureLink Genomic DNA Kit (Invitrogen, ThermoFisher Scientific) following manufacturer’s instructions. Target regions were amplified by PCR (Table S2).

### Micro-electrode array (MEA) recording and data processing

All recordings were performed using the MEA2100-System (Multi Channel Systems) and 60-6wellMEA200/30iR-Ti-rcr chips. The day prior to recording, a medium change was performed. Spontaneous electrophysiological activity of cortical networks was recorded for 20 minutes at 37°C under a constant flow of humidified gas (5% CO₂) using the Multi-Channel Experimenter software. Recording of neuronal networks activity was initiated after 5 minutes of data acquisition. These initial 5 minutes delay was implemented as acclimatisation period to allow for environmental stabilisation of cultures. Additionally, the delay allowed for experimenters to leave the room, thereby eliminating observer-activity that could interfere with the recordings. Before each recording a grounded stainless steel isolation box was placed over the recording device to minimise electromagnetic interference.

Raw electrophysiological signals were continuously acquired at a sampling rate of 10 kHz, ensuring high temporal resolution for spike detection. All further signal processing and analysis were performed offline. Electrophysiological traces were processed using Multi Channel Analyser. In details, Raw signals were first filtered using a Butterworth high pass filter at 200 Hz to eliminate low-frequency noise and improve spike detection accuracy. Spikes were then detected using the spike detector, with thresholds set at ±5 times the standard deviation of the signal amplitude for each individual electrode. Burst detection was performed using the Multi-Channel burst analyser with the following settings: a maximum interspike interval (ISI) of 100 ms to start a burst, maximum ISI of 150 ms to end a burst, and a minimum ISI of 150 ms between bursts. Each burst required a minimum duration of 10 ms and at least 3 spikes to qualify for analysis. Downstream analysis of spike and burst data was performed using custom in-house scripts in R Studio (v4.2.2).

The electrophysiological parameters assessed included: Spike Rate (SR; Hz), representing the overall firing frequency; Percentage of Random Spikes (%RS), defined as the proportion of spikes occurring outside of burst events relative to the total number of spikes; Burst Duration (BD; ms), indicating the temporal length of each burst; Burst Frequency (BF; Hz), reflecting the intra-burst spiking rate; Burst Rate (BR; bursts/min), quantifying the number of bursts per minute; Inter-Burst Interval (IBI; ms), measuring the time between successive bursts; and Spikes in Burst (SiB), denoting the number of spikes per burst event.

### RNA sequencing and data analysis

RNA sequencing was performed at the Translational Genomics (TGX) – Mental Health and Neuroscience Research Institute (MHeNs) Next Generation Sequencing (NGS) CORE facility at Maastricht University. RNA quality control was performed using Qubit fluorometric quantification and capillary electrophoresis on a Bioanalyzer system. Only RNA samples with high integrity (RIN ≥ 6.8) were processed for sequencing. Polyadenylated (Poly-A) mRNA was selectively enriched from total RNA using oligo(dT) magnetic beads, and strand-specific libraries were prepared using the TruSeq Stranded mRNA kit (Illumina). Sequencing was performed on a NovaSeq 6000 S1 flowcell (200 cycles, paired-end reads).

Raw FASTQ files obtained from the Illumina sequencing were processed on a Unix-based system using a custom command-line pipeline. First, quality control was performed using FastQC for each FASTQ file. Alignment was carried out using HISAT2, using the GRCh38 transcriptome-aware reference genome index (grch38_tran.tar.gz). Output SAM files and alignment summary logs were generated for each sample. SAM files were converted to sorted and indexed BAM files using Samtools. For optional visual inspection of alignments, sorted BAM files and their index files were loaded into the Integrative Genomics Viewer (IGV). Overall quality of reads and alignments was summarised using MultiQC, which aggregated reports from FastQC and HISAT2. Gene-level read counts were obtained using featureCounts from the Subread package. The resulting count matrices served as input for differential expression (DE) analysis.

The generated gene expression count matrices were processed and analysed in R using a suite of Bioconductor and CRAN packages, including DESeq2, edgeR, limma, and org.Hs.eg.db for annotation. Count data were normalised using variance-stabilising transformation (VST) as implemented in DESeq2. Sample clustering and batch effects were visualised via principal component analysis (PCA) and sample-to-sample distance heatmaps. Significant genes were defined as those with Benjamini-Hochberg-adjusted p-values (padj) < 0.01 and log_2_FoldChange > 2. Shrinkage of log_2_FoldChange values was applied using the apeglm method. To confirm robustness of DE results, analyses were replicated using both edgeR (quasi-likelihood F-tests) and limma-voom. Gene lists from all three methods were compared via Venn diagrams to assess concordance. The correlation of log_2_ fold-changes (log_2_FC) across methods was evaluated using scatterplot matrices. Concordance of directionality and effect sizes was confirmed through correlation analyses and visual inspection. DE genes accordantly identified by all three methods were further analysed for gene ontology (GO) enrichment analyses for Biological Process (BP), Molecular Function (MF), and Cellular Component (CC) using clusterProfiler. Additional pathway enrichment analyses were performed using Kyoto Encyclopedia of Genes and Genomes (KEGG). KEGG pathway maps with expression overlays were generated using pathview.

## Results

The generation and comprehensive characterisation of the genetically matched lines are detailed in the previous chapter of this thesis.

### Generation of glutamatergic and GABAergic cortical neurons

#### Generation of heterogeneous cortical neurons Obtained via directed differentiation

To generate a heterogeneous population of cortical neurons, hiPSCs were subjected to a developmentally guided differentiation protocol based on dual-SMAD inhibition. Using sequential exposure to media cocktails designed to inhibit SMAD signalling, both WT and *ST3GAL3*^−/−^ iPSCs were differentiated through distinct developmental stages, ultimately yielding mature cortical neuronal networks (Fig. 2A for a schematic representation).

**Figure 1.**
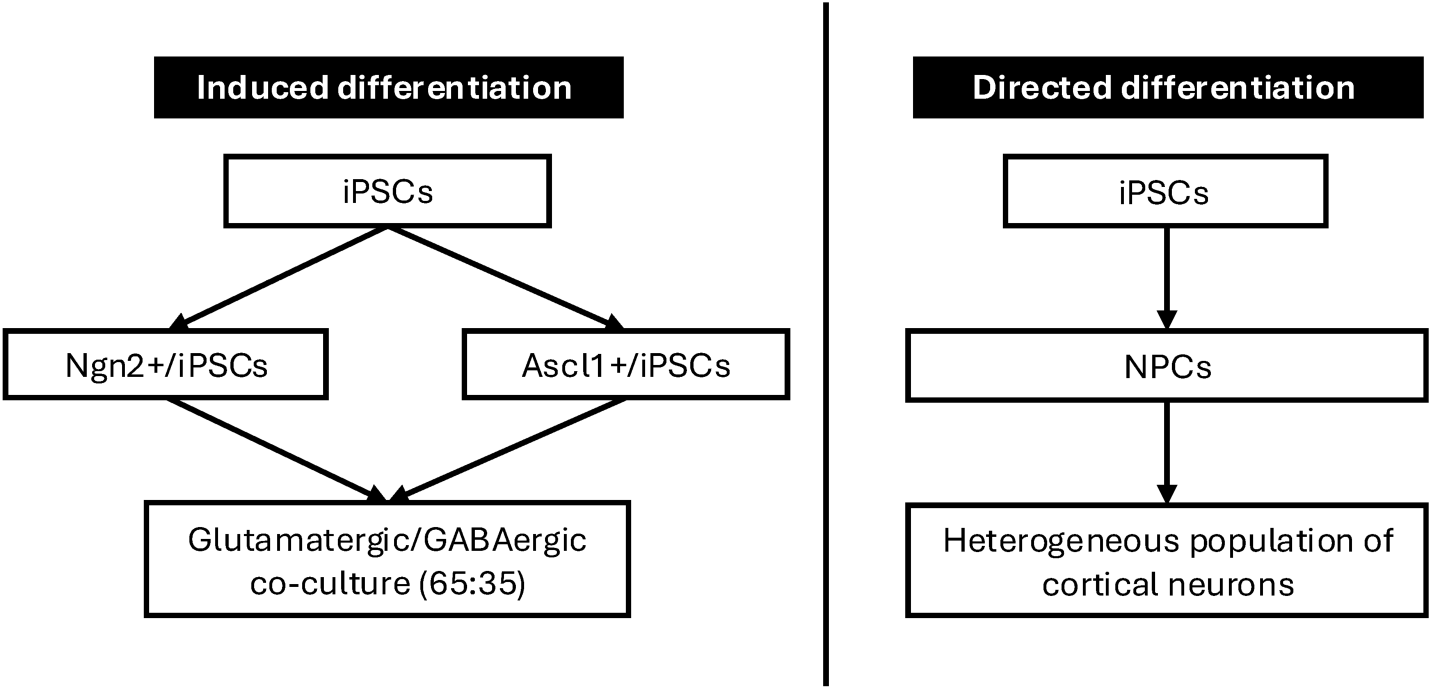
Schematic overview of neuronal differentiation strategies employed in this study. Induced differentiation **(left)** relies on transcription factor–mediated reprogramming of iPSCs into either neurogenin-2 positive (Ngn2⁺) or achaete-scute family BHLH transcription factor 1 positive (Ascl1⁺) lineages, subsequently combined in a defined ratio (65:35). and induced via doxycycline into glutamatergic/GABAergic co-cultures. Directed differentiation **(right)** involves the conversion of induced pluripotent stem cells (iPSCs) into neural progenitor cells (NPCs), followed by maturation into a heterogeneous population of cortical neurons.

**Figure 2.**
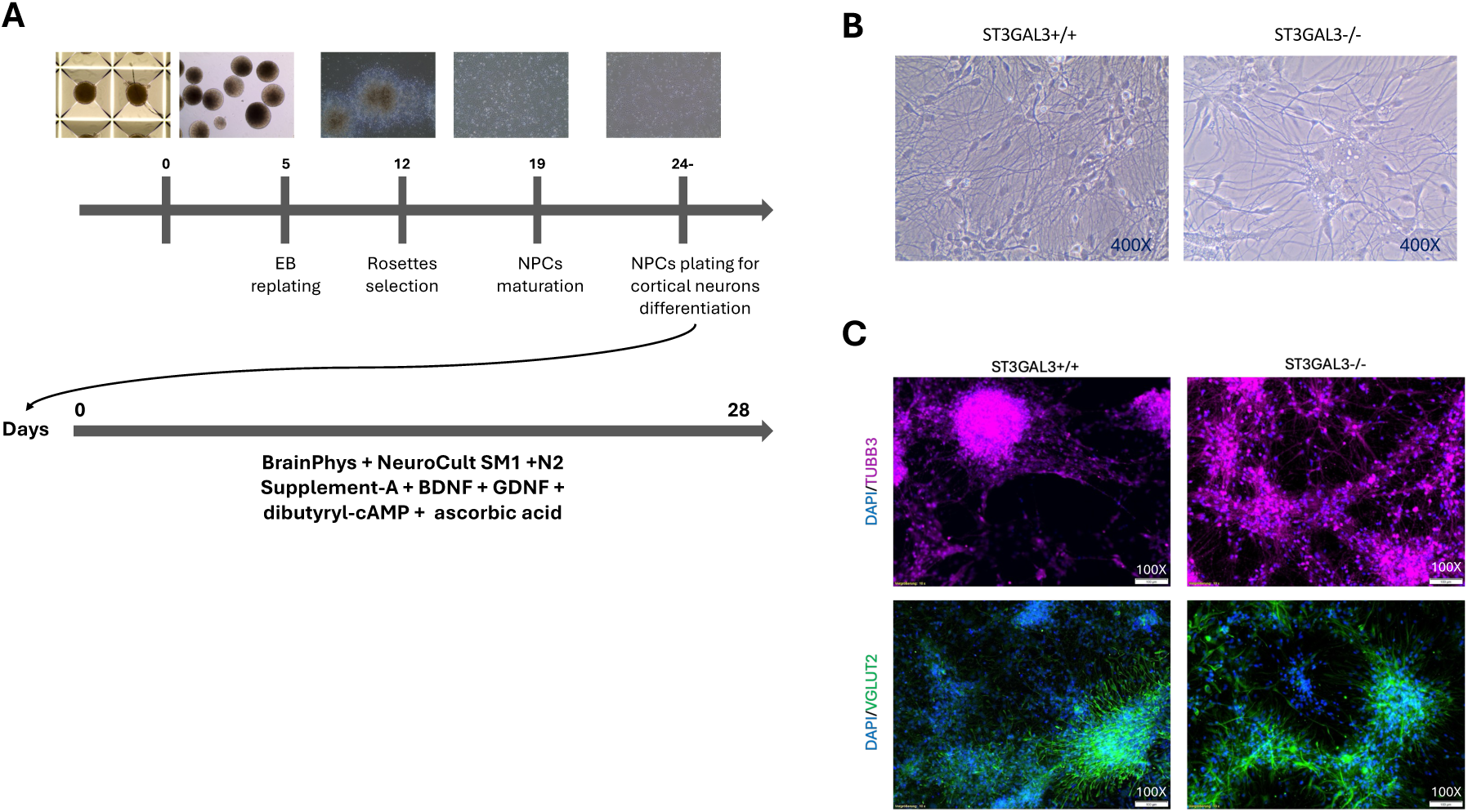
Differentiation of hiPSCs into cortical neurons using a dual-SMAD inhibition protocol (directed differentiation). (**A**) Schematic representation of the stepwise protocol for the directed differentiation of iPSCs into heterogeneous cortical neurons. **(B)** Brightfield images depict key characteristics of cortical neurons. **(C)** ICC analysis confirmed successful neuronal differentiation, as evidenced by expression of the mature neuronal markers βIII-tubulin (TUBB3) and vesicular glutamate transporter 2 (VGLUT2), with complete absence of pluripotency marker octamer-binding transcription factor 3/4 (OCT3/4), indicating elimination of undifferentiated iPSCs. Nuclear stain performed with 4′,6-diamidino-2-phenylindole (DAPI). EB embryoid body. NPC neural precursor cell.

The differentiation process commenced with the formation of EBs, which displayed characteristic spherical morphology and initiated commitment toward the ectodermal lineage (Fig. S4A). Upon plating (Fig. S4B), the EBs flattened and gave rise to radial arrangements of columnar neuroepithelial cells, known as neural rosettes (Fig. S4C). These rosette structures resemble the early neural tube and mark the emergence of patterned neural progenitors. On day 12, rosettes were selected (Fig. S4D) and plated, giving rise to NPC cultures with typical morphology, characterised by elongated bipolar cell bodies and prodromal neurite-like trailing processes (Fig. S4E,F). These features are indicative of radial glia-like neural progenitors undergoing active proliferation and early differentiation. From this stage, NPCs were maintained under neurotrophic conditions for an additional 30 days to facilitate their differentiation into mature cortical neurons (Fig. 2A). We obtained neural networks exhibiting extensive neuritic arborisation (Fig. 2B, S5A,B). Neural-faith was validated via ICC by the expression of mature neuronal markers, including βIII-tubulin (TUBB3) and vesicular glutamate transporter 2 (VGLUT2) (Fig. 2B). Negative OCT3/4 staining confirmed absence of iPSCs in neural cultures (Fig. 2C).

Electrophysiological properties detected via MEA analysis are discussed in a dedicated paragraph.

#### Generation of scalable Glut/GABA co-cultures via induced differentiation

To achieve induced differentiation into glutamatergic and GABAergic neuronal subtypes, iPSCs were transduced with lentiviral particles encoding the rtTA-regulated transcription factors Ngn2 and Ascl1. Owing to the integrating nature of the lentiviral vectors, the transgenes were stably incorporated into the host genome, enabling the generation of iPSC lines with inducible expression of Ngn2 or Ascl1 under the control of a tetracycline-responsive promoter. Upon doxycycline exposure, the reverse tetracycline transactivator (rtTA) binds the promoter and initiates robust transcriptional activation, thereby committing the transduced iPSCs to either a glutamatergic (Ngn2) or GABAergic (Ascl1) neuronal fate. At the initiation of differentiation, iPSCs transduced with Ngn2 and Ascl1 constructs were combined in a 65:35 ratio and cultured for 49 days in doxycycline-supplemented medium to promote the simultaneous induction of glutamatergic and GABAergic neuron fates (Fig. 3A for schematic representation). This specific ratio and differentiation timeline were selected based on the previously validated methodology described by Wang et al (2023) [53].

**Figure 3.**
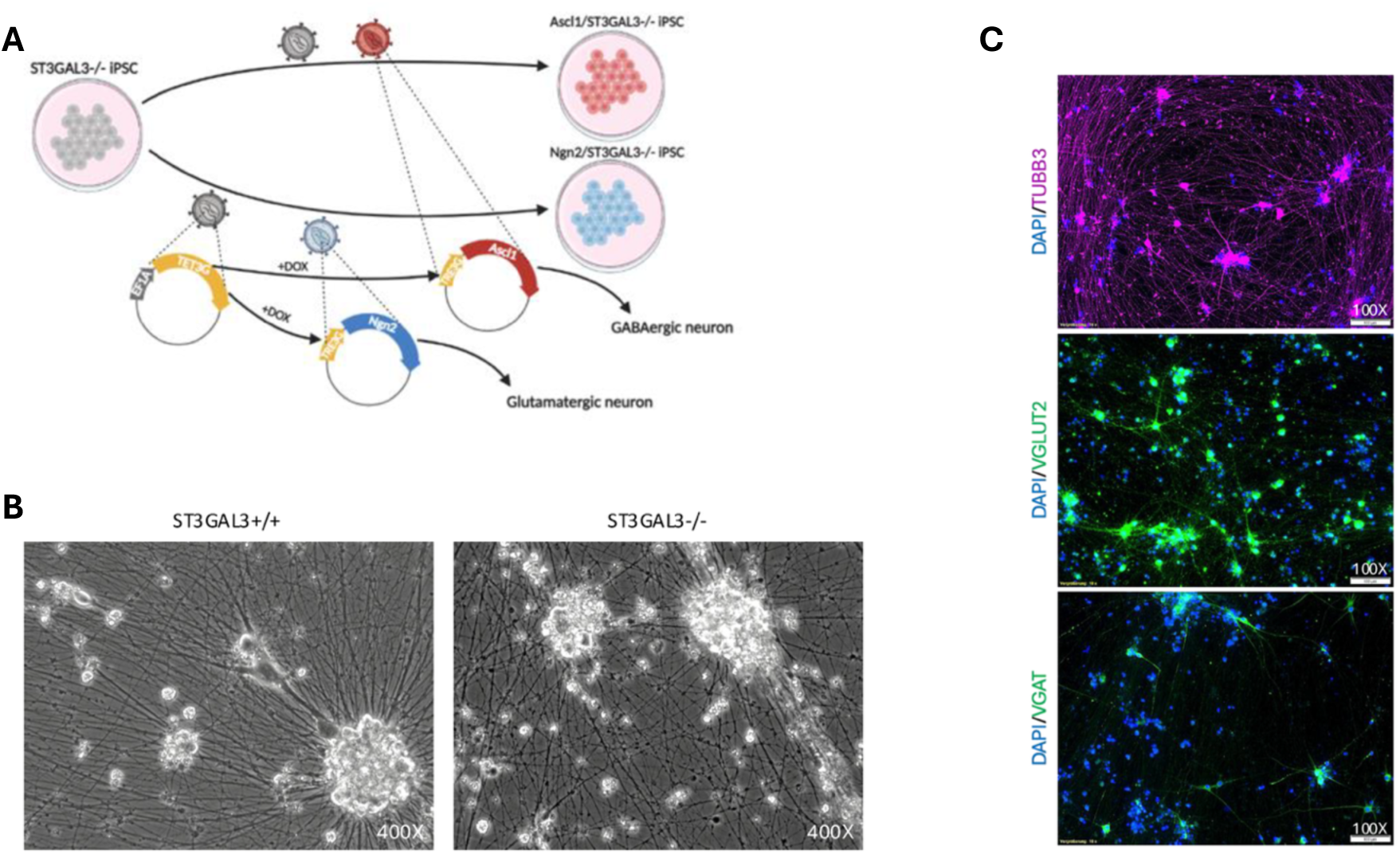
Differentiation of hiPSCs into Glutamatergic/GABAergic co-cultures via a doxycycline-dependent overexpression of Ngn2 and Ascl1 (Induced differentiation). **(A)** Schematic overview of the inducible differentiation system. **(B)** Representative brightfield images depicting key characteristics of cortical neurons. **(C)** ICC analysis confirmed successful neuronal differentiation, as evidenced by expression of the mature neuronal markers βIII-tubulin (TUBB3), vesicular glutamate transporter 2 (VGLUT2), and vesicular GABA transporter (VGAT) Absence of pluripotency marker octamer-binding transcription factor 3/4 (OCT3/4), indicates elimination of undifferentiated iPSCs. Nuclear stain performed with 4′,6-diamidino-2-phenylindole (DAPI). Dox Doxycycline.

Following lentiviral transduction of both *ST3GAL3*^+/+^ and *ST3GAL3*^−/−^ iPSC lines, antibiotic selection with G418 and puromycin was performed to ensure that only cells harbouring stably integrated vectors were retained in the cultures. Successful genomic integration of the lentivirally delivered transgenes was confirmed via PCR and gel electrophoresis: all Ngn2– and Ascl1-transduced iPSCs were positive for the TET3G regulatory element as well as the corresponding Ngn2 or Ascl1 constructs (Fig. S1A-C).

The pluripotent potential of the *ST3GAL3*^−/−^ line was preserved following lentiviral transduction, as evidenced by the consistent expression of protein-level pluripotency markers, OCT3/4, TRA-1-60, and SSEA-4, in *ST3GAL3*^−/−^, Ascl1/*ST3GAL3*^−/−^, and Ngn2/*ST3GAL3*^−/−^ iPSCs (Fig. S2A-C). Furthermore, all lines retained their capacity to differentiate into cell types representing the three germ layers (Fig. S3A-D).

To minimise potential bias arising from variability in transgene expression across ST3GAL3 KO and ST3GAL3 WT lines, multiple clonal iPSC populations were generated. Clones exhibiting markedly divergent levels of Ngn2 or Ascl1 transcript expression following 4 days of doxycycline induction, as assessed by qRT-PCR, were excluded from downstream analyses.

Obtained neurons exhibited extensive neuritic arborisation (Fig. 3B). ICC analysis confirmed the successful neuronal identity and subtype specification of the differentiated cultures. At day *in vitro* (DIV) 49, βIII-tubulin (TUBB3) immunostaining revealed extensive neuronal networks with dense neuritic arborisation, indicative of robust neuronal differentiation and network maturation. Co-labelling with VGLUT2 confirmed the presence of excitatory glutamatergic neurons, while vesicular GABA transporter VGAT staining demonstrated the presence of inhibitory GABAergic neurons (Fig. 3B).

### Transcriptomic characterisation of ST3GAL3-deficient neuronal cultures

To explore the molecular consequences of ST3GAL3 deficiency in human cortical neurons, RNAseq was performed on terminally differentiated cultures derived from both the directed and induced differentiation protocols. Samples from *ST3GAL3*^+/+^ and *ST3GAL3*^−/−^ iPSC lines were harvested at DIV30 (directed) and DIV49 (induced), corresponding to mature stages of network development in each model. For clarity, throughout this analysis the directed differentiation protocol will be referred to as protocol A, and the induced differentiation protocol as protocol B.

### Differential expression analysis

As an initial quality control and exploratory step, PCA was performed on the variance-stabilised RNAseq data. The first two principal components accounted for 81% of the total variance (principal component [PC]1: 61%, PC2: 20%), clearly separating samples according to both differentiation protocol and genotype (Fig. 4). WT and KO cultures clustered distinctly within each protocol, indicating consistent genotype-dependent transcriptional signatures. Furthermore, the large separation between protocols along PC1 reflects substantial global transcriptomic differences attributable to the differentiation strategy, while the separation along PC2 seems to highlight genotype-specific variation within each method. The tight clustering of biological replicates within each group confirms high reproducibility and sample quality for downstream analyses (Fig. 4). A notable observation was made for the protocol A_WT group, where samples appeared to form two distinct sub-clusters. No obvious deviations or technical issues were noted during the differentiation process for any of these replicates, and therefore all were retained for analysis. This increased within-group variability is expected to result in more conservative significance thresholds; however, as will be demonstrated in subsequent sections, the analysis still revealed robust, reproducible, and biologically meaningful differences.

**Figure 4.**
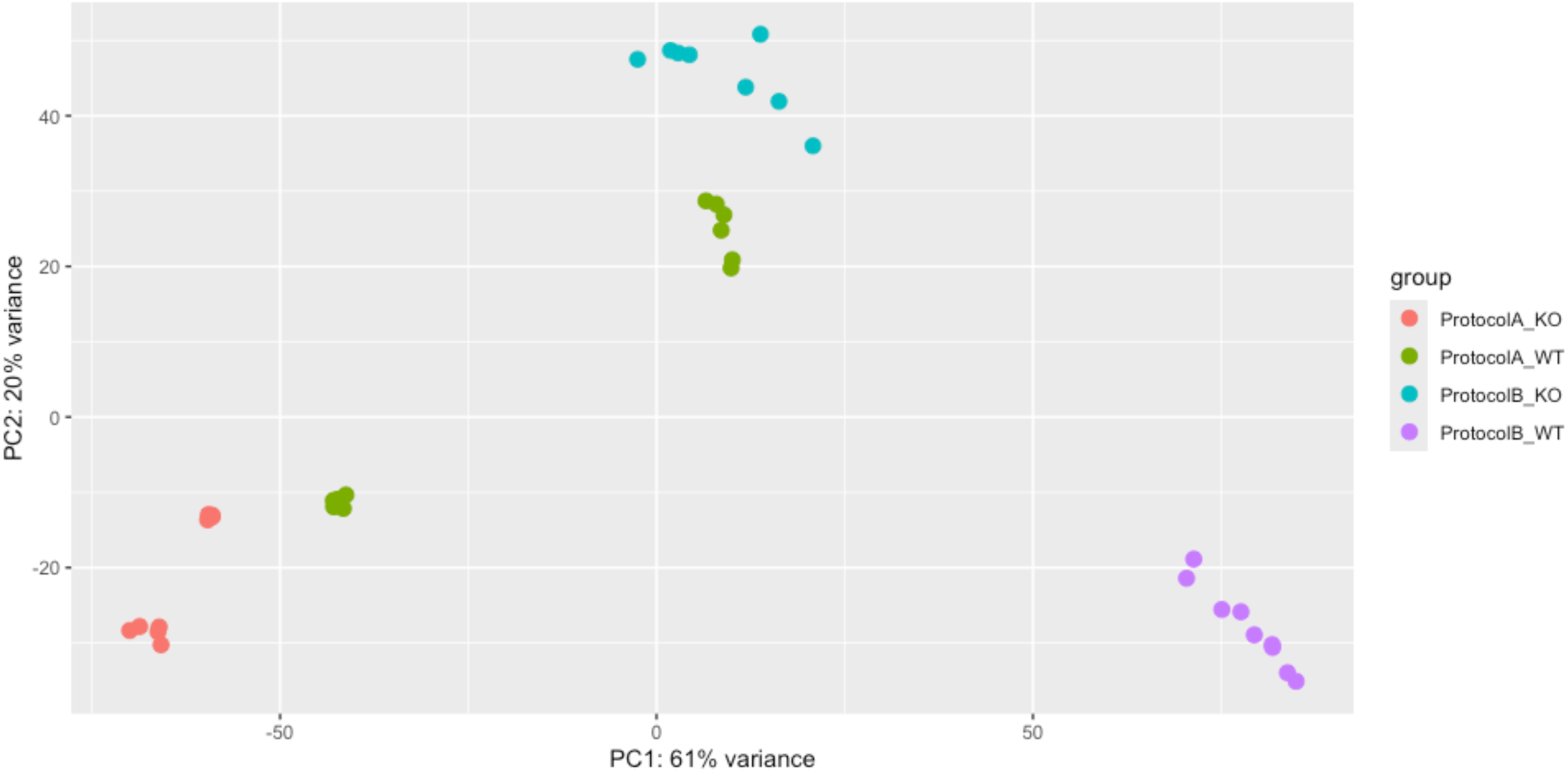
Principal component analysis (PCA) of variance-stabilised RNAseq data. The first two principal components (PC1: 61%, PC2: 20%) explain 81% of the total variance across samples. Clear clustering by differentiation protocol (Protocol A vs. protocol A) is observed along PC1, while PC2 separates samples according to genotype (ST3GAL3 WT vs. ST3GAL3 KO). WT and KO cultures formed distinct clusters within each protocol, reflecting consistent genotype-dependent transcriptional signatures. The tight grouping of biological replicates within each condition demonstrates high reproducibility, with the exception of Protocol A_WT, which showed two sub-clusters despite the absence of technical deviations during differentiation.

Using DESeq2, we identified a total of 2,181 DE genes (DEGs) in *ST3GAL3*^−/−^ vs. *ST3GAL3*^+/+^ in protocol A, applying a significance threshold of padj<0.01 and an absolute log₂FC greater than 2 (Fig. 5A). Similarly, 3,884 differentially expressed genes were detected in protocol B (Fig. 5B). Notably, 921 of these genes were consistently differentially expressed across both protocol A and protocol B (Fig. S7).

**Figure 5.**
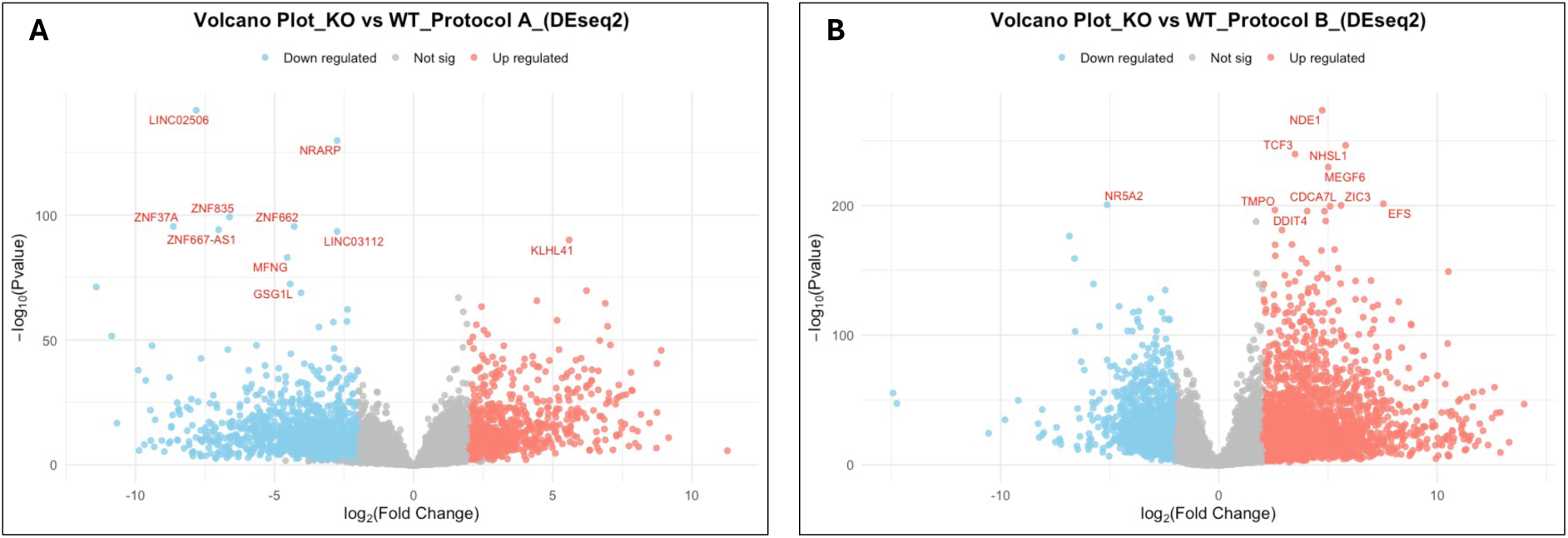
Volcano plots of differentially expressed genes (DEGs) identified via DESq2 in *ST3GAL3*^−/−^ (KO) versus *ST3GAL3*^+/+^ (WT) iPSC-derived neurons. Volcano plots generated with DESeq2 showing the distribution of DEGs for **(A)** protocol A and **(B)** protocol B. The x-axis represents log_2_FC, and the y-axis represents –log₁₀ adjusted p-value. Genes significantly downregulated in KO relative to WT are shown in blue, whereas upregulated genes are shown in red; non-significant genes are indicated in grey. The top 10 DEGs per protocol, ranked by adjusted p-value, are labelled. Significance was determined with threshold of Padj < 0.01 and an absolute log_2_FC > 2.

Given the high number of significantly DEGs, we sought to enhance the robustness of our findings by applying two additional, widely used RNAseq differential expression analysis methods: limma-voom and edgeR (Fig. 6A-D). In support of our findings, the three methods showed a high degree of overlap in the differentially expressed genes they identified, with 1,023 genes common to all three analyses in protocol A and 3,648 in protocol B (Fig. S8A,B). All three methods also demonstrated a strong correlation in their estimated log₂FC (Fig. S9A,B). For downstream analyses, we therefore focused on the subset of genes consistently detected by all three statistical methods for each protocol.

**Figure 6.**
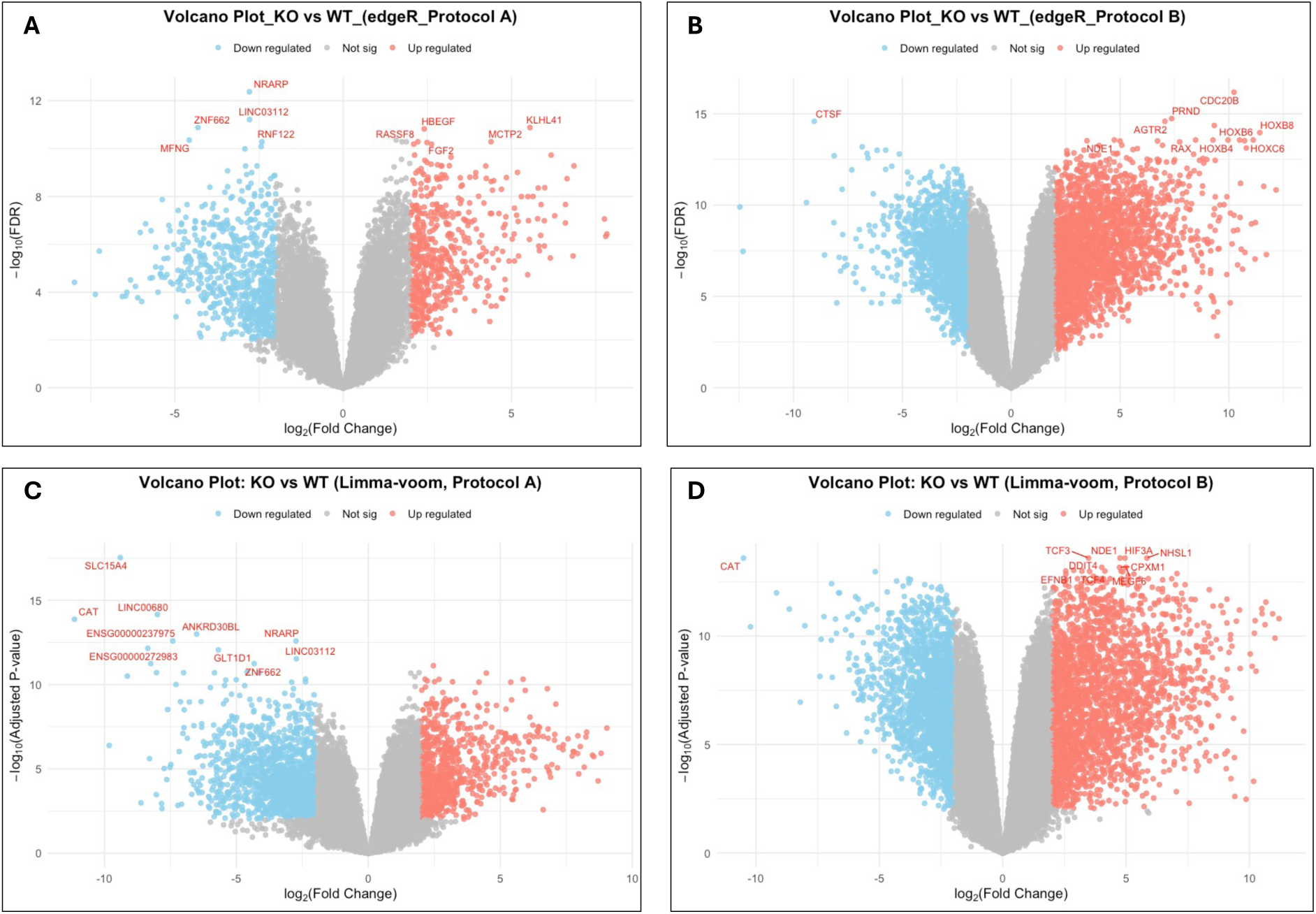
Volcano plots of differentially expressed genes (DEGs) in *ST3GAL3*^−/−^ (KO) versus *ST3GAL3*^+/+^ (WT) iPSC-derived neurons identified with edgeR and limma-voom. Volcano plots generated using **(A, B)** edgeR and **(C, D)** limma-voom showing the distribution of DEGs for protocol A **(A, C)** and protocol B **(B, D)**. The x-axis represents log_2_FC, and the y-axis represents –log₁₀ adjusted p-value (FDR for edgeR, padj for limma-voom). Genes significantly downregulated in KO relative to WT are shown in blue, whereas upregulated genes are shown in red; non-significant genes are indicated in grey. The top 10 DEGs per protocol, ranked by adjusted p-value, are labelled. Significance was determined with threshold of adjusted P< 0.01 and an absolute log_2_FC > 2

### Gene ontology analysis

We conducted GO enrichment analyses for BP, CC, and MF using clusterProfiler, enabling us to examine functional changes from broad biological categories down to more specific molecular details. The BP analysis on protocol A, revealed that many differentially expressed genes were associated with pathways directly relevant to our hypothesis, including *vesicle-mediated transport at the synapse*, *learning or memory*, *cognition*, and *neurotransmitter secretion* (Fig. 7A, top 20 pathways). In addition to these pathways, several muscle tissue-related terms were also enriched. While not biologically relevant to our study, this phenomenon is a recognised occurrence in RNAseq analyses of iPSC-derived neurons, reflecting the substantial overlap between genes involved in neurotransmission and those associated with muscle function.

**Figure 7.**
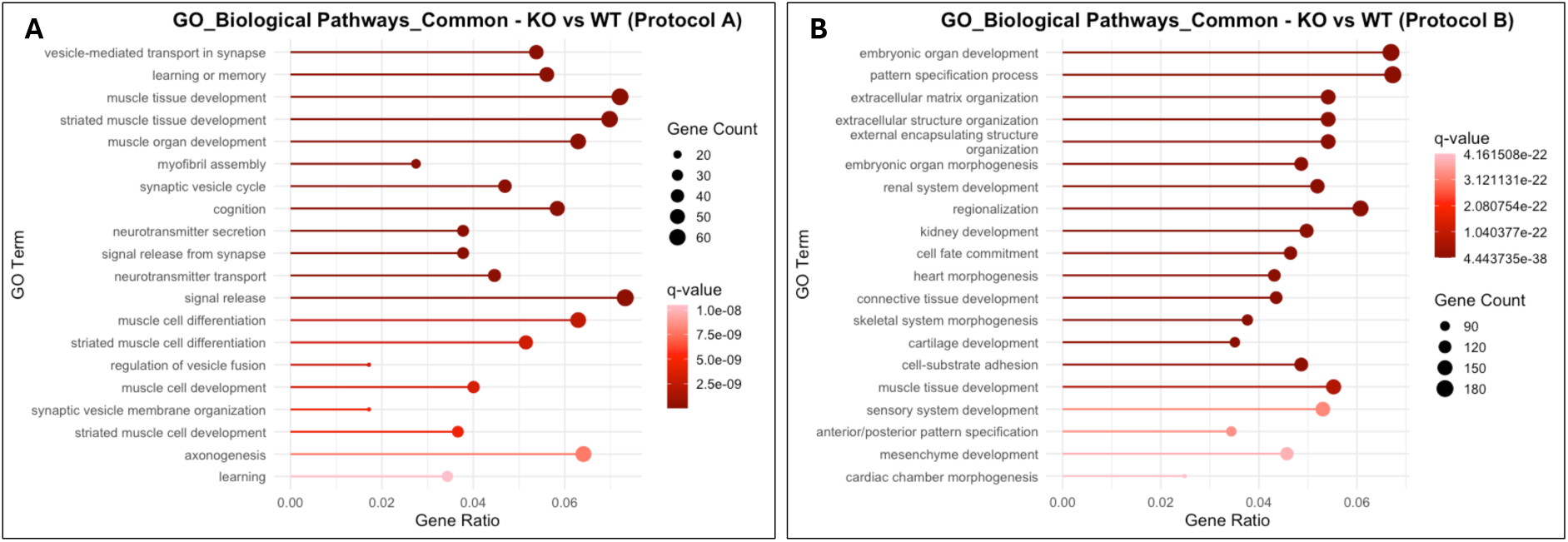
BP enrichment analysis of DEGs in *ST3GAL3*^−/−^ (KO) versus *ST3GAL3*^+/+^ (WT) neurons. Bubble plots showing the top 20 enriched GO biological processes for DEGs identified with DESeq2, limma-voom, and edgeR in **(A)** Protocol A and **(B)** protocol B. Pathways are ranked according to enrichment significance (q-value). The x-axis represents the gene ratio (proportion of DEGs annotated to the GO term), while bubble size corresponds to the number of DEGs and bubble colour reflects the q-value. GO gene ontology.

Interestingly, the vast majority of genes within the identified relevant pathways were significantly downregulated in KO lines compared with WT (Fig. 8). Notably, nearly all of the relevant pathways identified were associated with synapse function, with the exception of *axonogenesis*, which also contained numerous genes that were downregulated (Fig. 8).

**Figure 8.**
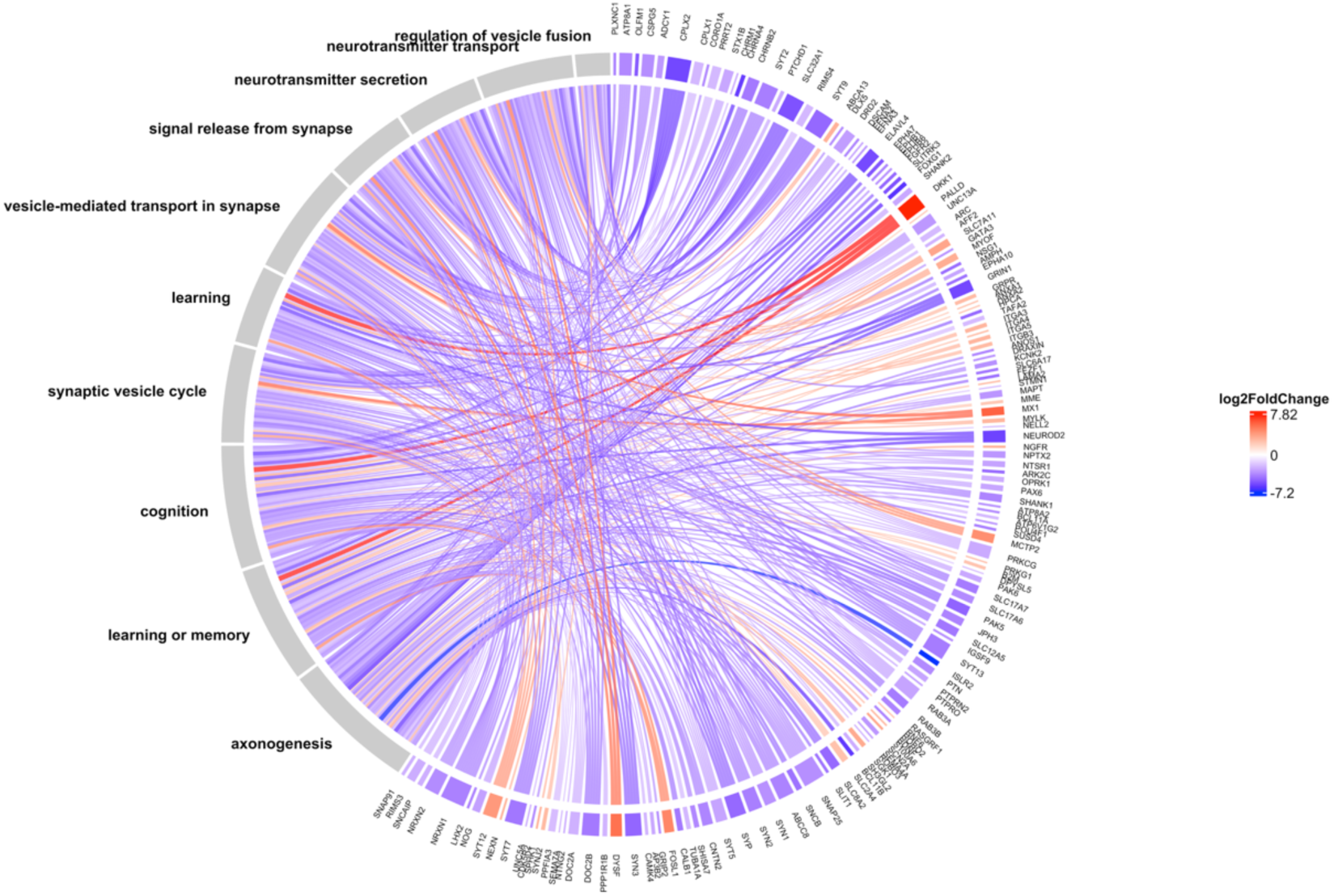
BP Chord diagram linking selected GO biological processes to DEGs in *ST3GAL3*^−/−^ (KO) neurons derived from protocol. **A**. Curated set of ten relevant GO Biological Process terms—chosen for their pertinence to the study design and q-values—are shown connected to DEGs identified from DESeq2, limma-voom and edgeR from the comparison of KO versus WT neurons obtained via directed differentiation (protocol A). Chords denote gene–term associations; outer labels list GO terms (left) and genes (right). Colours encode gene log_2_FC (red = upregulated, blue = downregulated in KO). Log_2_FC values were derived from DESeq2.

Protocol B exhibited a markedly greater variability in BPs affected by ST3GAL3 KO (Fig. 7B), reflecting the substantially higher number of differentially expressed genes identified in this protocol compared with protocol A. Interestingly, many BPs identified for protocol B were associated with extracellular matrix organisation or cell-substrate adhesion (Fg. 7B). Given the established role of sialylated epitopes in mediating extracellular matrix interactions, these findings are particularly noteworthy in the context of ST3GAL3 function.

Importantly, many BPs were shared between the two protocols, including *neurotransmitter transport*, *cognition*, and *vesicle-mediated transport at the synapse*. However, in protocol B, these pathways displayed greater variability in the direction of change compared to protocol A, with a substantial proportion of genes upregulated in ST3GAL3 KO-derived neurons relative to WT (Fig. 9).

**Figure 9.**
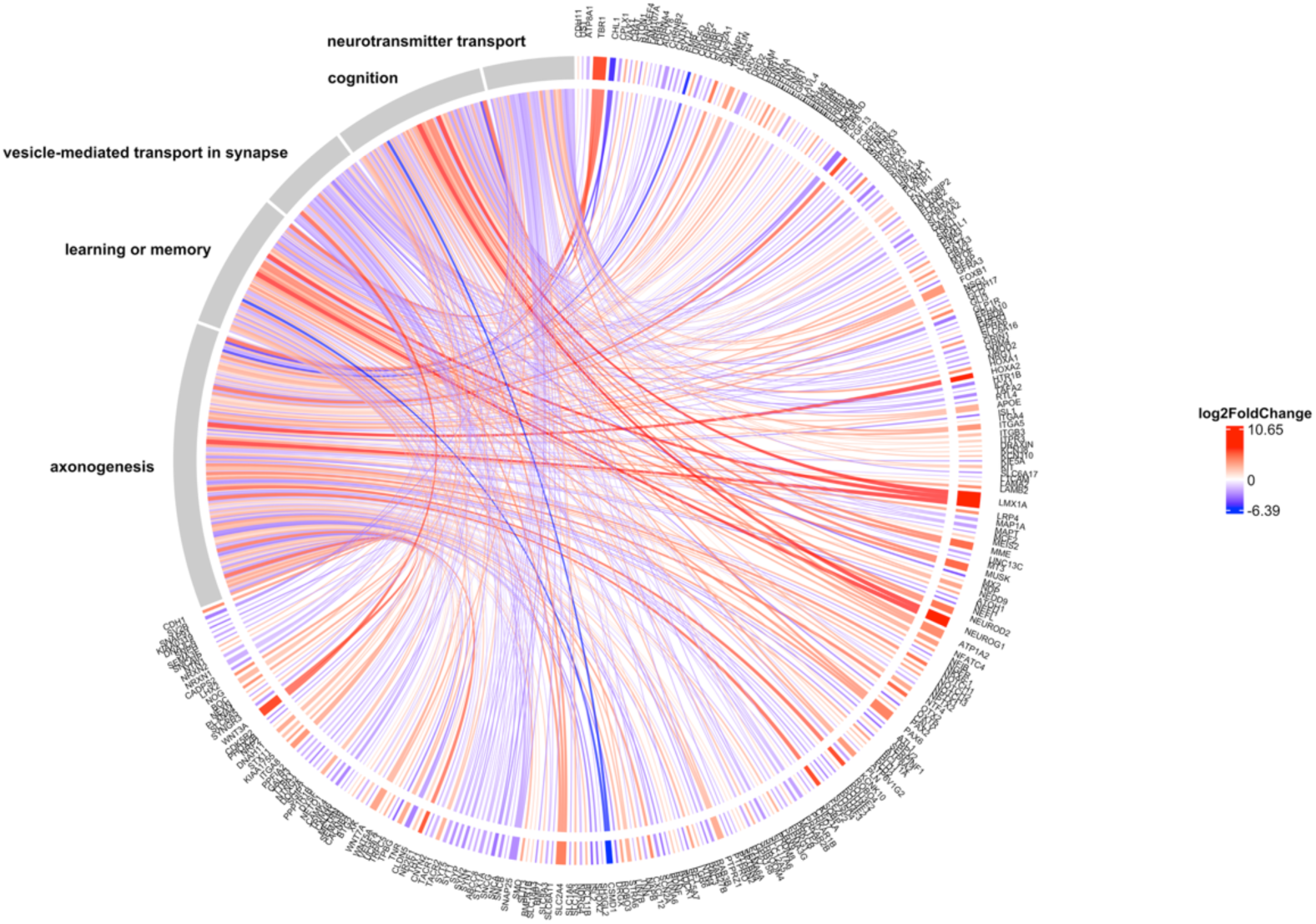
BP Chord diagram linking selected GO biological processes to DEGs in *ST3GAL3*^−/−^ (KO) neurons derived from protocol B. Curated set of 5 relevant GO Biological Process terms – chosen for their pertinence to the study design and q-values – are shown connected to DEGs identified from DESeq2, limma-voom and edgeR from the comparison of KO versus WT neurons obtained via induced differentiation (protocol B). Chords denote gene–term associations; outer labels list GO terms (left) and genes (right). Colours encode gene log_2_FC (red = upregulated, blue = downregulated in KO). log_2_FC values were derived from DESeq2.

The CC analysis for protocol A provided strong support for our hypothesis regarding the pivotal role of ST3GAL3 in synaptic function. Among the top 20 most significantly enriched CC terms were: *synaptic membrane*, *postsynaptic membrane*, *synaptic vesicle membrane*, *exocytic vesicle membrane*, *synaptic vesicle*, *neuron projection terminus*, *postsynaptic specialisation*, *neuron to neuron synapse*, *pre-synaptic membrane*, and *post-synaptic specialisation membrane* (Fig. 10A). *Distal axon* and *axon terminus* were also strongly enriched, suggesting an unknown role of ST3GAL3 in axon formation (Fig. 10A). As expected from BP analysis, most of this relevant CC terms were characterised by a massive downregulation of their genes in ST3GAL3 KO-derived cultures (Fig. 11).

**Figure 10.**
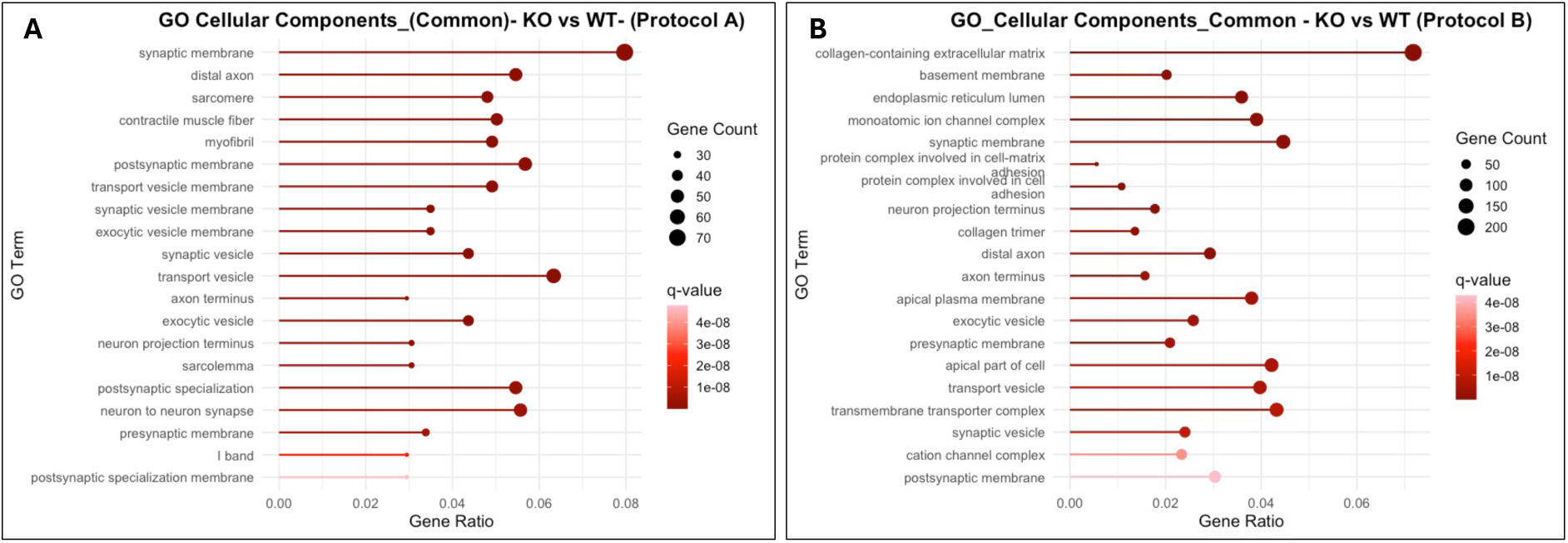
CC enrichment analysis of DEGs in *ST3GAL3*^−/−^ (KO) versus ST3GAL3^+/+^ (WT) neurons. Bubble plots showing the top 20 enriched GO cellular components for DEGs identified with DESeq2, limma-voom, and edgeR in **(A)** Protocol A and **(B)** protocol A. Pathways are ranked according to enrichment significance (q-value). The x-axis represents the gene ratio (proportion of DEGs annotated to the GO term), while bubble size corresponds to the number of DEGs and bubble colour reflects the q-value. GO gene ontology.

**Figure 11.**
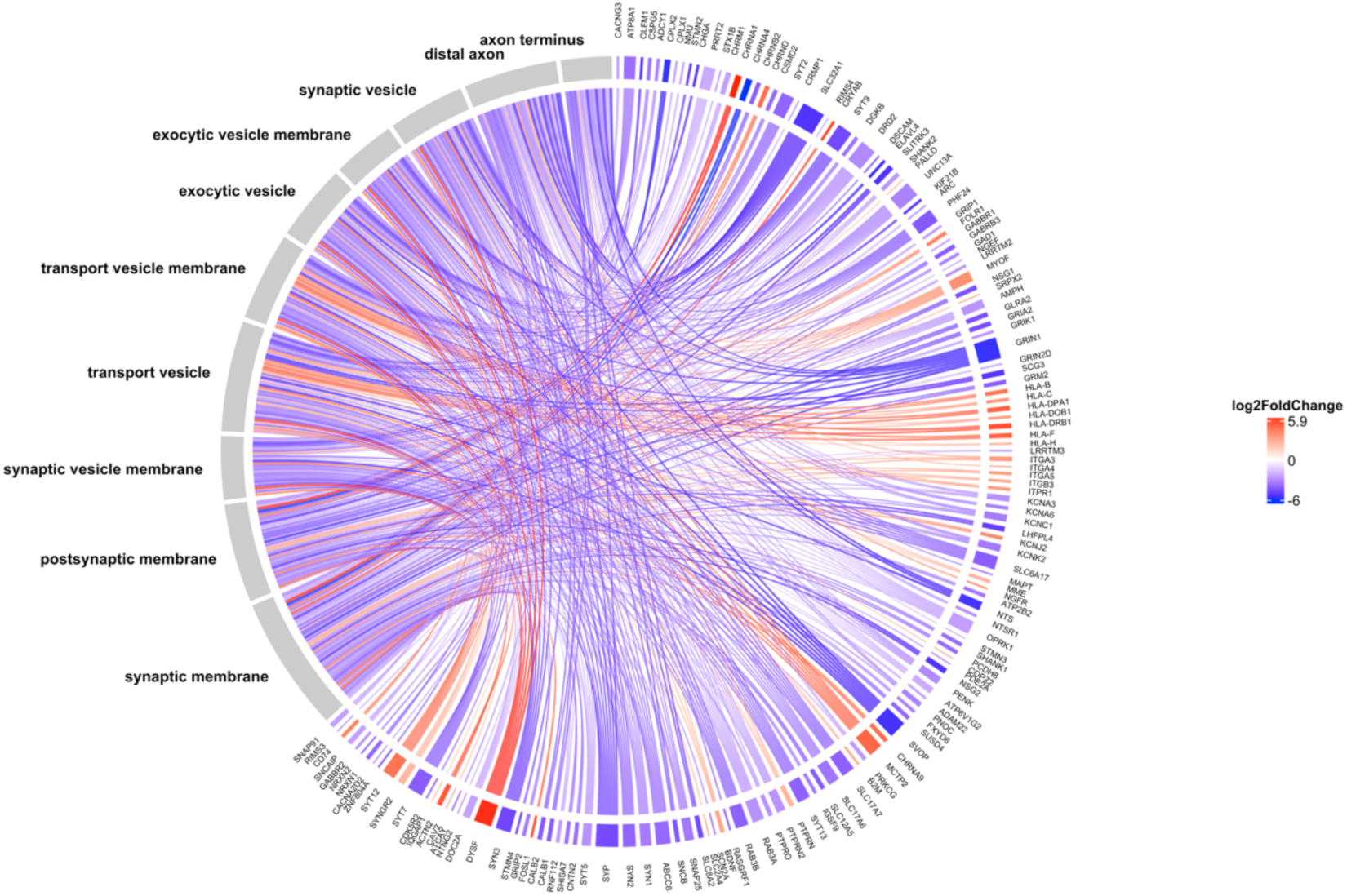
CC Chord diagram linking selected GO cellular components to DEGs in *ST3GAL3^−/−^* (KO) neurons derived from protocol A. Curated set of ten relevant GO cellular component terms—chosen for their pertinence to the study design and q-values—are shown connected to DEGs identified from DESeq2, limma-voom and edgeR from the comparison of KO versus WT neurons obtained via directed differentiation (protocol A). Chords denote gene–term associations; outer labels list GO terms (left) and genes (right). Colours encode gene log_2_FC (red = upregulated, blue = downregulated in KO). Log_2_FC values were derived from DESeq2.

The CC analysis for protocol B further reinforced the synaptic findings and supported our central hypothesis. Among the top 20 most significantly enriched CCs were key synapse-associated terms, including *synaptic membrane*, *neuron projection terminus*, *exocytic vesicle*, *presynaptic membrane*, *transport vesicle*, *synaptic vesicle*, and *postsynaptic membrane* (Fig. 10B). This pattern underscores the pervasive involvement of ST3GAL3 in molecular pathways critical for synaptic architecture and function. Consistent with the BP findings, several of the top 20 CC terms were related to cell–matrix and cell–cell interactions. These included the *collagen-containing extracellular matrix*, *protein complexes involved in cell–matrix adhesion*, *protein complexes in cell adhesion*, and the *collagen trimer*. Given the established mechanistic role of sialylated epitopes expressed on cell surfaces in interacting with the external environment, it is not surprising to observe these terms alongside those related to synapse formation. Notably, *distal axon* and *axon terminus* CC terms were also strongly enriched in the protocol B analysis, further supporting a strong association between *ST3GAL3* and axon-related gene networks (Fig. 10B). Notably, neural related gene-networks, such as the *synaptic membrane*, *neuron projection terminus*, and *distal axon* terms exhibited strong downregulation, mirroring the patterns observed in protocol A, whereas the protein complexes involved in cell–matrix adhesion and cell–cell adhesion were markedly upregulated (Fig. 12).

**Figure 12.**
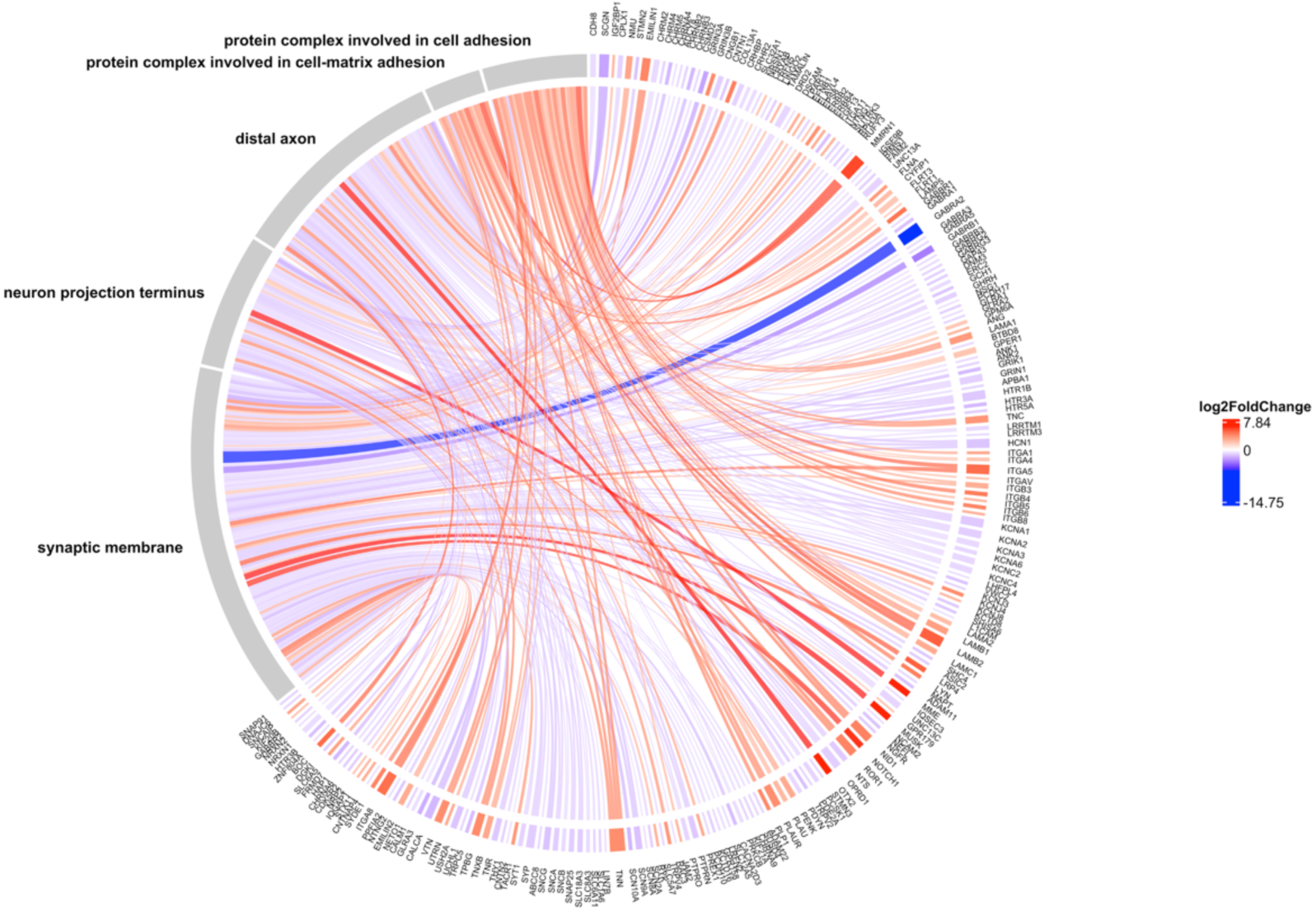
CC Chord diagram linking selected GO cellular components to DEGs in *ST3GAL3^−/−^* (KO) neurons derived from protocol B. Curated set of ten relevant GO cellular component terms—chosen for their pertinence to the study design and q-values—are shown connected to DEGs identified from DESeq2, limma-voom and edgeR from the comparison of KO versus WT neurons obtained via directed differentiation (protocol B). Chords denote gene–term associations; outer labels list GO terms (left) and genes (right). Colours encode gene log_2_FC (red = upregulated, blue = downregulated in KO). Log_2_FC values were derived from DESeq2.

The MF analysis for protocol A revealed strong enrichment across two principal domains: ion transmembrane transport, and protein/lipid-binding complexes (Figure. 13A). Ion transport–related terms featured prominently, including *gated channel activity*, *voltage-gated monoatomic cation channel activity*, *calcium ion transmembrane transporter activity*, and *metal ion transmembrane transporter activity*. These terms point to a significant role for ST3GAL3 in calcium-dependent ion transport across membranes, a process critically important for synaptic function. Protein-binding related terms are: *calmodulin binding*, *glycosaminoglycan binding*, *calcium-dependent phospholipid binding*, *heparin binding*, *integrin binding*, *peptide antigen binding*, and *SNARE binding*. These latter identified terms clearly link transcriptomic alterations to synaptic vesicle fusion, neurotransmitter release, and extracellular interactions. Together, these results support a role for ST3GAL3 in regulating neuronal excitability and synaptic transmission, while the enrichment for extracellular matrix-related functions aligns with the known role of sialylated epitopes in cell–matrix and cell-cell communication.

**Figure 13.**
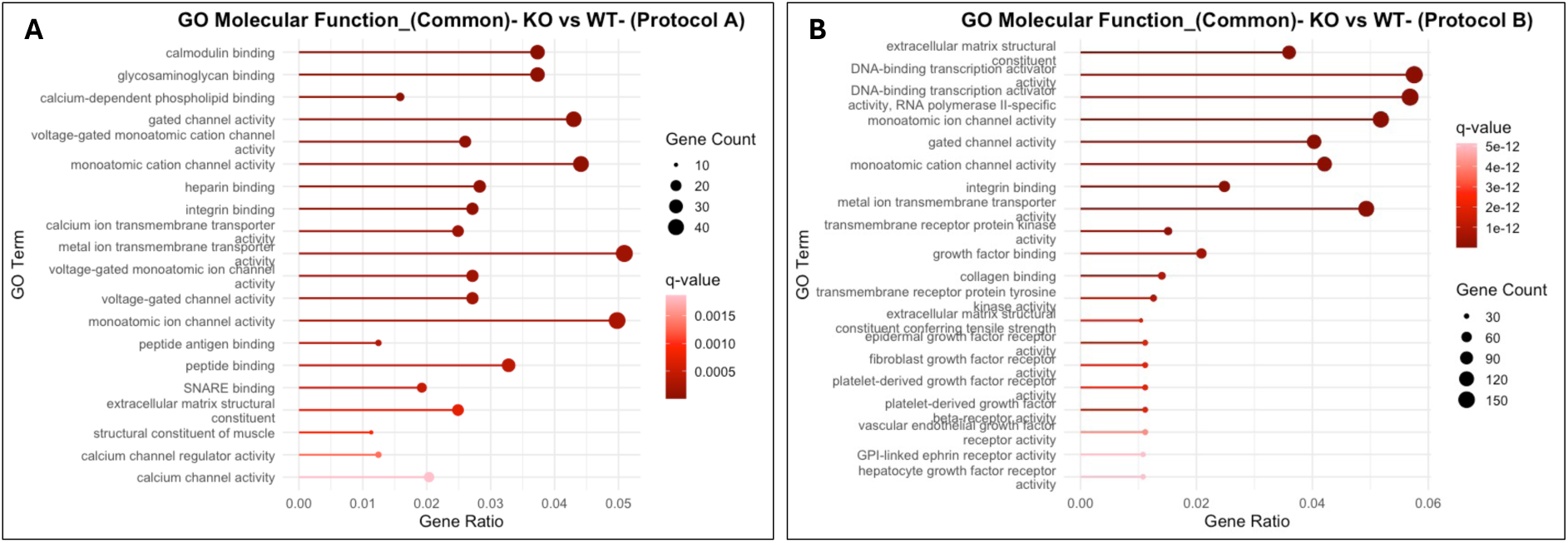
MF enrichment analysis of DEGs in *ST3GAL3^−/−^* (KO) versus *ST3GAL3^+/+^* (WT) neurons. Bubble plots showing the top 20 enriched GO molecular functions for DEGs identified with DESeq2, limma-voom, and edgeR in **(A)** protocol A and **(B)** protocol A. Pathways are ranked according to enrichment significance (q-value). The x-axis represents the gene ratio (proportion of DEGs annotated to the GO term), while bubble size corresponds to the number of DEGs and bubble colour reflects the q-value. GO gene ontology.

The MF analysis for protocol B strongly corroborated the findings from protocol A. Among the top 10 terms ranked by q-value were *extracellular matrix structural constituent*, *monoatomic ion channel activity*, *gated channel activity*, *monoatomic cation channel activity*, *integrin binding*, and *metal ion transmembrane transport activity* (Fig. 13B). These results underscore protocol-independent effects of ST3GAL3 KO on these molecular functions, highlighting consistent alterations in the presented pathways. Two pathways not identified in protocol A displayed strong fluctuations in transcript expression in protocol B: *DNA-binding transcription activator activity* and *RNA polymerase II–specific DNA-binding transcription activator activity*.

### KEGG pathway enrichment analysis

To corroborate our main hypothesis and the patterns identified through differential expression analysis and GO enrichment, we mapped and visualised the differentially expressed genes onto selected molecular pathways relevant to the excitatory/inhibitory balance in neural networks. This was achieved using KEGG pathway enrichment analysis in combination with Pathview. KEGG terms selected were *glutamatergic synapse* (hsa04724), *GABAergic synapse* (hsa04727), *long-term depression* (hsa04730), *long-term potentiation* (hsa04720), *neuroactive ligand signalling* (hsa04082), *synaptic vesicle cycle* (hsa04721), *axon guidance* (hsa04360), and *cell adhesion molecules* (hsa04514). From the set of cell adhesion molecule–related genes, we selected only those associated with neurons, referring to this subset as *neuron-specific cell adhesion* genes. Log2FC were extracted from DESeq2, as this method showed the highest correlation with the other two differential expression (DE) approaches used (Fig S9A,B). We then highlighted the genes identified as differentially expressed by all three methods to ensure the statistical robustness. Overall, this analysis enabled the identification of specific gene transcripts with high statistical and biological relevance pertinent to the aims of our study.

In view of *neuroactive ligand signalling*, we observed an overall downregulation of this gene set in ST3GAL3 KO-derived cortical neurons (Fig. S11 and S12). No genes were identified as significantly upregulated across all three statistical methods in protocol A, whereas protocol B contained only a small subset of upregulated genes (Fig. S11A,B). The significantly downregulated genes showed a high degree of overlap between the two protocols, with most of those identified in protocol A also being significantly downregulated in protocol B (Fig. S12A,B). The marked overlap in gene expression changes observed between the two protocols suggests that the effects of ST3GAL3 KO on this subset of genes are independent of the differentiation method employed. The greater proportion of significantly DEGs in protocol B (Fig. S11B) is consistent with our earlier DE analysis, which revealed a more pronounced segregation of KO and WT transcriptomic signatures in this protocol.

Turning to the more specific genes associated with glutamatergic and GABAergic synapses, we observed, in both protocol A and protocol B, a pronounced downregulation of these genes in ST3GAL3 KO–derived cultures (Fig. 14 and 15).

**Figure 14.**
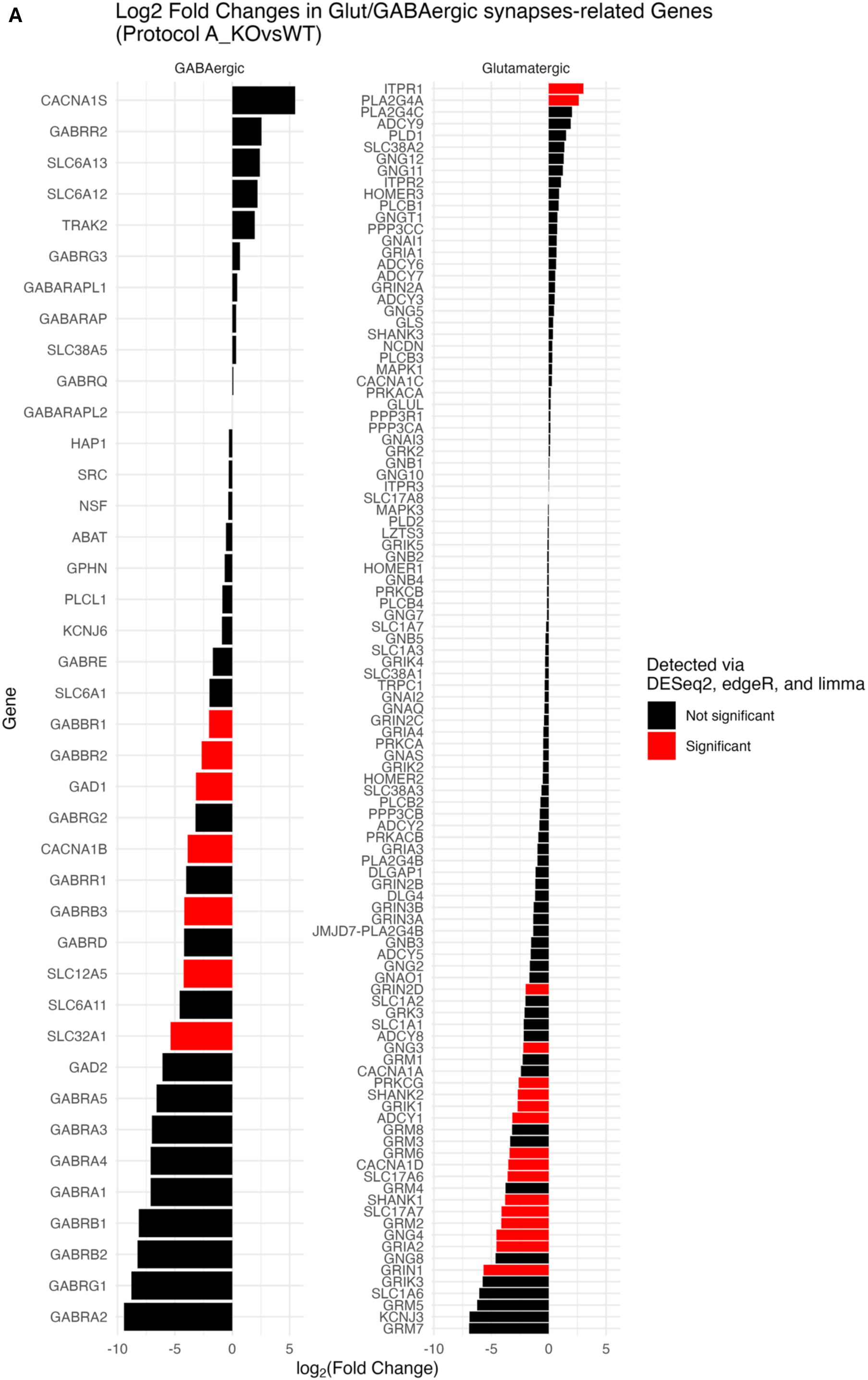
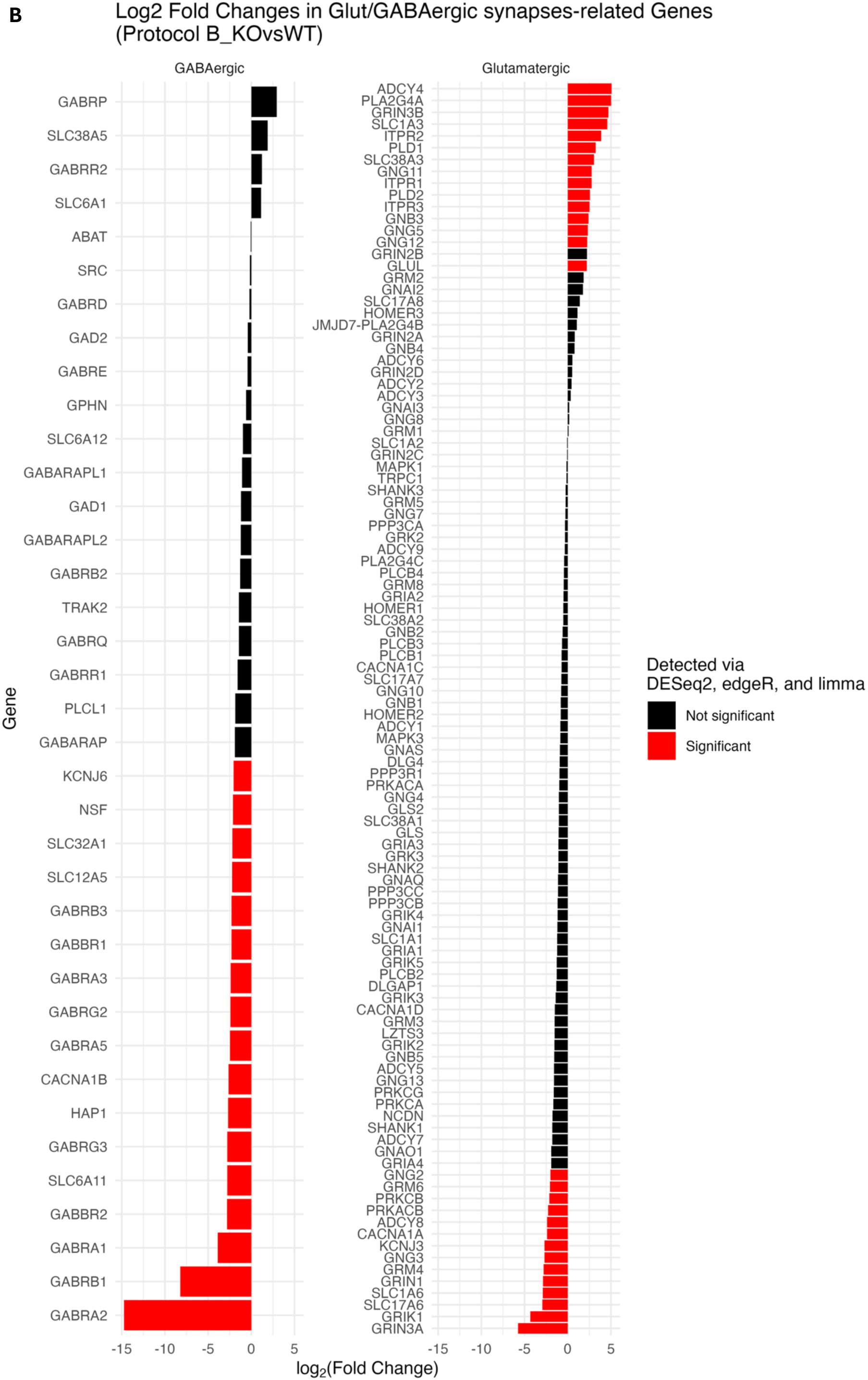
log_2_FC of Glutamatergic synapse (hsa04724) and GABAergic synapse (hsa04727) genes in *ST3GAL3−/−* (KO) versus ST*3GAL3^+/+^* (WT) cultures. (**A**) Results from obtained via directed differentiation (protocol A). **(B)** Results from obtained via induced differentiation (protocol B) Genes were considered significant when consistently detected by DESeq2, edgeR, and limma. Significant differential expression is highlighted in red, while black bars denote non-significant genes. Log_2_FC values derived from DESeq2.

**Figure 15.**
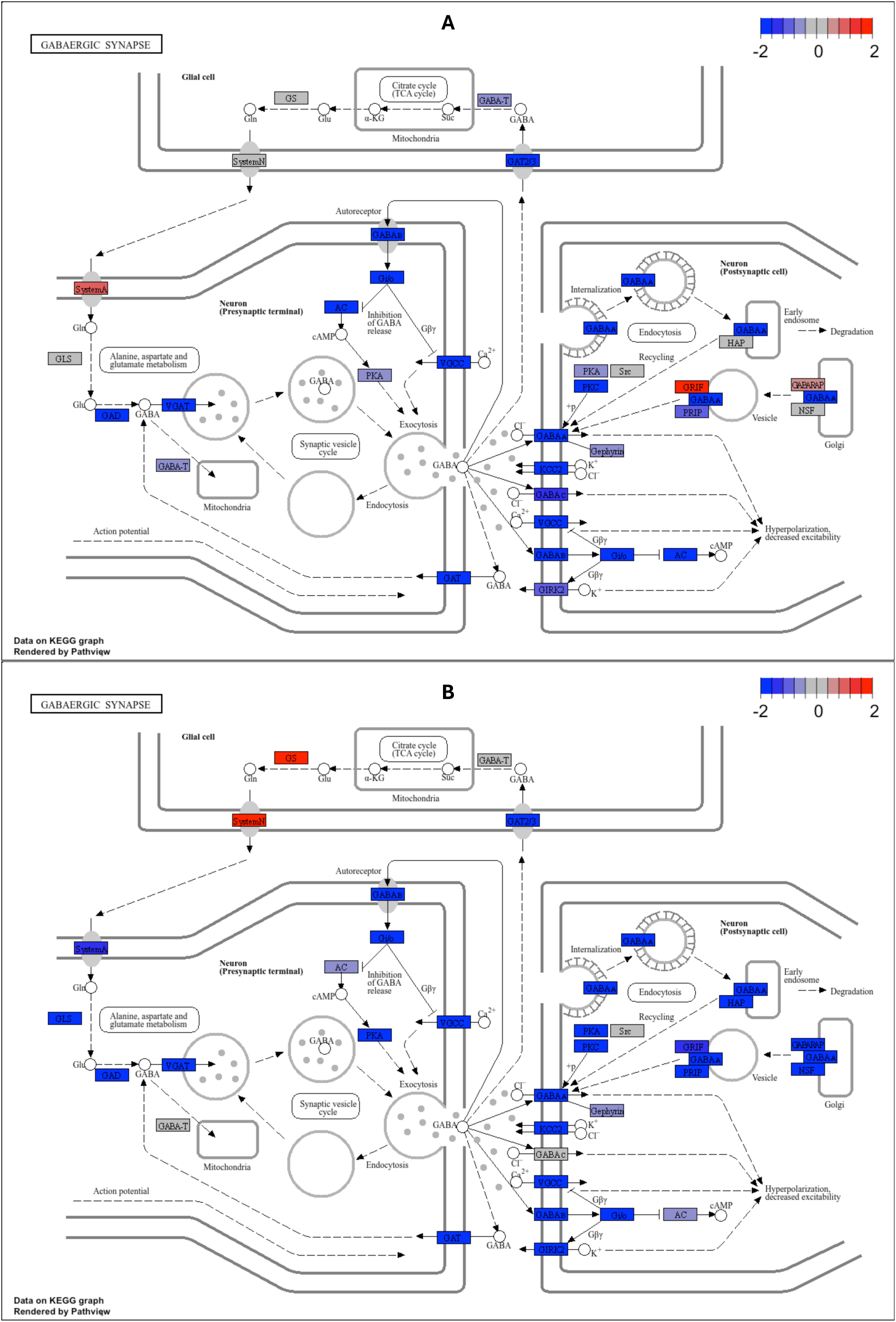
KEGG pathway representation of GABAergic synapse (hsa04727) genes. Log_2_FC between *ST3GAL3^−/−^* (KO) and *ST3GAL3^+/+^* (WT) neurons differentiated via **(A)** protocol A or **(B)** protocol B were mapped onto their corresponding proteins in the KEGG *GABAergic synapse* pathway using Pathview. Boxes represent proteins encoded by the DEGs identified via DEseq2, coloured according to log_2_FC (red = upregulated, blue = downregulated in KO relative to WT).

The analysis of genes associated with *GABAergic synapses* yielded notable findings. None were significantly upregulated in ST3GAL3 KO-derived neural cultures in either protocol, whereas a substantial number showed significant downregulation (Fig. 14A,B). Additionally, with the exception of *GAD1*, which was not identified as significant in protocol B, genes found to be significantly downregulated by all three statistical methods in protocol A showed complete (100%) overlap with those in protocol B. These included *SLC32A1, SLC12A5, GABRB3, CACNA1B, GABBR2,* and *GABBR1*, indicating a strong association between ST3GAL3 function and the regulation of this gene set (Fig. 15A,B).

It is also noteworthy that, in both protocol A and protocol B, *GABRA2* exhibited the largest log_2_FC (Fig. 14). Although it was not identified as significant by all three statistical methods for protocol A, DESeq2 analysis revealed strong significance (padj = 1.91 × 10⁻¹⁰) with a log_2_FC of –9.45, and even greater significance (padj = 4.22 × 10⁻⁴⁸) with a log_2_FC of –14.75 in protocol B. Overall, we observed a consistent downregulation of GABA receptor subunits in ST3GAL3 KO-derived neural cultures, regardless of the differentiation protocol used (Fig. 15).

Genes associated with *glutamatergic synapses* showed a broadly similar distribution; however, a subset was upregulated in ST3GAL3 KO neurons, particularly in protocol B (Fig. 14 and 16). Among these, *ITPR1* and *PLA2G4A* were found to be common to both protocol A and protocol B (Fig. 14). A high degree of overlap was observed in the significantly downregulated genes between the two protocols. *GNG3, GRIK1, GRM6, SLC17A6, GRM2,* and *GRIN1* were identified as significant by all three statistical methods in both protocols (Fig. 14).

**Figure 16.**
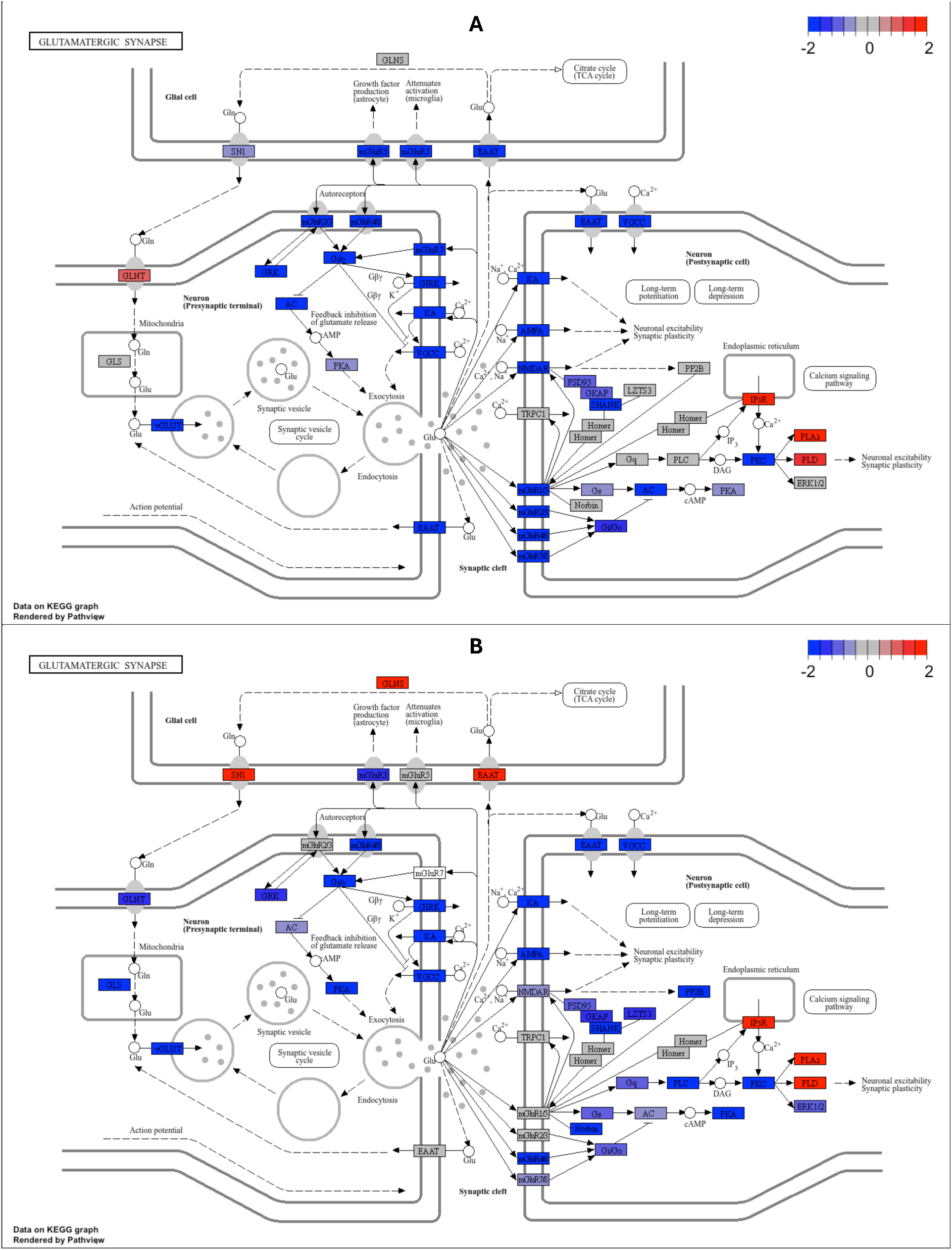
KEGG pathway representation of Glutamatergic synapse (hsa hsa04724) genes. Log_2_FC between *ST3GAL3^−/−^* (KO) and *ST3GAL3^+/+^* (WT) neurons differentiated via **(A)** protocol A or **(B)** protocol B were mapped onto their corresponding proteins in the KEGG *Glutamatergic synapse* pathway using Pathview. Boxes represent proteins encoded by the DEGs identified via DEseq2, coloured according to log_2_FC (red = upregulated, blue = downregulated in KO relative to WT).

We therefore examined genes associated with the *synaptic vesicle cycle* (Fig. 17 and S13). For this gene set as well, there appeared to be a general trend towards downregulation in ST3GAL3 KO–derived cortical neurons across both protocols. However, in protocol B, these genes displayed the largest fold changes among those that were significantly upregulated (Fig. 17B). These included *SLC1A3, TCIRG1, SLC6A5,* and *ATP6V0D2*. Of these genes, TCIRG1 was the only significantly identified by all three statistical methods for protocol A as well (Fig. 17A). Even in this case, the significantly downregulated genes—despite differing in the magnitude of change—showed high overlap between the two protocols. These comprised *SLC32A1, ATP6V1G2, CACNA1B, UNC13A, CPLX1, SLC17A6,* and *SNAP25* (Fig 17, and S13).

**Figure 17.**
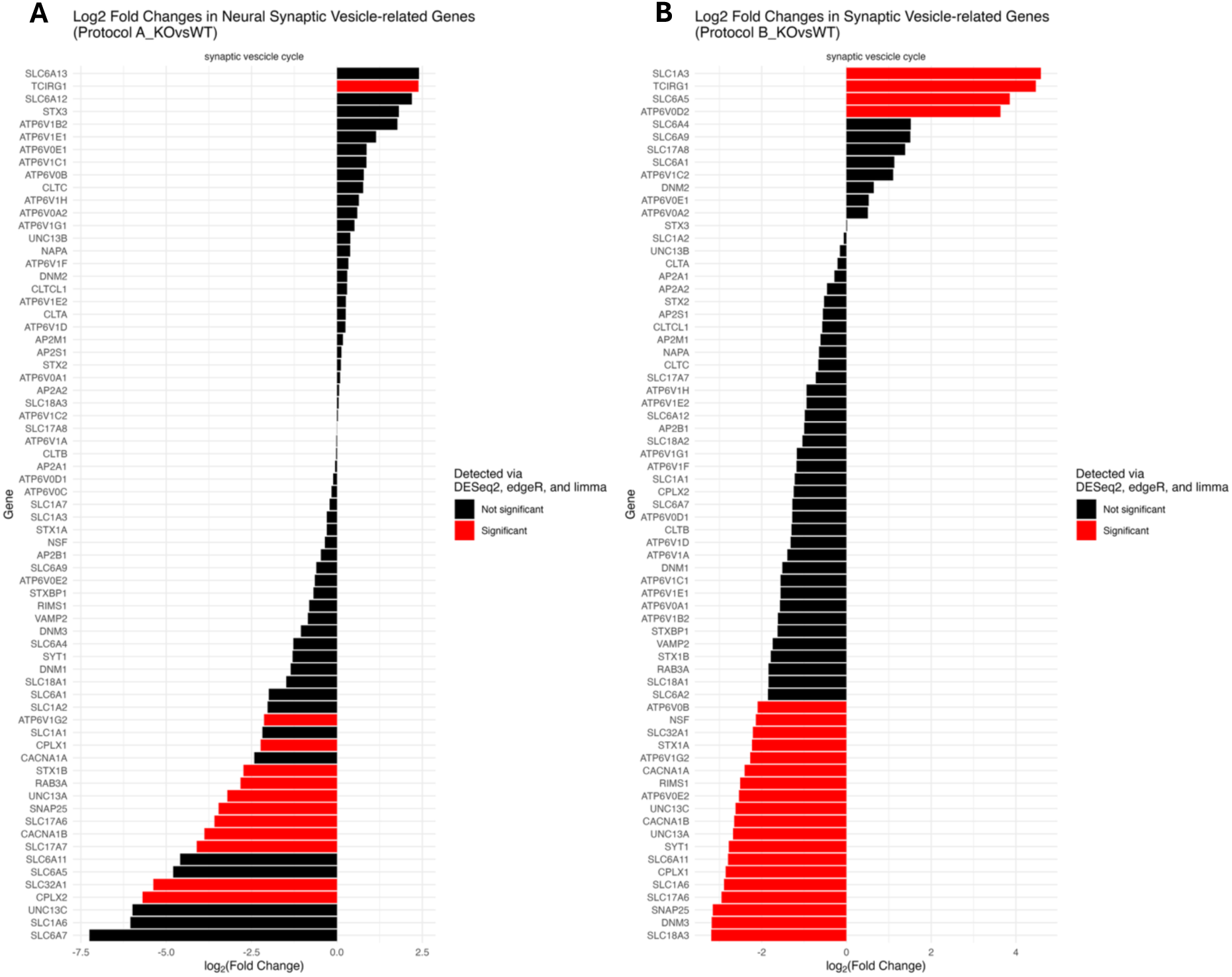
Log_2_FC of Synaptic vesicle-cycle (hsa04721) genes in *ST3GAL3^−/−^ (KO)* versus *ST3GAL3^+/+^* (WT) cultures. (**A**) Results from obtained via directed differentiation (protocol A). **(B)** Results from obtained via induced differentiation (protocol B) Genes were considered significant when consistently detected by DESeq2, edgeR, and limma. Significant differential expression is highlighted in red, while black bars denote non-significant genes. Log_2_FC values derived from DESeq2.

Turning to genes associated with *neuron-specific cell adhesion* molecules, we identified only one significantly upregulated gene in KO cultures under protocol A, as detected by all three statistical methods—PTPRN (Fig. 18A). Notably, this gene was instead significantly upregulated in KO cultures generated using protocol B, indicate of protocol specific effects of ST3GAL3 KO (Fig. 18B). Protocol B–derived ST3GAL3 KO neurons also showed significant upregulation of *ITGA8, ITGB8,* and *ITGAV* (Fig. 18B). Although these genes followed the same trend in protocol A, their changes did not reach statistical significance (Fig. 18A). Importantly, all significantly downregulated genes in ST3GAL3 KO neural cultures derived using protocol A were detected as significantly downregulated in protocol B. These included *NTNG2, NEGR1, CNTN2, NRXN1,* and *SLITRK3* (Fig. 18A,B). Notably, *NCAM1* and *CADM1* (also known as *synCAM*), which encode two well-known polysialylated proteins and substrates of ST3GAL3, were not identified as significant by all three statistical methods. *NCAM1* alone showed downregulation in KO neurons derived with protocol A, and this finding was supported solely by DESeq2 (Fig. S14).

**Figure 18.**
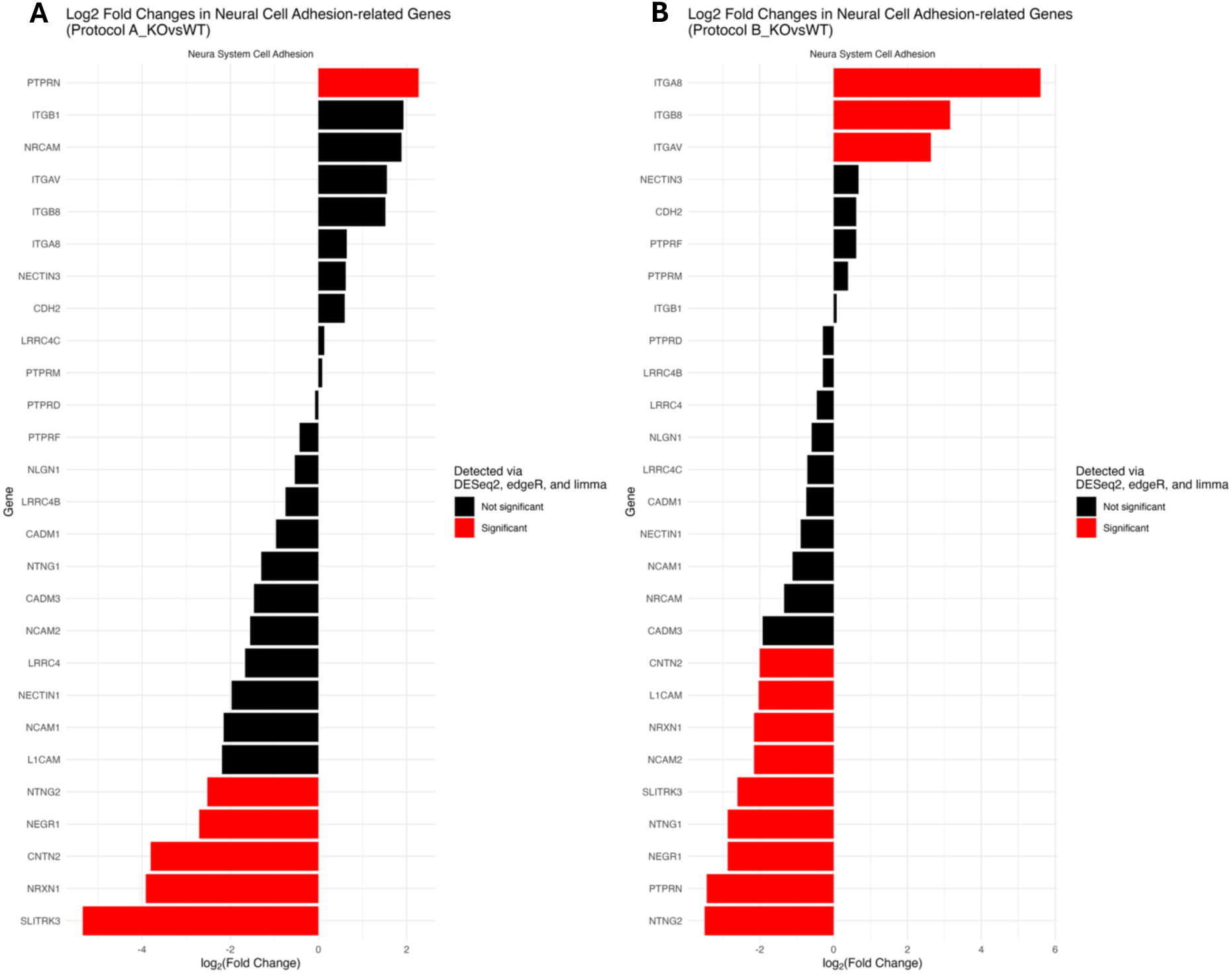
Log_2_FC of Neural cell adhesion genes in *ST3GAL3^−/−^* (KO) versus *ST3GAL3^+/+^* (WT) cultures. (**A**) Results from obtained via directed differentiation (protocol A). **(B)** Results from obtained via induced differentiation (protocol B) Gene set was extracted from cell adhesion molecules (hsa04514). Genes were considered significant when consistently detected by DESeq2, edgeR, and limma. Significant differential expression is highlighted in red, while black bars denote non-significant genes. Log_2_FC values derived from DESeq2.

Genes associated with *long-term depression* and *long-term potentiation* revealed pathways prone to interpretation to elucidate the impact of ST3GAL3 KO on the synaptic functioning of cortical neurons (Fig. 19). Indeed, whereas LTP-related pathways displayed a clear skewness towards downregulation, LTD-associated genes exhibited the highest number of significantly differentially expressed transcripts to be upregulated in STGAL3^KO^ neural cultures. Importantly, this pattern was evident in cortical neurons derived from both protocol A and protocol B (Fig 19, left panels).

**Figure 19.**
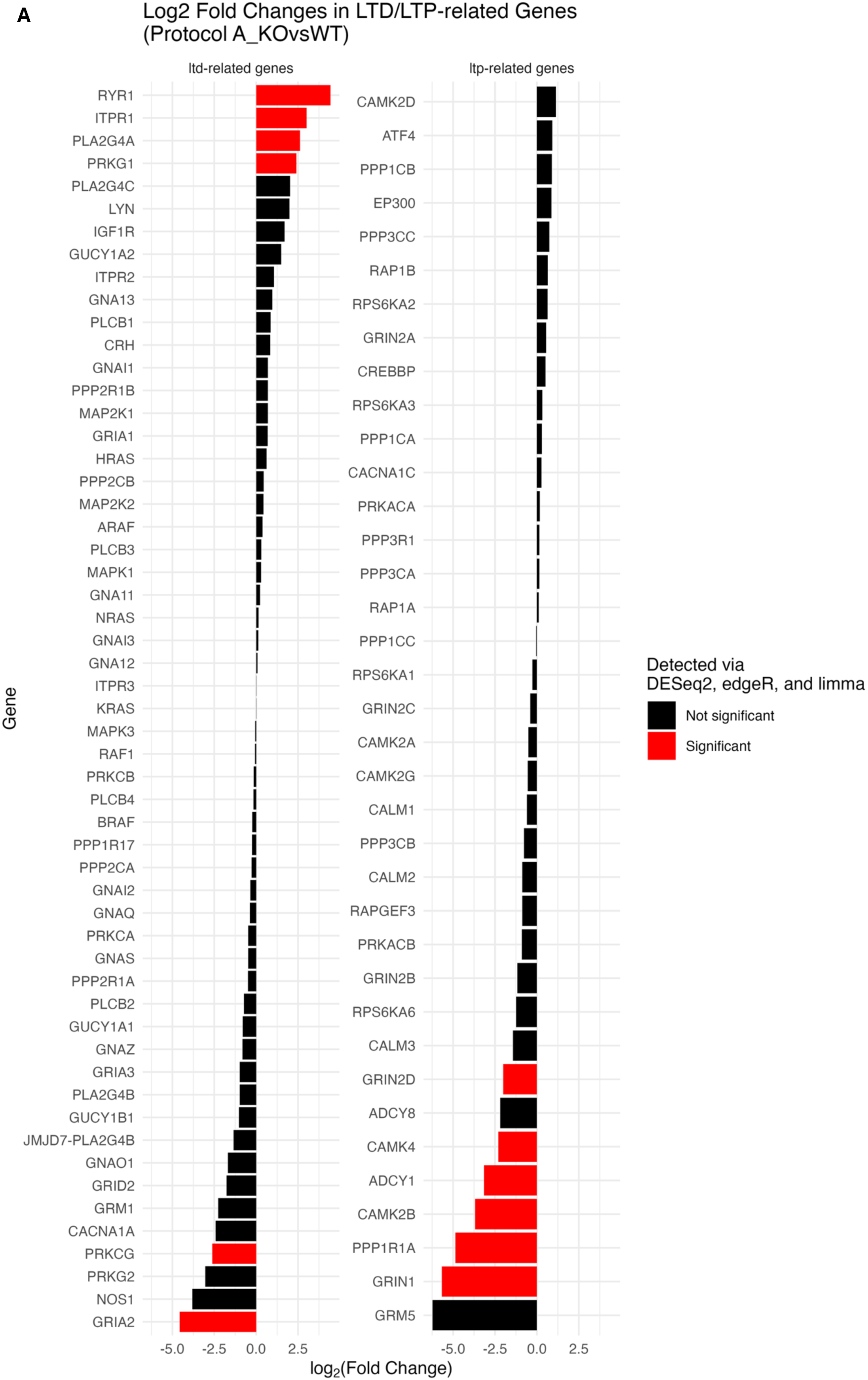
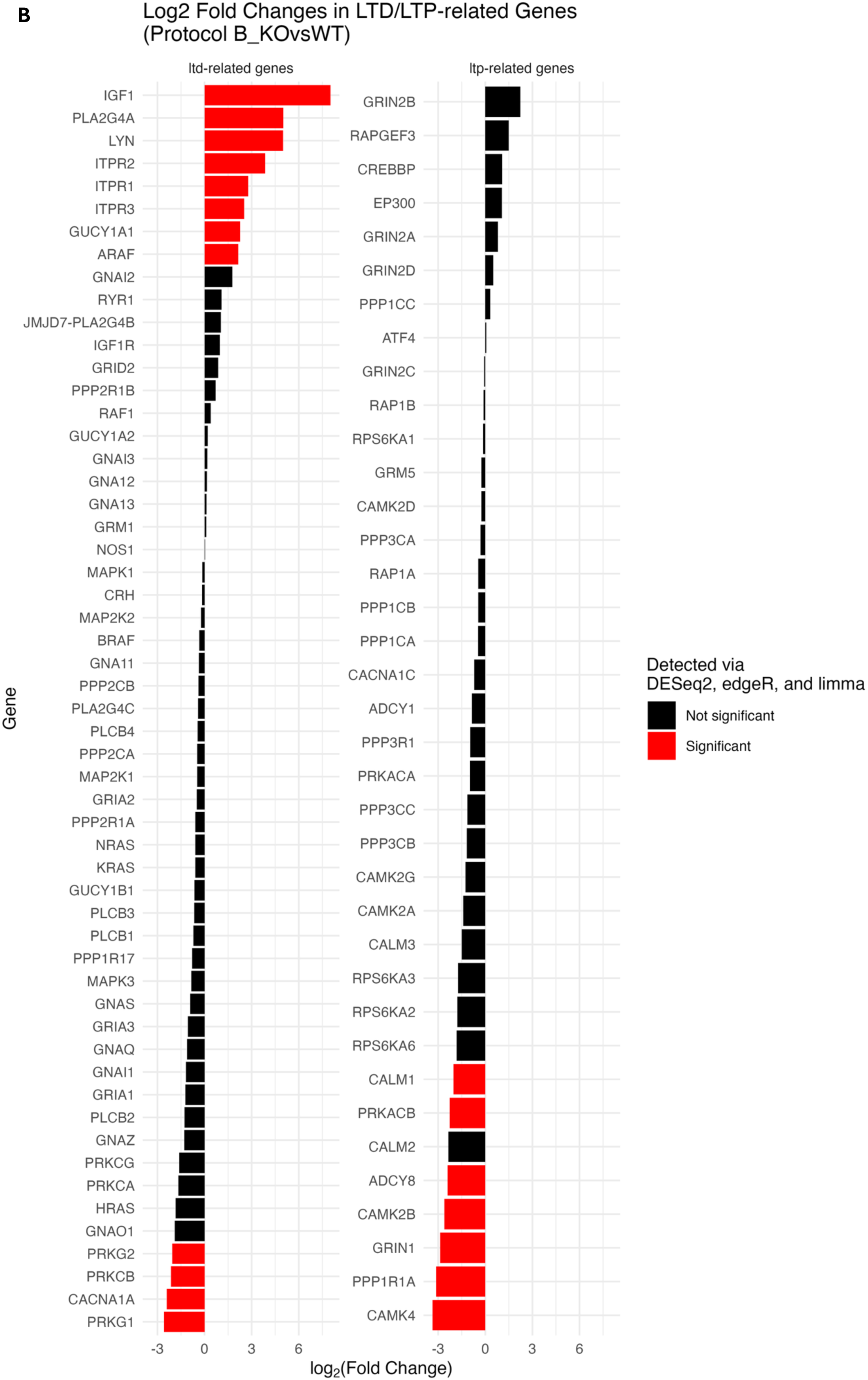
Log_2_FC of LTD (hsa04730) and LTP (hsa04720) genes in *ST3GAL3^−/−^* (KO) versus *ST3GAL3^+/+^*. **(WT) cultures. (A)** Results from obtained via directed differentiation (protocol A). **(B)** Results obtained via induced differentiation (protocol B). Genes were considered significant when consistently detected by DESeq2, edgeR, and limma. Significant differential expression is highlighted in red, while black bars denote non-significant genes. Log_2_FC values derived from DESeq2.

*Long-term potentiation*-related genes exhibited no signs of upregulation (Fig. 19, right panels). Instead, *CAMK4, CAMK2B, PPP1R1A,* and *GRIN1* were significantly downregulated in ST3GAL3 KO–derived neurons across both protocols (Fig, 19, right panels). Conversely, none of the significantly downregulated genes in the LTD gene set were detected in both protocol A and protocol B (Fig 19, left panels). Among these, *PRKG1* was significantly downregulated in KO cultures from protocol B but significantly upregulated in those from protocol A *ITPR1* and *PLA2G4A* were significantly upregulated in ST3GAL3 KO cultures across both protocols (Fig. 19, left panels).

Finally, in light of the CC enrichment analysis indicating axon-related gene involvement, we conducted a more detailed investigation of *axon guidance*–related genes. At first glance, no clear pattern was apparent (Fig. 20A,B). In protocol A–derived ST3GAL3 KO cortical neurons, three genes were significantly upregulated compared to WT, two of which *MYLS* and *EPHA2* were also significantly upregulated in protocol B–derived ST3GAL3 KO cortical neurons (Fig. 20)

**Figure 20.**
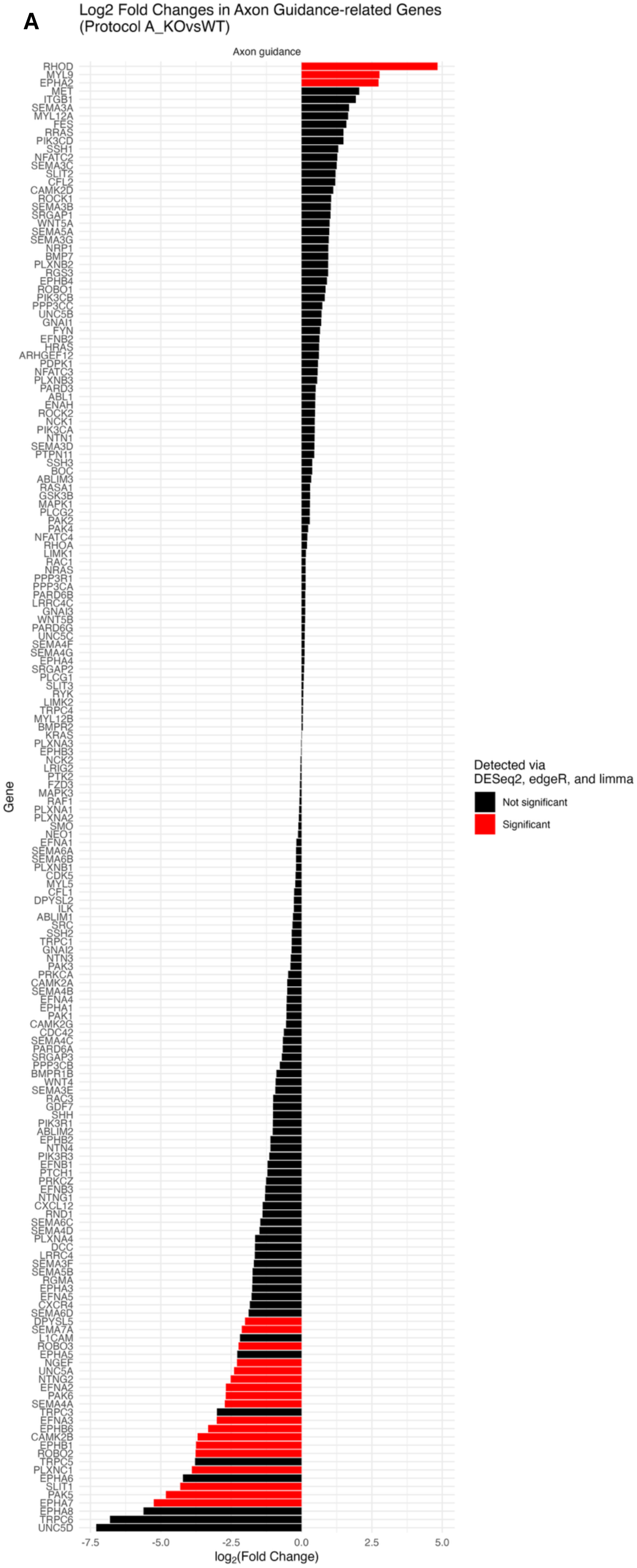
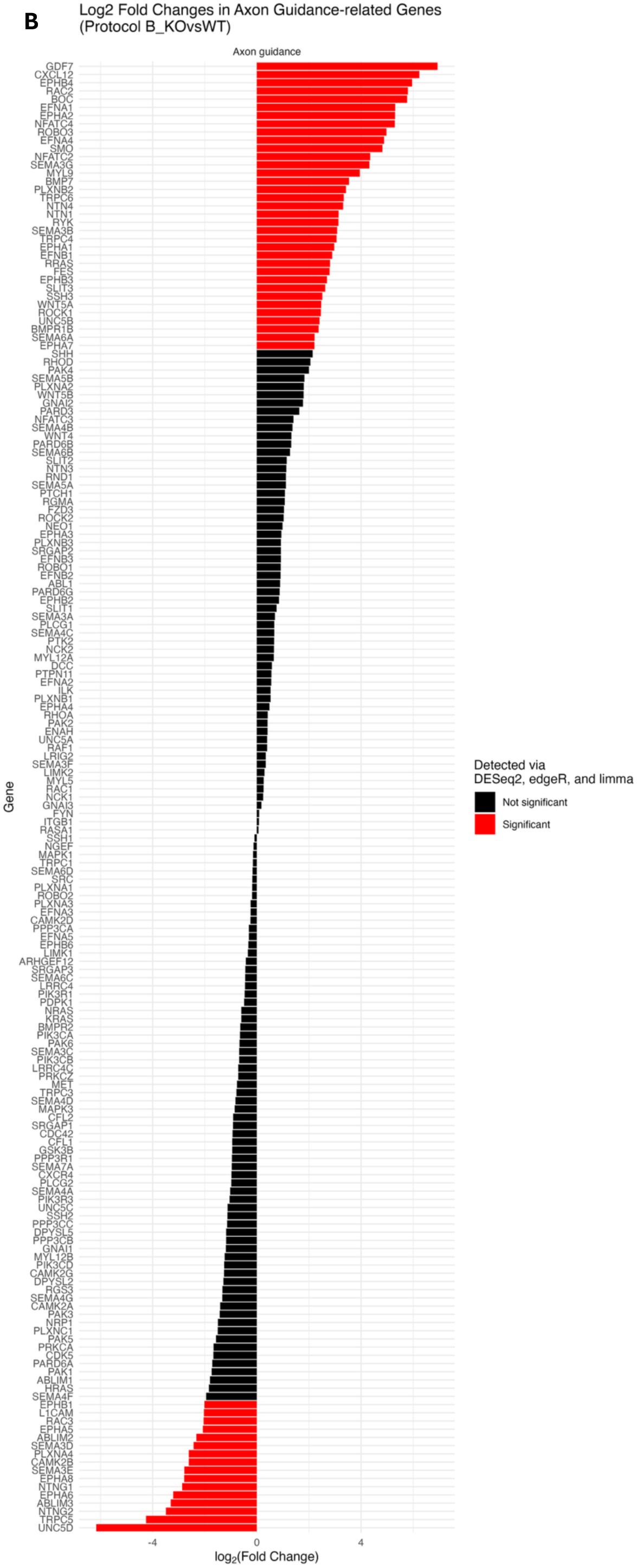
Log_2_FC of axon guidance (hsa04360) genes in *ST3GAL3^−/−^* (KO) versus *ST3GAL3^+/+^*. (WT) cultures. (A) Results from obtained via directed differentiation (protocol A). **(B)** Results from obtained via induced differentiation (protocol B) Gene set was extracted from cell adhesion molecules (hsa04514). Genes were considered significant when consistently detected by DESeq2, edgeR, and limma. Significant differential expression is highlighted in red, while black bars denote non-significant genes. Log_2_FC values derived from DESeq2.

Among the significantly downregulated genes, *UNC5D* showed the largest log_2_FC in ST3GAL3 KO neurons from both protocols, although it did not reach significance across all three statistical methods in protocol A–derived neurons (Fig. 20). Despite the relative abundance of significantly downregulated genes pertaining to this set in ST3GAL3 KO neurons compared to WT within both protocols, most of them were not shared across the two. Only exceptions were *NTNG2, CAMK2B,* and *EPHB1* (Fig. 20 and S17). Overall, findings regarding transcript levels of axon guidance-related genes remained inconclusive.

### Functional characterisation of ST3GAL3-deficient neuronal cultures

To investigate the role of ST3GAL3 in cortical neuronal networks at the functional level, we employed MEA recordings. This approach enabled the assessment of spontaneous electrophysiological activity across neuronal populations, providing insight into synaptic functionality in the absence of ST3GAL3. Following the generation of NPCs, ST3GAL3 KO and WT lines were plated onto MEA chips (6wellMEA200/30iR-Ti-rcr) and further matured into cortical neurons following the directed differentiation protocol (Fig. S5B). Spontaneous electrophysiological activity was recorded at DIV 30. Following quality control and exclusion of low-quality recordings, the final dataset comprised 11 independent KO replicates and 9 WT replicates, which were included in the downstream MEA analyses.

Electrophysiological parameters from individual replicates were initially averaged over time and electrodes to obtain a single mean value per technical replicate. These mean values were then subjected to PCA (Fig. 21 and S18 for loadings). The resulting PCA plot showed no clear separation between ST3GAL3 KO and WT neuronal lines based on the assessed parameters. This visual impression was confirmed by statistical analysis: PERMANOVA testing (Euclidean distance, 999 permutations) revealed no significant effect of genotype on the multivariate distribution of electrophysiological parameters (R² = 0.031, F = 0.580, p = 0.682). Furthermore, a permutation test for homogeneity of multivariate dispersions indicated no significant difference in dispersion between groups (F = 0.079, p = 0.786), suggesting comparable within-group variability. Consistently, statistical comparisons of individual mean parameters revealed no significant differences between ST3GAL3 KO and WT cortical neuron cultures (Fig. 22 and Tables S7,8).

**Figure 21.**
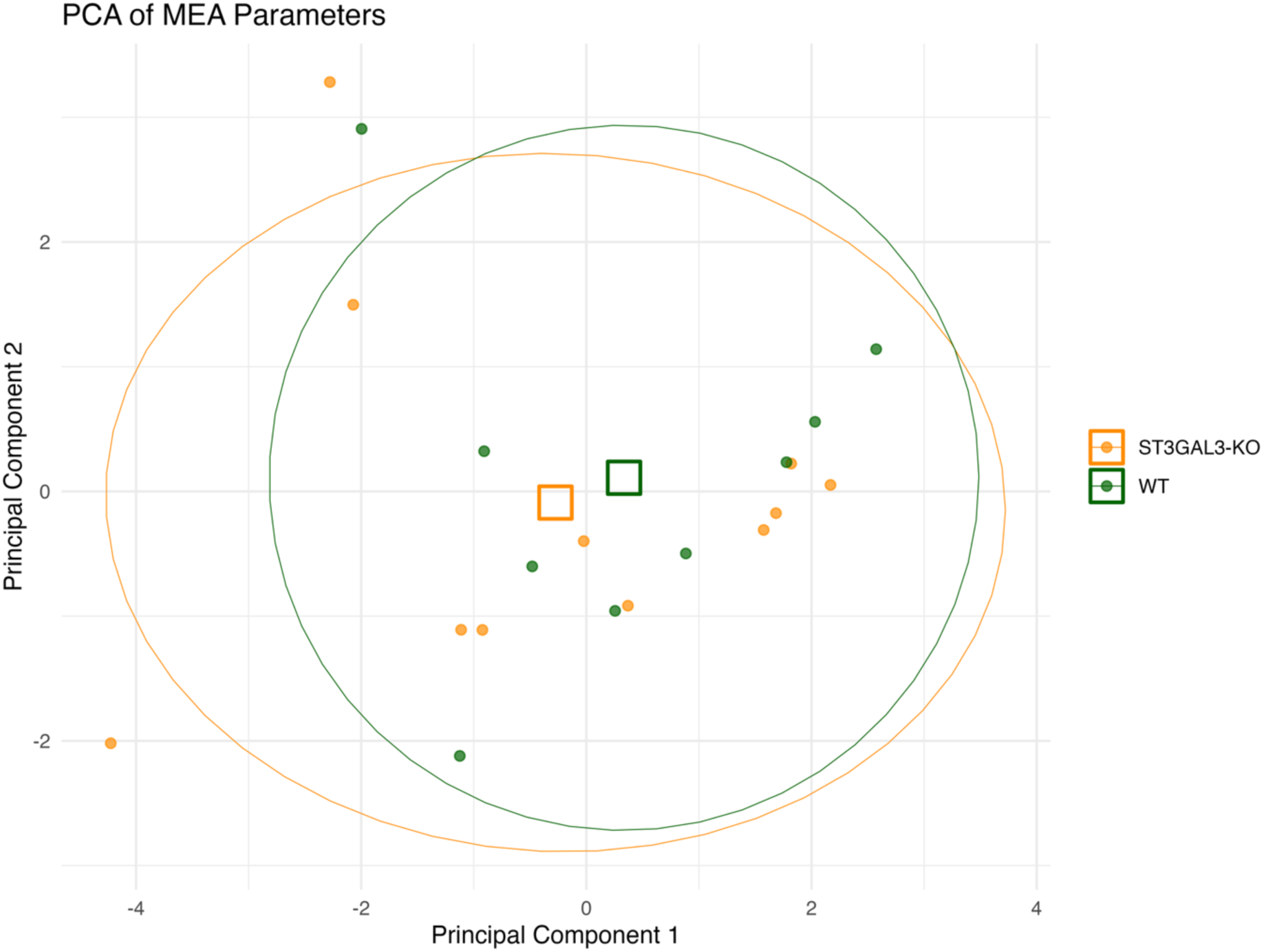
PCA of MEA-derived mean network parameters in *ST3GAL3^−/−^* (KO) and *ST3GAL3^+/+^* (WT) cortical neuron cultures. Scores for PC1 versus PC2 are shown; each point represents the replicate-level mean of the electrophysiological parameters. Open squares denote group centroids, calculated as the arithmetic means of PC1 and PC2 within genotype. Ellipses depict the 80% confidence regions. Colour coding: *ST3GAL3* KO (orange) and WT (green). No clear separation between genotypes was observed. This was confirmed by PERMANOVA (Euclidean distance, 999 permutations: R² = 0.031, F = 0.580, p = 0.682) and by a test for homogeneity of multivariate dispersions (F = 0.079, p = 0.786), indicating comparable within-group variability. PCA principal component analysis. PC1 principal component 1. PC2 principal component 2. MEA microelectrode array.

**Figure 22.**
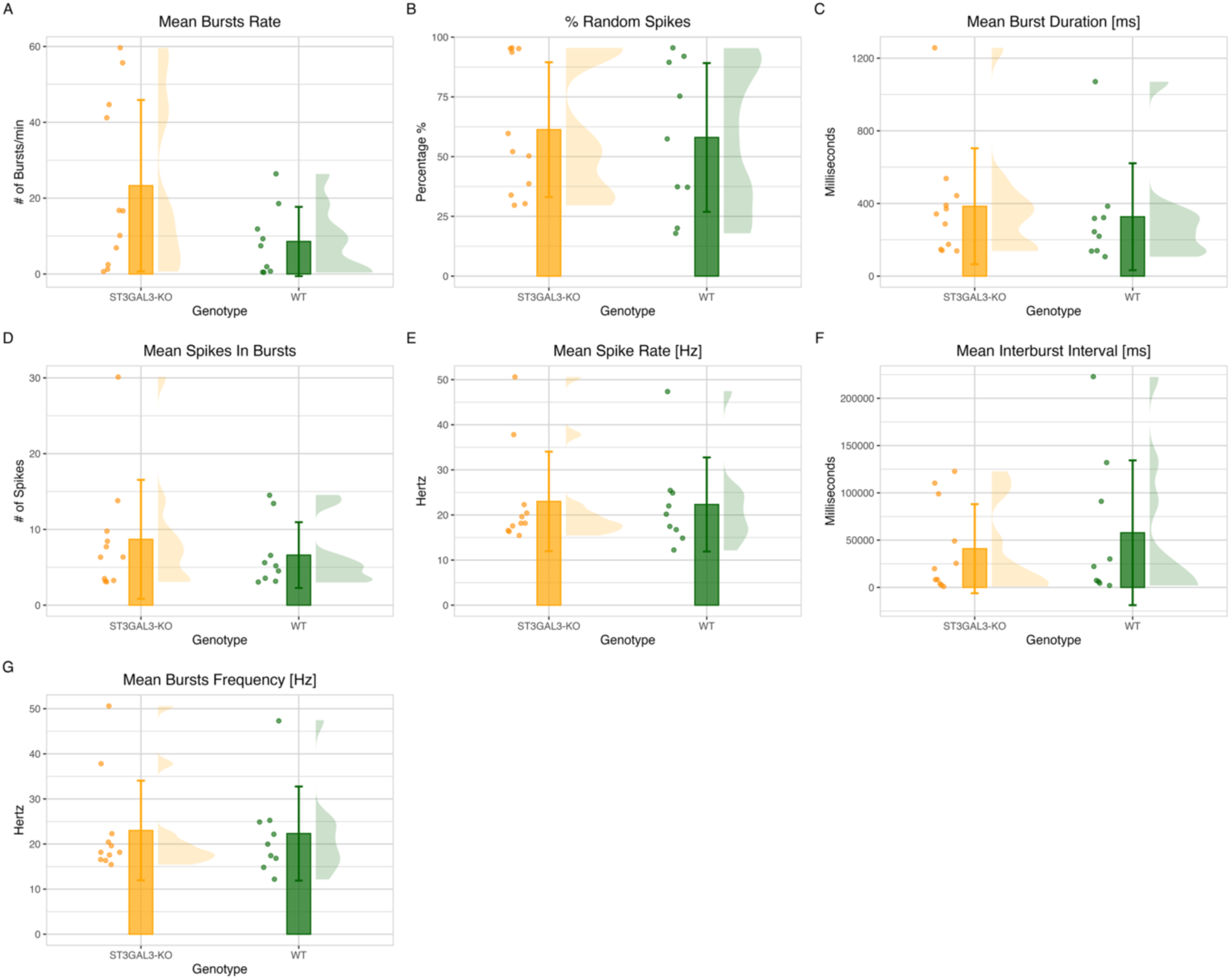
Comparison of electrophysiological mean-parameters between *ST3GAL3^−/−^* (KO) versus *ST3GAL3^+/+^* (WT) cortical neuron cultures. Each panel depicts the distribution of a specific mean parameter derived from MEA recordings. Each dot represents the mean value of an individual technical replicate (averaged across time and electrodes). Bars represent group mean, with error bars indicating the standard deviation. Pannels are: **(A)** mean burst rate (MBR), **(B)** percentage of random spikes (%RS), **(C)** mean burst duration (MBD), **(D)**, mean spikes in burst (MSiB), **(E)** mean spike rate (MSR), **(F)** mean inter-burst interval (MIBI), **(G)** mean burst frequency (MBF). No significant difference was detected between ST3GAL3 KO and WT neurons across either mean variable (Table S7,8). Yellow = ST3GAL3 KO; Green = WT. min minute. # number. Hz hertz. ms milliseconds.

To gain deeper insight into the electrophysiological behaviour of our cultures, we examined the characteristics of individual burst events. Notably, BD exhibited a marked divergence between ST3GAL3 KO and WT cultures that was not apparent in the per-well averaged data (Fig. 23). ST3GAL3 KO cortical neurons tended to produce bursts of longer duration than WT counterparts. Visualisation of log10-transformed BD values revealed that ST3GAL3 KO networks displayed a broader distribution, encompassing both longer and shorter bursts (Fig. 24). This increased variability, coupled with the high density of events around the median values in both groups, likely accounted for the lack of statistical significance in analyses based on mean BD.

**Figure 23.**
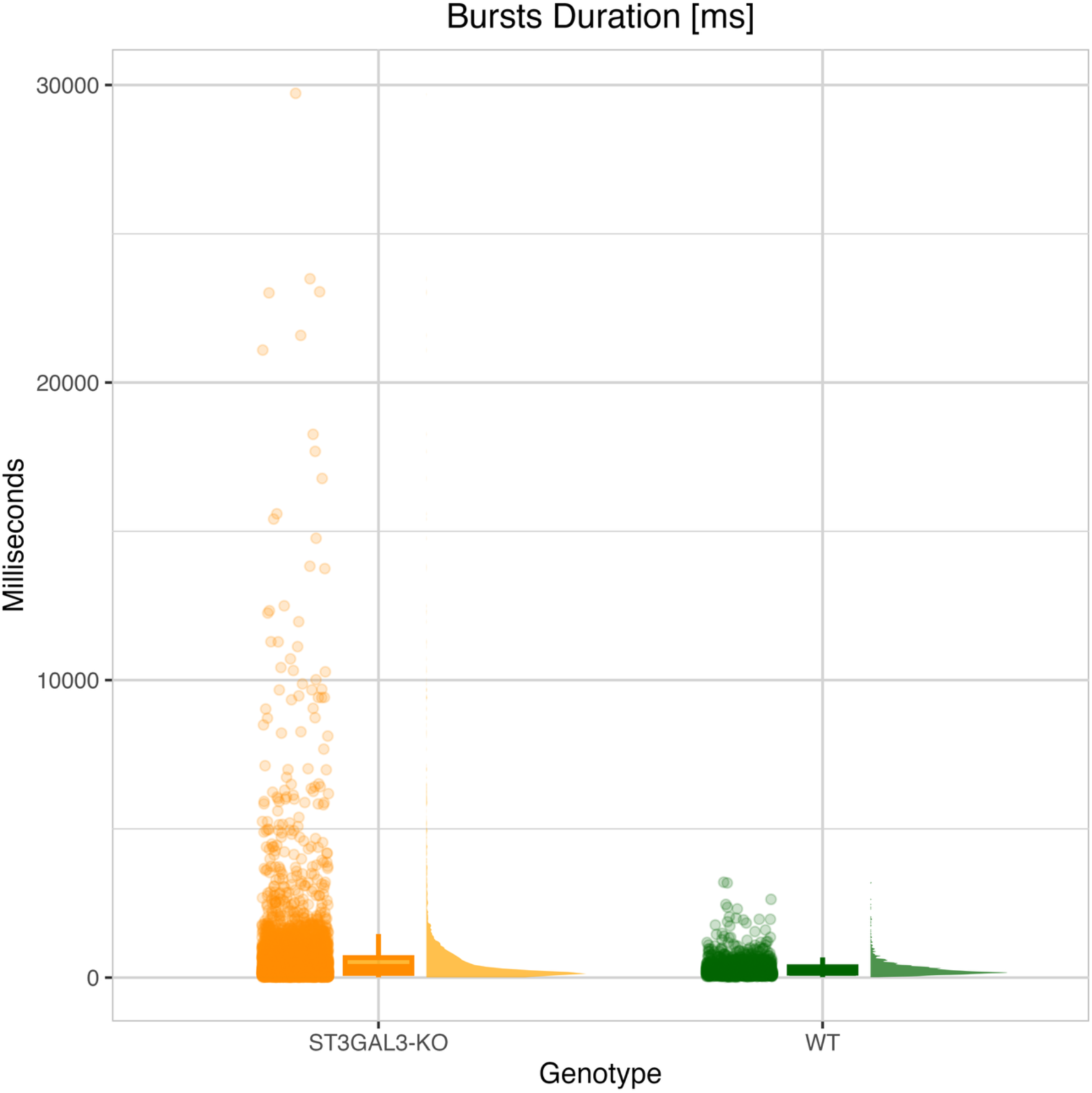
Raincloud plot of burst durations in *ST3GAL3^−/−^* (KO) and *ST3GAL3^+/+^* (WT) cortical neuron cultures. Each dot represents a single burst event detected from MEA recordings. The raincloud plot combines a boxplot (showing the median and interquartile range), a half violin plot (depicting the kernel density estimation of the distribution), and individual data points (scatter) to visualise both central tendency and data spread. ms milliseconds.

**Figure 24.**
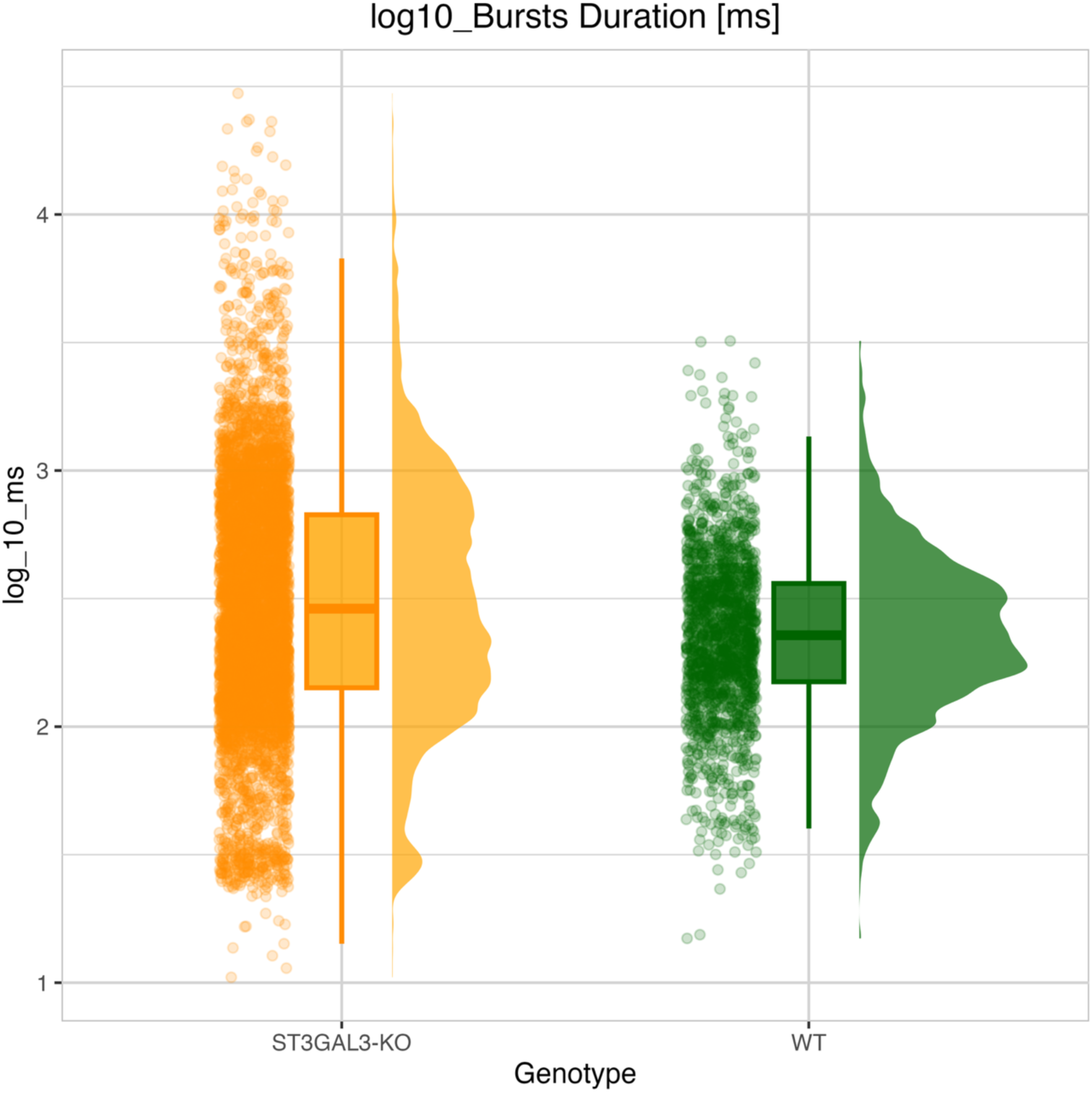
Raincloud plot of normalised burst durations in *ST3GAL3^−/−^* (KO) and *ST3GAL3^+/+^* (WT) cortical neuron cultures. Burst durations were log₁₀-transformed to reduce skewness and improve visualisation of distributional differences. Each dot represents a single burst event detected from MEA recordings. The raincloud plot combines a boxplot (showing the median and interquartile range), a half violin plot (depicting the kernel density estimation of the distribution), and individual data points (scatter) to visualise both central tendency and data spread. ms milliseconds.

To test the hypothesis that ST3GAL3 KO-derived cortical neurons display greater variability in burst duration, we conducted a chunked data analysis. The complete BD dataset from both genotypes was stratified into five categories based on percentile thresholds: Very Short (<10th percentile), Short (10th–25th percentile), Medium (25th–75th percentile), Long (75th–90th percentile), and Very Long (>90th percentile). This categorisation enabled the calculation and visualisation of the relative proportion of bursts in each category for both genotypes (Fig. 25). As anticipated, ST3GAL3 WT cultures displayed a more consistent and centralised burst-type distribution, with 67% of all bursts classified as Medium, in contrast to only 45% in ST3GAL3 *KO* cultures. In line with a shift towards prolonged burst durations, ST3GAL3 KO networks exhibited a greater propensity to generate both Long bursts (17% of events, compared to 7% in WT cultures) and Very Long bursts (13% of events, compared to only 2% in WT cultures). ST3GAL3 KO cultures also exhibited a slightly higher proportion of Very Short bursts (11% compared to 8% in WT cultures) (Fig. 25). The proportion of Short bursts was similar between genotypes, with ST3GAL3 KO cultures showing 15% and WT cultures 16% of events.

**Figure 25.**
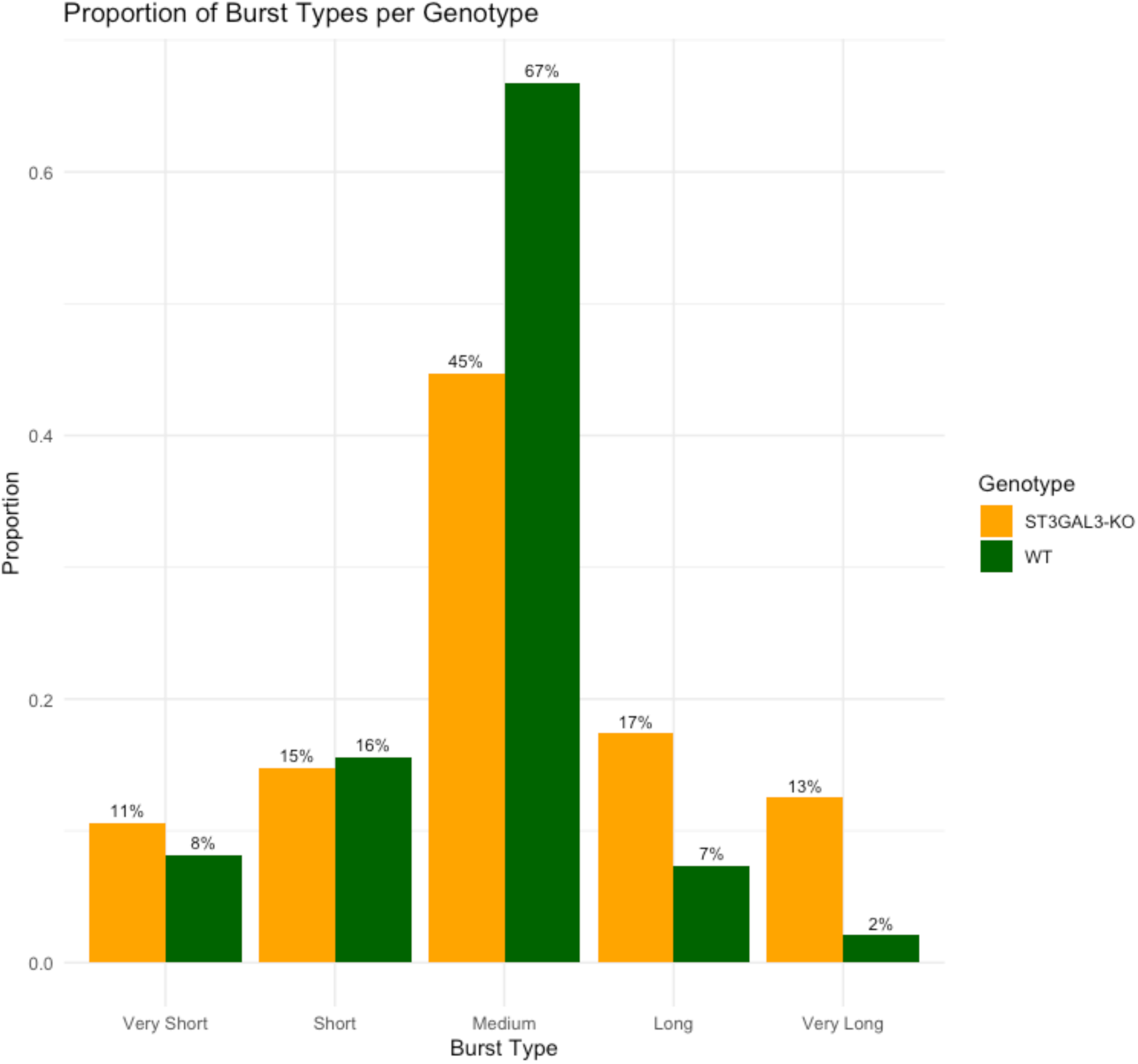
Proportional distribution of burst duration categories in *ST3GAL3^−/−^* (KO) and *ST3GAL3^+/+^* (WT) cortical neuron cultures. Burst durations from MEA recordings were stratified into five categories based on percentile thresholds calculated across the entire dataset: Very Short (<10th percentile), Short (10th–25th), Medium (25th–75th), Long (75th–90th), and Very Long (>90th percentile). Bars represent the proportion of total burst events falling into each category for each genotype.

To statistically evaluate whether genotype was predictive of burst category distribution, we performed a Bayesian multinomial logistic regression with burst category as the outcome variable and genotype as the fixed predictor, including a random intercept for recording well to account for repeated measures. The model compared the likelihood of Very Short, Medium, Long, and Very Long bursts relative to the reference category (Short bursts). ST3GAL3 KO neurons showed significantly increased odds of producing Very Short bursts (β = 0.44, 95% CI [0.01, 0.87]) and Very Long bursts (β = 0.73, 95% CI [0.27, 1.19]) compared to WT-cultures. Conversely, the odds of producing Medium bursts were significantly lower in the ST3GAL3 KO group (β = –0.57, 95% CI [-0.83, –0.32]). No significant difference was observed in the odds of generating Long bursts (β = 0.21, 95% CI [-0.13, 0.55]). These findings refine the chunked analysis results by quantifying the strength of association between genotype and specific burst duration categories, confirming that ST3GAL3 deficiency is associated with a redistribution of burst activity towards more extreme durations.

Based on the hypothesis that longer bursts might be followed by proportionally longer resting intervals, we sought to determine whether a correlation existed between BD and IBI. Scatterplot of individual events across the two variables quickly denoted absence of correlation between the two variables (Fig. S19A). Statistical analysis confirmed the observations, with Pearson’s correlation analysis yielding r = –0.388, p = 0.091, and a 95% confidence interval of [-0.709, 0.066], indicating no statistically significant association between burst duration and inter-burst interval on mean variables (Fig. S19B).

Throughout these analyses, we consistently observed a higher number of burst events in ST3GAL3 KO cultures compared with WT-derived counterparts (Fig S20). Looking at the results obtained from MBR, although the difference did not reach statistical significance, ST3GAL3 KO cultures exhibited a trend towards producing more bursts per minute (Fig. 22A). The elevated total number of detected bursts was partly attributable to a greater proportion of burst-detecting electrodes: 35.4% of all recording electrodes in ST3GAL3 KO cultures detected bursts, compared with 17.4% in WT cultures (Fig. 21A). However, per-well analyses again revealed no significant genotype differences, likely due to high variability across wells (Fig. 21B). Coupled with the presence of two more replicates in the ST3GAL3 KO-derived culture group, these factors account for the greater overall abundance of bursts observed in the ST3GAL3 KO compared to WT but remains inconclusive regarding a potential hyper excitability of ST3GAL3 KO-derived cortical neurons.

No difference on BF was observed between ST3GAL3 KO and WT-derived cortical neurons (Fig. 26 and 22G). Furthermore, although an apparent correlation between BD and BF can be observed (Fig. S22), this likely reflects an artefact of the burst detection parameters rather than a genuine physiological association. Specifically, because longer bursts inherently contain more spikes, the calculated intra-burst frequency tends to converge towards the predefined detection settings, producing a spurious correlation.

**Figure 26.**
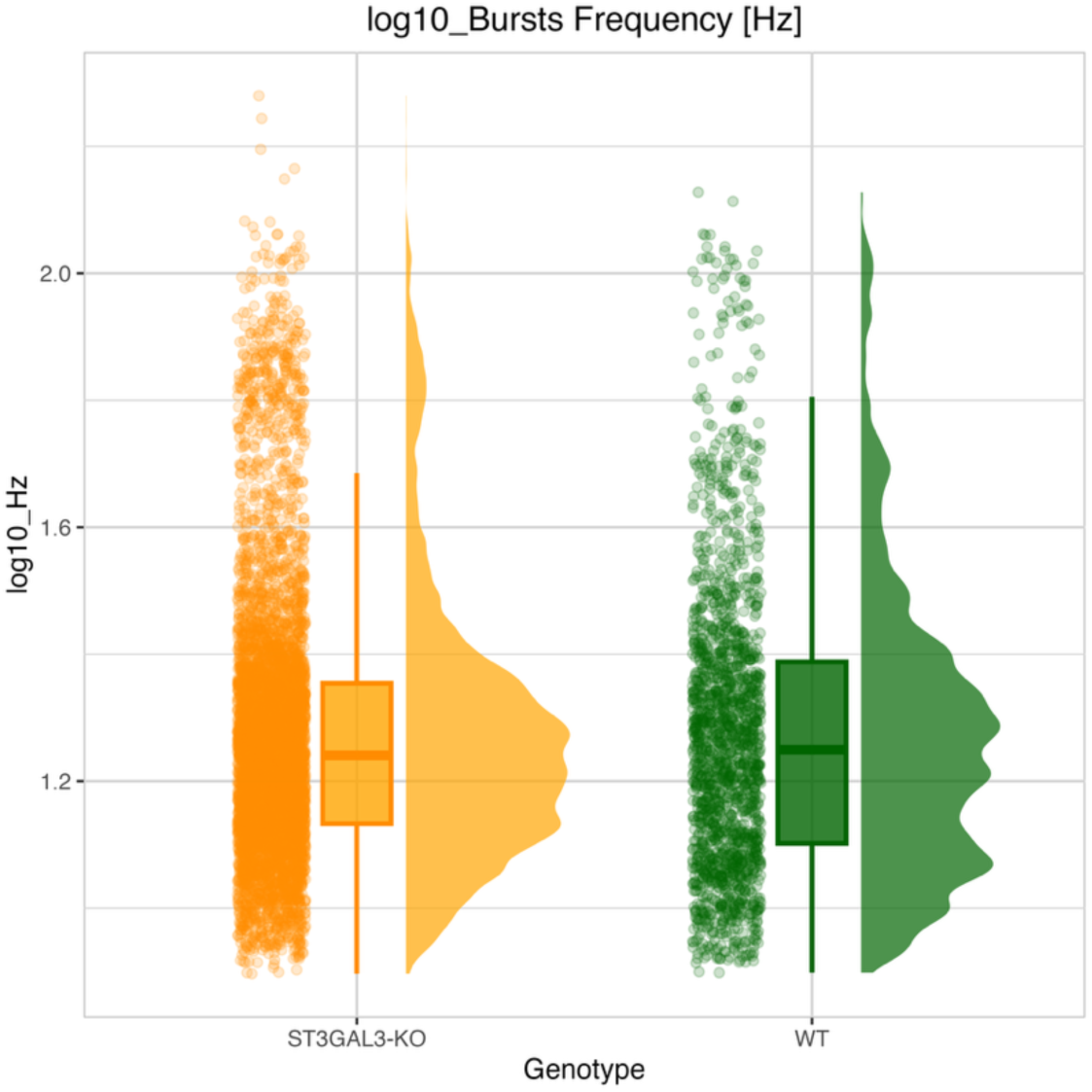
Raincloud plot of normalised burst frequencies in *ST3GAL3^−/−^* (KO) and *ST3GAL3^+/+^* (WT) cortical neuron cultures. Burst frequencies were log₁₀-transformed to reduce skewness and improve visualisation of distributional differences. Each dot represents a single burst event detected from MEA recordings. The raincloud plot combines a boxplot (showing the median and interquartile range), a half violin plot (depicting the kernel density estimation of the distribution), and individual data points (scatter) to visualise both central tendency and data spread. Hz hertz.

In agreement with the lack of genotype-dependent differences on BF, analysis of SiB data revealed an almost perfect linear correlation with BD (Fig. S23A,B) (Pearson’s correlation analysis yielding r = 0.9057, p = < 3.953e-08, and a 95% confidence interval of [-0.773, 0.963]. This relationship reflects the straightforward principle that bursts containing more spikes are, at constant frequency, of longer duration. Consequently, further SiB analysis was not pursued since it would not yield additional insights beyond those already captured by the BD analysis.

## Discussion

This study aimed at investigating the role of ST3GAL3 in regulating the E/I balance of human cortical neurons, employing a ST3GAL3 KO iPSC model with targeted inactivation of *ST3GAL3*. Using the two most widely applied protocols for differentiating iPSCs into cortical neurons, i.e. directed and induced differentiation, by means of integrated morphological, transcriptomic, and electrophysiological analyses, we found that loss of ST3GAL3 disrupts the molecular and functional dynamics of E/I networks, with direct consequences for synaptic signalling. These observations build upon previous genetic and clinical evidence implicating *ST3GAL3* in NDDs such as NSARID, EIEE, and ADHD. Importantly, through this human-based model, our results provide the first direct evidence that ST3GAL3 plays a critical role in maintaining E/I balance, thereby situating it within the broader neurodevelopmental framework and underscoring its essential contribution to synaptic integrity.

*ST3GAL3* encodes a sialyltransferase responsible for initiating glycoprotein polysialylation, a post-translational modification that plays a pivotal role in regulating both cell–cell adhesion and interactions with the extracellular environment. The negative charge of sialyl epitopes contributes to repulsive forces that may be fundamental in limiting excessive synapse formation. Based on the convergence of clinical evidence and the enzyme’s established biochemical role, we hypothesised that inactivation of *ST3GAL3*, drives critical alterations in synaptic dynamics that ultimately contribute to the NDDs described above.

Molecular characterisation using RNAseq data of cortical neuron cultures derived from ST3GAL3 KO lines, compared with their isogenic WT controls, revealed a pronounced divergence in transcriptomic profiles. The use of two distinct differentiation methodologies enabled cross-validation of the findings, minimising protocol-specific effects and strengthening the case for biologically relevant results.

Consistent with our hypothesis, GO analysis of DEGs identified synaptic domains among the most profoundly affected across both differentiation methods. These included processes such as synaptic vesicle-mediated transport, neurotransmitter release, pre– and post-synaptic components, synaptic membrane organisation, and synaptic specialisation. Furthermore, analyses of neurons generated through induced differentiation revealed transcriptomic upregulation in pathways related to cell–matrix and cell–cell adhesion in ST3GAL3 KO^-^derived cultures. While these latter findings suggest a potential causal link between ST3GAL3 deficiency and hyperconnectivity of cortical neurons—possibly leading to imbalances in E/I synapse formation—this hypothesis remains speculative, as a conclusive causal correlation cannot not be established from transcriptomic datasets.

GO analysis strongly indicated the involvement of specific functional complexes in synaptic regulation and cell–cell interactions. Across both differentiation protocols, the analysis consistently highlighted ion transmembrane transport and protein/lipid-binding functions, underscoring a central role for ST3GAL3 in calcium-dependent ion transport and synaptic vesicle fusion—processes essential for synaptic transmission and neuronal excitability. Ion channel–related genes were predominantly downregulated, while protein/lipid-binding genes, including those associated with extracellular matrix and cell–cell communication, displayed more variable expression changes. These protocol-independent findings emphasise ST3GAL3’s pivotal contribution to synaptic regulation and network stability.

KEGG enrichment analysis allowed us to investigate the transcriptomic signature of ST3GAL3 KO cortical neurons in relation to selected E/I-related gene-sets. The substantial concordance in both significance and direction of transcriptional changes across all gene sets strengthens the robustness of our findings and underscores the critical role of *ST3GAL3* in regulating these genes, irrespective of methodological differences.

Relative to the isogenic WT cultures, ST3GAL3 KO neurons exhibited a broad downregulation of synaptic genes across both excitatory and inhibitory pathways. It is tempting to speculate on these findings in respect to the overall E/I balance. Whereas both excitatory and inhibitory synaptic genes were downregulated in ST3GAL3 KO neurons, the most pronounced reductions seemed to have occurred in GABAergic genes. This pattern suggests that ST3GAL3 disruption may have a disproportionately strong impact on GABAergic neurons, thereby shifting the E/I balance towards the excitatory side, a notion that is supported by the MEA results. Notably, the profound transcriptional changes observed in excitatory or inhibitory synapses might reflect compensatory mechanisms within the neuronal network, aiming to restore the E/I balance disrupted by *ST3GAL3* loss. In this respect, transcripts encoding GABAergic receptor subunits, including *GABRA2, GABBR1,* and *GABRB3*, were among the most strongly affected, suggesting that *ST3GAL3* loss may exert especially strong effects on GABA-receptive postsynaptic elements. Regardless of the precise causal sequence, these findings clearly establish ST3GAL3 as essential for the proper functioning of both glutamatergic and GABAergic synapses.

As noted above, ST3GAL3 is critical for polysialylation, a process that so far has not been shown to directly affect GABA or glutamate receptor function. Therefore, the observed changes in these gene sets are likely an indirect consequence of disruptions to other synaptic processes. In this regard, our findings on genes related to synaptic vesicle cycle and neuronal cell adhesion—processes in which sialylation has been shown to play a pivotal and more direct role—may represent the primary drivers of the ST3GAL3 KO–associated disruptions. The synaptic vesicle cycle emerged as one of the most affected pathways, suggesting a previously unrecognised role for ST3GAL3 in this process. Future research should clarify ST3GAL3’s precise contribution to the vesicle cycle and its interaction with the genes identified here. Notably, our analysis of neuronal cell adhesion molecules revealed a new set of genes and proteins potentially regulated by *ST3GAL3*, including *NTNG2, NEGR1, CNTN2, NRXN1,* and *SLITRK3*—showing strong, protocol-independent downregulation in ST3GAL3 KO neurons. It is important to note that these results do not rule out NCAM1 and CADM1, which did not display differential expression, as key contributors in this context. Their interaction with ST3GAL3 may not be detectable at the transcriptomic level but could be evident at the proteomic level.

Our analysis of long-term potentiation– and long-term depression-related genes further underscores the importance of ST3GAL3 in processes critical for maintaining the E/I balance. Specifically, we observed downregulation of long-term potentiation-associated genes—particularly *CAMK4, CAMK2B, PPP1R1A,* and *GRIN1*—alongside upregulation of long-term depression-associated genes, notably *ITPR1* and PLA2G4A. This pattern suggests that *ST3GAL3* disruption may shift the equilibrium between these synaptic plasticity mechanisms. Determining whether these transcriptomic changes are causal or consequential remains an important question for future research.

As these results are derived from transcriptomic analyses, they do not permit causal inferences or definitive conclusions regarding *ST3GAL3*’s direct regulatory role on the identified genes and related proteins. Nonetheless, these transcriptomic datasets provide valuable insights into ST3GAL3’s role in synaptic homeostasis and its influence on E/I balance, with the strength of the findings reinforced by their consistency across two distinct neuronal differentiation protocols. This cross-validation reduces the likelihood of protocol-specific artefacts and strengthens the evidence for biologically relevant effects. While the data reveal reproducible alterations in pathways central to synaptic regulation and suggest previously unrecognised roles for ST3GAL3 in additional, unexplored biological processes.

Functionally, our MEA recordings revealed that ST3GAL3 KO neurons displayed aberrant bursting patterns, particularly prolonged burst durations and heightened variability. These disruptions mirror the electrophysiological signatures of E/I imbalance observed in other NDD models. These findings position *ST3GAL3* as a critical molecular determinant of neuronal excitability. By integrating our molecular findings—highlighting altered expression of genes involved in cognition, learning, and memory—with the observed functional consequences of *ST3GAL3* deficiency, we find strong alignment with clinical observations in patients with biallelic *ST3GAL3* loss-of-function leading to NSARID and EIEE The cognitive and learning delays reported in these patients are mirrored in our cultures through gene expression alterations in pathways underpinning these processes. Likewise, albeit speculative, the clinically observed epileptic seizures may be reflected in our model by the spontaneous occurrence of abnormally prolonged bursts.

Additionally, the statistical approach used for the MEA recordings in this study highlights the need for more sophisticated analytical methods to characterise the electrophysiological behaviour of cultures. Relying solely on mean value comparisons flattens temporal scales and diminishes the unique advantages that MEA offers over other electrophysiological techniques. For future studies, using MEA chips and systems with higher capacity and resolution would be advantageous. Modern platforms can accommodate 24-well plates—compared with the 6-well plates used here—and feature over 200 electrodes per well, rather than the 9 used in this study. Such upgrades would improve technical replicability, enhance spatial resolution, and provide deeper insights into neuronal network dynamics.

The principal strength of this study lies in its integrative approach, combining isogenic iPSC models, dual differentiation protocols, and both transcriptomic and electrophysiological analyses to comprehensively examine the role of ST3GAL3 in synaptic function. Notably, the models implemented are entirely human-based, without the use of non-human components such as rat astrocytes, which could otherwise introduce confounding variables or obscure genotype-driven differences. The isogenic knockout model allows for precise attribution of phenotypes to *ST3GAL3* loss by eliminating background genetic variability, thereby ensuring that any observed differences in neuronal development, network activity, or gene expression can be confidently ascribed to the absence of ST3GAL3. Additionally, the convergence between molecular and functional deficits provides compelling evidence for a causal relationship between ST3GAL3 deficiency and E/I imbalance. The methodologies employed—such as RNA sequencing and MEA analysis—are high-throughput and offer superior resolution and scalability compared to traditional techniques like qRT-PCR and patch clamp, thereby enhancing the analytical power and translational relevance of our findings.

However, some limitations must be acknowledged. While the model captures key aspects of cortical neurodevelopment, it does not recapitulate the full in vivo complexity of neural circuits, including glial interactions and other neural subtypes connectivity. Furthermore, although the study highlights ST3GAL3’s role in synaptic regulation, it does not dissect the specific downstream pathways through which altered sialylation leads to the observed functional impairments. Deeper proteomic investigations will be necessary to bridge the gap between transcriptomic alterations and the observed functional phenotypes. While the use of isogenic iPSC lines represents a powerful and controlled system for dissecting gene-function relationships, it also necessitates additional validation. Complementary studies using patient-derived iPSC lines or *in vivo* models will be essential to confirm the translatability and clinical relevance of our findings. Future work should expand on these mechanistic links and explore the translational relevance of *ST3GAL3* modulation in disease-specific contexts.

Taken together, the convergence between our functional and molecular data supports the emerging view that sialylation is not a peripheral biochemical process but a central regulatory mechanism for synaptic communication and plasticity. These broad implications, together with the previously reported involvement of *ST3GAL3* in ADHD, support the hypothesis that E/I imbalance represents a shared pathophysiological substrate across NDDs and highlight *ST3GAL3* as a pivotal factor within this framework. This aligns with the transdiagnostic framework increasingly adopted in neuroscience, which recognises that neurodevelopmental and psychiatric disorders often share underlying neurobiological disruptions despite divergent clinical presentations.

In conclusion, our findings delineate a mechanistic link between ST3GAL3 and synaptic regulation in human neurons. By bridging transcriptomic alterations with functional deficits, this study advances our understanding of how a single molecular perturbation can reverberate across multiple neural processes. The ST3GAL3 KO iPSC model presented here provides a valuable platform for future research into therapeutic strategies aimed at restoring E/I synaptic balance.

