## Supplementary material for "*ST3GAL3* loss-of-function disrupts synaptic integrity and excitatory/inhibitory cortical dynamics"

**Table S1. Reagents and plasmids used for lentiviral transduction.**

| **Name** | **Manufacturer** |
|---|---|
| psPAX2 lentiviral packaging vector | Addgene |
| pMD2.G lentiviral packaging vector | Addgene |
| Lenti-X Concentrator | Takara Bioscience |
| Puromycin | InvivoGen |
| G418 | Sigma-Aldrich |
| Polybrene | Sigma-Aldrich |
| Primocin | InvivoGen |
| **Transfer vectors** | |
| pLVX-EF1α-(Tet-On-Advanced)-IRES-G418(R) | Department of Cognitive Neuroscience, Radboudumc, Nijmegen, The Netherlands |
| pLVX-(TRE-thight) Ngn2- PGK-Puromycin(R) | Department of Cognitive Neuroscience, Radboudumc, Nijmegen, The Netherlands |
| pLVX- (TRE-thight) - Ascl1-PGK-Puromycin (R) | Department of Cognitive Neuroscience, Radboudumc, Nijmegen, The Netherlands |

**Table S2. Sequences of primers used for the detection of TET3G, Ngn2, and Ascl1 genes**

| **Target** | **Primer type** | **Sequence (5'->3')** |
|---|---|---|
| TET3G | Forward | CTGGGAGTTGAGCAGCCTAC |
| | Reverse | AGAGCACAGCGGAATGACTT |
| Ascl1 | Forward | GTCCTGTCGCCCACCATCTC |
| | Reverse | CAGCAGCTCTTGTTCCTCTG |
| Ngn2 | Forward | AGACGGTGCAGCGCATCAAGAA |
| | Reverse | AGCGTCTCGATCTTCGTGAGCT |

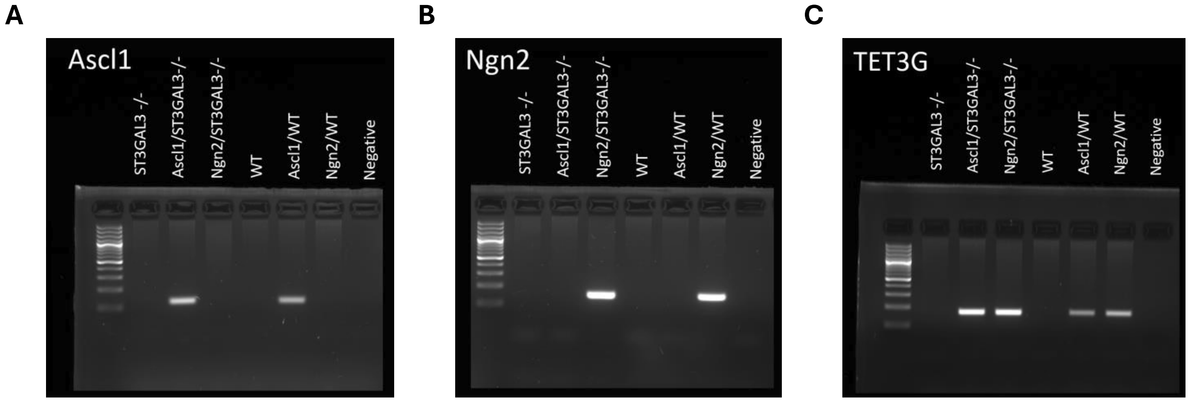


**Figure S1. Transgene genotyping of Ascl1+ and Ngn2+ iPSCs.** PCR amplification of genomic DNA demonstrates the presence of transgenes used for inducible neuronal differentiation. **(A)** Amplicons corresponding to murine Ascl1 (~150 bp), **(B)** murine Ngn2 (~150 bp), and **(C)** TET3G transactivator (~150 bp) were detected by agarose gel electrophoresis in isogenic wild-type (WT), ST3GAL3 KO (*ST3GAL3*$^{-/-}$), and dual-transgene iPSC lines. Genotypes include WT, Ascl1/WT, Ngn2/WT, *ST3GAL3*$^{-/-}$, *Ascl1/ST3GAL3*$^{-/-}$, and *Ngn2/ST3GAL3*$^{-/-}$. 'Negative' denotes the negative control condition, in which PCR was performed using water instead of genomic DNA. The presence of bands at the expected molecular weight confirms stable genomic integration of each construct.

### Table S3. Primary antibodies used for immunocytochemistry

| Target Protein | Host | Target Location | Dilution | Supplier |
|---|---|---|---|---|
| OCT3/4 | mouse | Intracellular | 1:50 | Santa Cruz Biotechnology |
| SSEA-4 | mouse | Extracellular | 1:200 | Invitrogen |
| TRA-1-60 | mouse | Extracellular | 1:50 | Santa Cruz Biotechnology |
| VE-CAD | goat | Extracellular | 1:100 | R&D Biosystems |
| FOXA2 | mouse | Intracellular | 1:250 | Santa Cruz Biotechnology |
| PAX6 | mouse | Intracellular | 1:100 | Invitrogen |
| SOX2 | goat | Intracellular | 1:250 | Santa Cruz Biotechnology |
| β3-tubulin | guinea pig | Intracellular | 1:500 | Synaptic Systems |
| VGLUT1 | rabbit | Intracellular | 1:1000 | Synaptic Systems |
| VGLUT2 | rabbit | Intracellular | 1:500 | Synaptic Systems |
| SV2B | rabbit | Intracellular | 1:500 | Synaptic Systems |
| GAD67 | mouse | Intracellular | 1:70 | Millipore |
| VGAT | rabbit | Intracellular | 1:500 | Synaptic Systems |

### Table S4. Secondary antibodies used for immunocytochemistry

| Target Species | Host | Fluorophore | Dilution | Supplier |
|---|---|---|---|---|
| rabbit | donkey | Alexa Fluor 488 | 1:500 | Invitrogen |
| goat | donkey | Alexa Fluor 488 | 1:500 | Invitrogen |
| mouse | donkey | Alexa Fluor 555 | 1:500 | Invitrogen |
| guinea pig | donkey | Alexa Fluor 647 | 1:500 | Jackson IR |

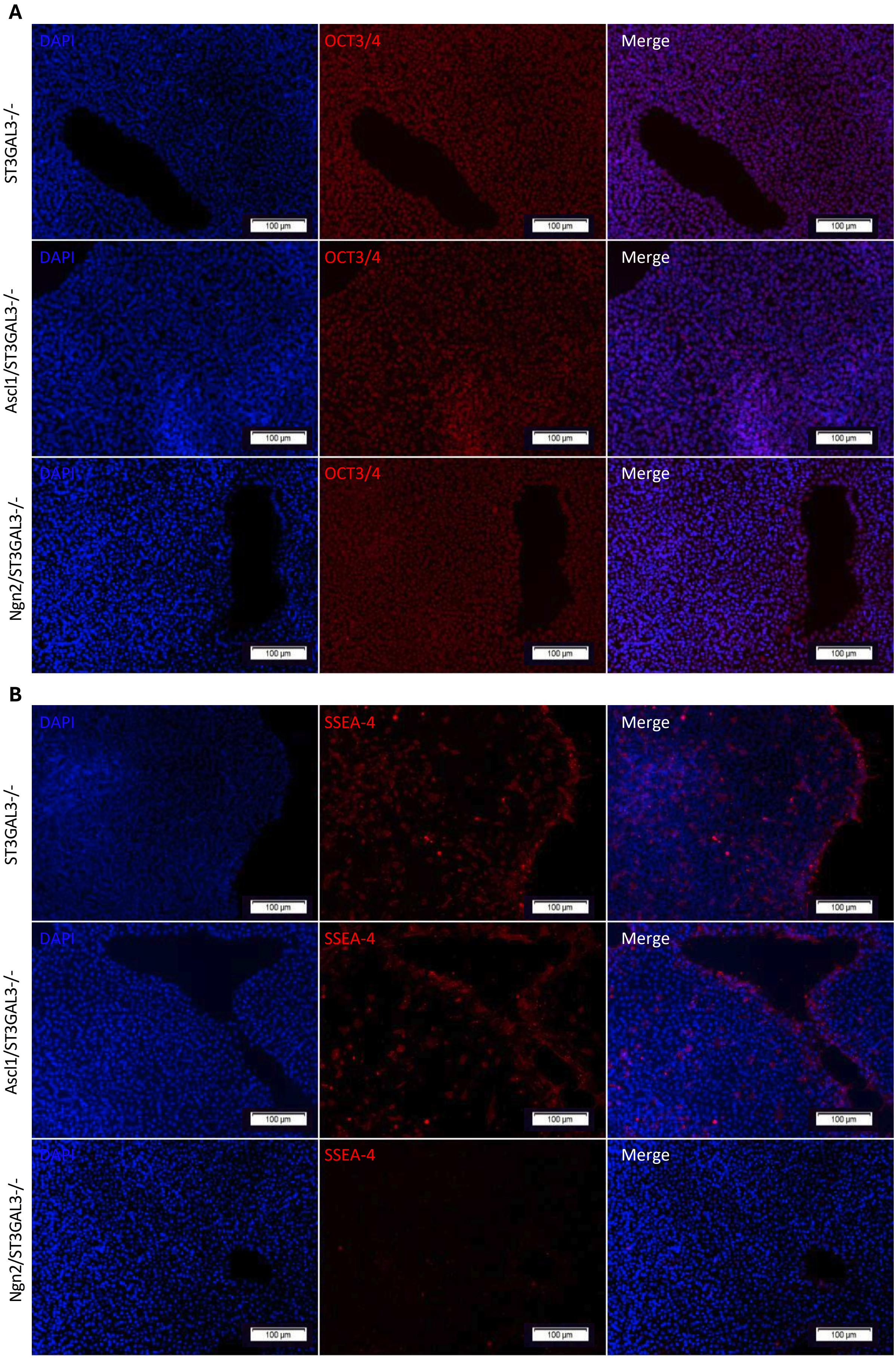
A
DAPI
OCT3/4
Merge
ST3GAL3-/-
Ascl1/ST3GAL3-/-
Ngn2/ST3GAL3-/-
100 µm
B
DAPI
SSEA-4
Merge
ST3GAL3-/-
Ascl1/ST3GAL3-/-
Ngn2/ST3GAL3-/-
100 µm

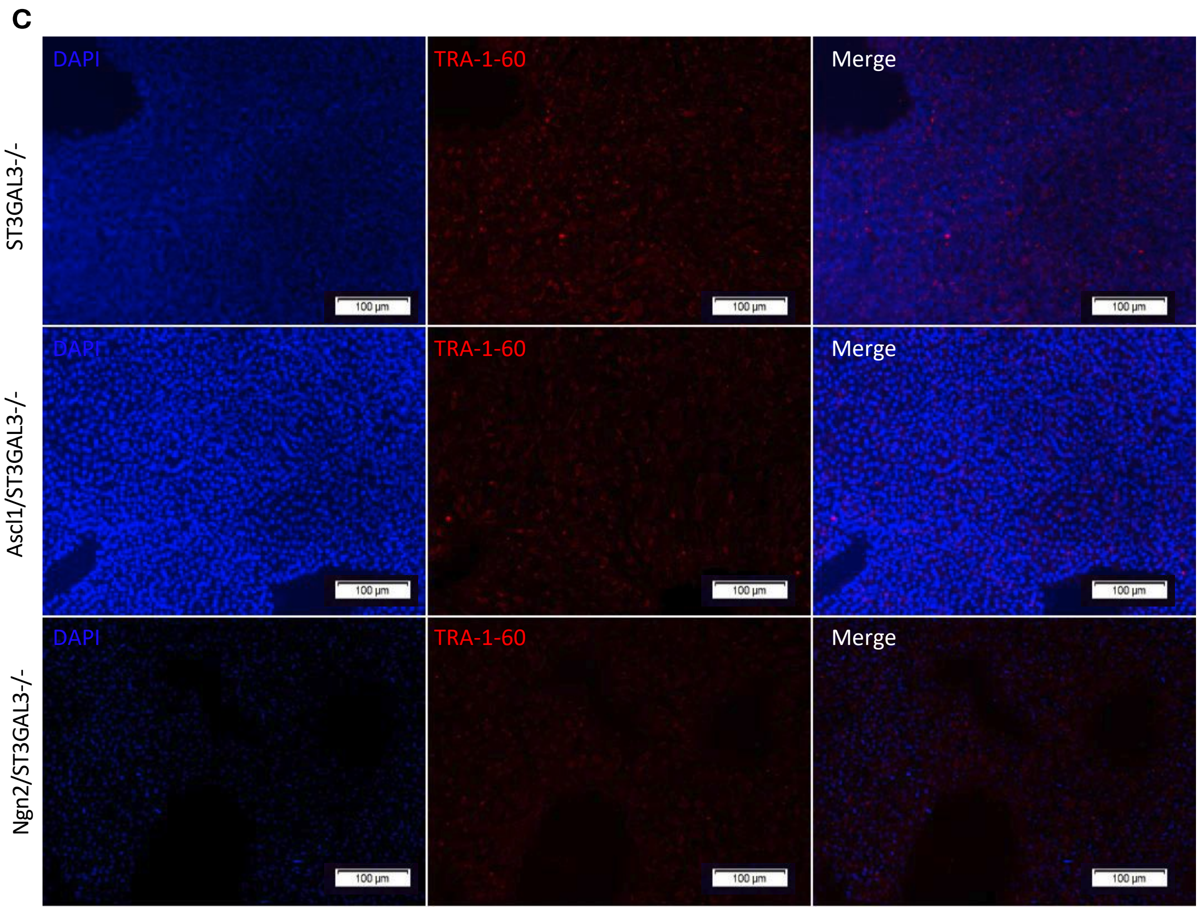


**Figure S2. ICC analysis of pluripotency in ST3GAL3-deficient iPSC lines.** Representative images illustrating the expression of canonical pluripotency markers in *ST3GAL3* $^{-/-}$, *Ascl1/ST3GAL3* $^{-/-}$, and *Ngn2/ST3GAL3* $^{-/-}$ iPSCs. All lines exhibited robust staining for **(A)** OCT3/4, **(B)** SSEA-4, and **(C)** TRA-1-60, confirming maintenance of the pluripotent state following genetic manipulation.

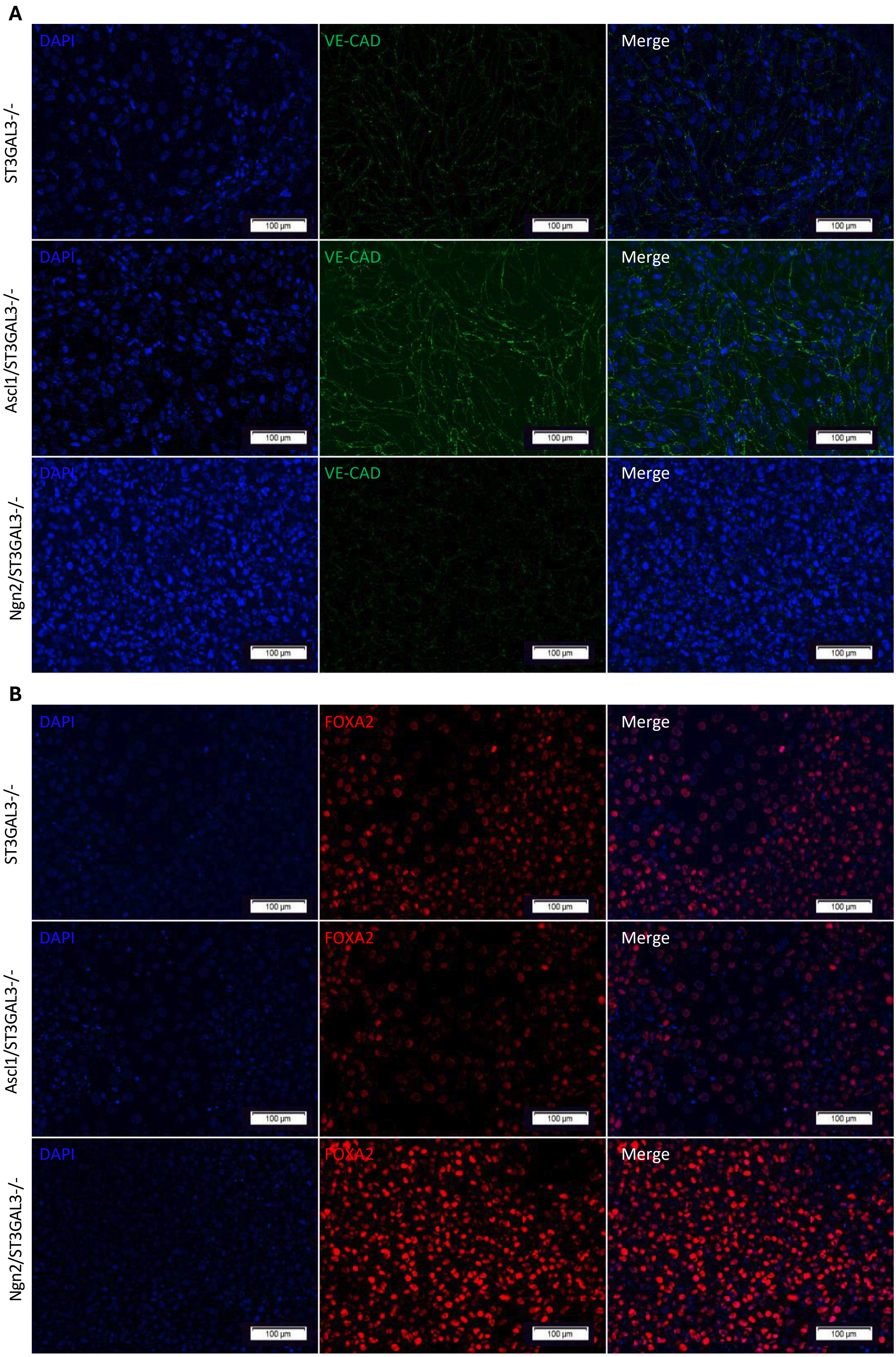
A
DAPI
VE-CAD
Merge
ST3GAL3-/-
Ascl1/ST3GAL3-/-
Ngn2/ST3GAL3-/-
100 µm
B
DAPI
FOXA2
Merge
ST3GAL3-/-
Ascl1/ST3GAL3-/-
Ngn2/ST3GAL3-/-
100 µm

C

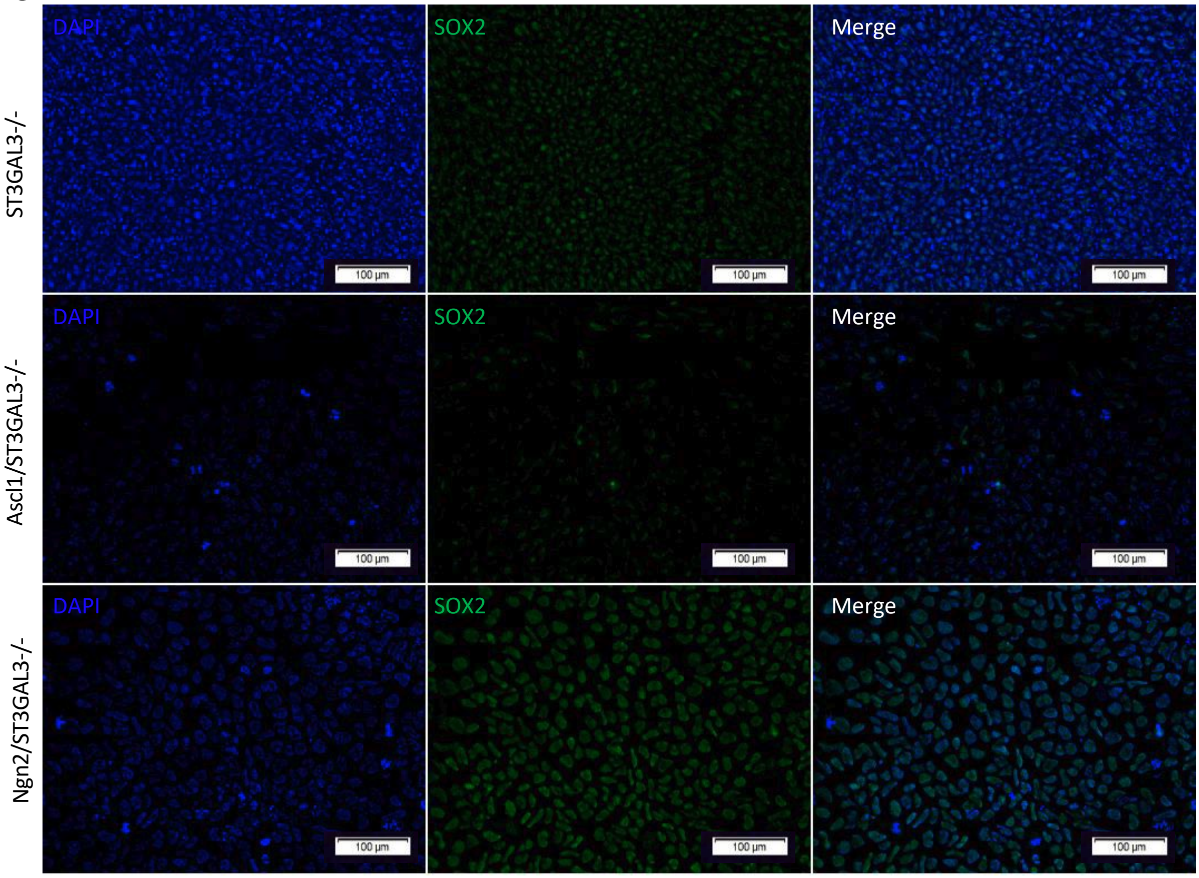

DAPI
SOX2
Merge
ST3GAL3-/-
Ascl1/ST3GAL3-/-
Ngn2/ST3GAL3-/-
100 µm


D

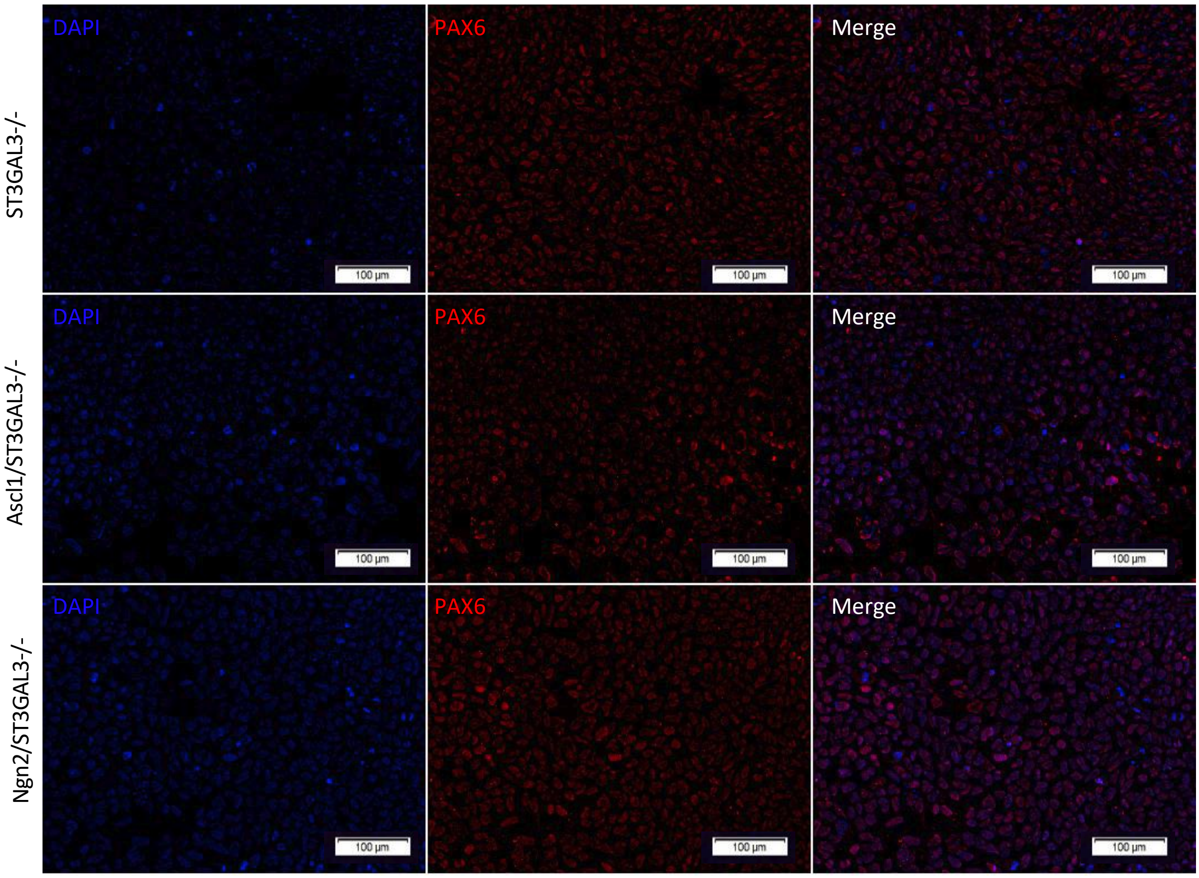

DAPI
PAX6
Merge
ST3GAL3-/-
Ascl1/ST3GAL3-/-
Ngn2/ST3GAL3-/-
100 µm

**Figure S3. ICC analysis of trilineage differentiation potential in ST3GAL3-deficient iPSC lines.** Representative images showing expression of lineage-specific markers in *ST3GAL3*$^{-/-}$, *Ascl1/ST3GAL3*$^{-/-}$, and *Ngn2/ST3GAL3*$^{-/-}$ iPSCs following directed differentiation into the three germ layers. All lines demonstrated positive staining for **(A)** vascular endothelial cadherin (VE-CAD; mesoderm), **(B)** forkhead box A2 (FOXA2; endoderm), **(C)** sex-determining region Y-box 2 (SOX2; ectoderm), and **(D)** paired box 6 (PAX6; ectoderm), confirming preservation of trilineage differentiation capacity following genetic manipulation.

**Table S5. Media and reagents list employed for induced differentiation of iPSCs into neurons.**

| Reagent | Provider |
|---|---|
| B-27 (503), serum free | ThermoFisher Scientific |
| Cytosine b-D-arabinofuranoside hydrochloride (Ara-C) | Sigma |
| DMEM/F-12 | ThermoFisher Scientific |
| DMEM, High Glucose, GlutaMAX | ThermoFisher Scientific |
| Doxycycline | Sigma |
| DPBS, No calcium, No magnesium | ThermoFisher Scientific |
| DPBS, Calcium, Magnesium | ThermoFisher Scientific |
| StemMACS™ iPS-Brew XF, human | Miltenyi Biotec |
| Forskolin | Sigma |
| GlutaMAX supplement | ThermoFisher Scientific |
| G418 | Sigma-Aldrich |
| Human Recombinant laminin-521 | Biolamina |
| Mouse laminin | Sigma |
| MEM non-essential amino acid solution (NEAA) | Sigma |
| Matrigel | Corning |
| N-2 Supplement (1003) | ThermoFisher Scientific |
| Neurobasal | ThermoFisher Scientific |
| Penicillin/Streptomycin | Sigma |
| Poly-L-Ornithine | Sigma |
| Primocin | ThermoFisher Scientific |
| Puromycin | Sigma |
| PBS | Sigma |
| Recombinant Human BDNF | PeproTech |
| Recombinant Human Neurotrophin 3 (NT3) | Promocell - PromoKine |
| Y-27632 | Stem Cell Technologies |

**Table S6. Media and reagents list employed for directed differentiation of iPSCs into neurons.**

| | |
|---|---|
| STEMdiff Neural Induction Medium | Stem Cell Technologies |
| STEMdiff SMADi Neural Induction Supplement | Stem Cell Technologies |
| Gentle Cell Dissociation Reagent | Stem Cell Technologies |
| DMEM/F-12 | ThermoFisher Scientific |
| DPBS, No calcium, No magnesium | ThermoFisher Scientific |
| Y-27632 | Stem Cell Technologies |
| Poly-L-Ornithine | Sigma |
| Mouse Laminin | Sigma |
| Matrigel | Corning |
| STEMdiff Neural Rosette Selection Reagent | Stem Cell Technologies |
| AggreWell 800 24-well Plate | Stem Cell Technologies |
| Anti-Adherence Rinsing Solution | Stem Cell Technologies |
| 37 µm Reversible Strainer | Stem Cell Technologies |
| STEMdiff Neural Progenitor Medium 05833 | Stem Cell Technologies |
| STEMdiff Neural Progenitor Freezing Medium 05838 | Stem Cell Technologies |
| BrainPhys Neuronal Medium | Stem Cell Technologies |
| NeuroCult SM1 Neuronal Supplement | Stem Cell Technologies |
| N2 Supplement-A | Stem Cell Technologies |
| Human Recombinant BDNF | Stem Cell Technologies |
| Human Recombinant GDNF | Stem Cell Technologies |
| Dibutyryl-cAMP | Stem Cell Technologies |
| Ascorbic Acid | Stem Cell Technologies |
| Primocin | Thermo Fisher Scientific |

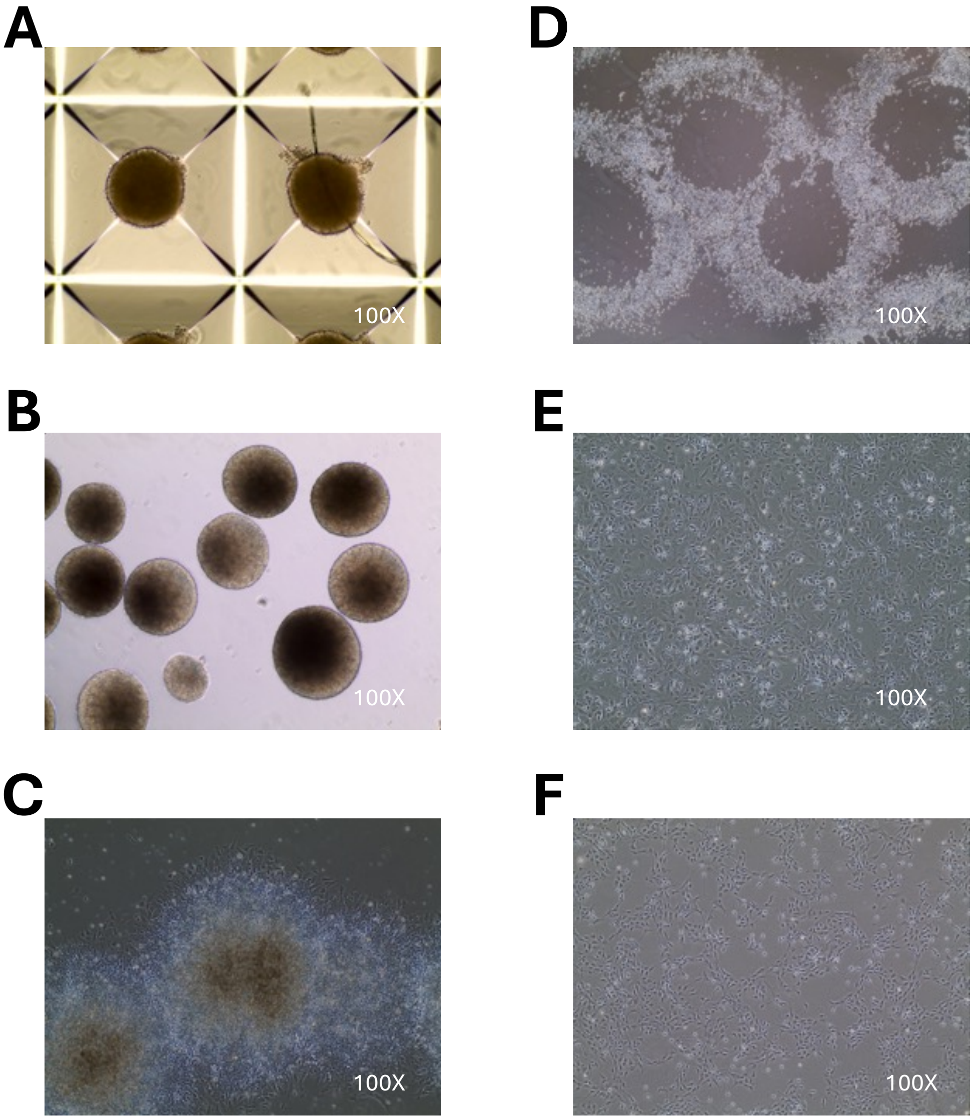


**Figure S4 Sequential stages of neural progenitor cell (NPC) generation via directed differentiation.** Bright-field images at 100× magnification showing the progressive steps of the differentiation protocol: **(A)** embryoid bodies (EBs), **(B)** EBs after medium replacement, **(C)** mature neural rosettes, **(D)** selection of rosettes, **(E)** first passage (P1) NPCs, and **(F)** third passage (P3) NPCs representing a stable population of mature NPCs.

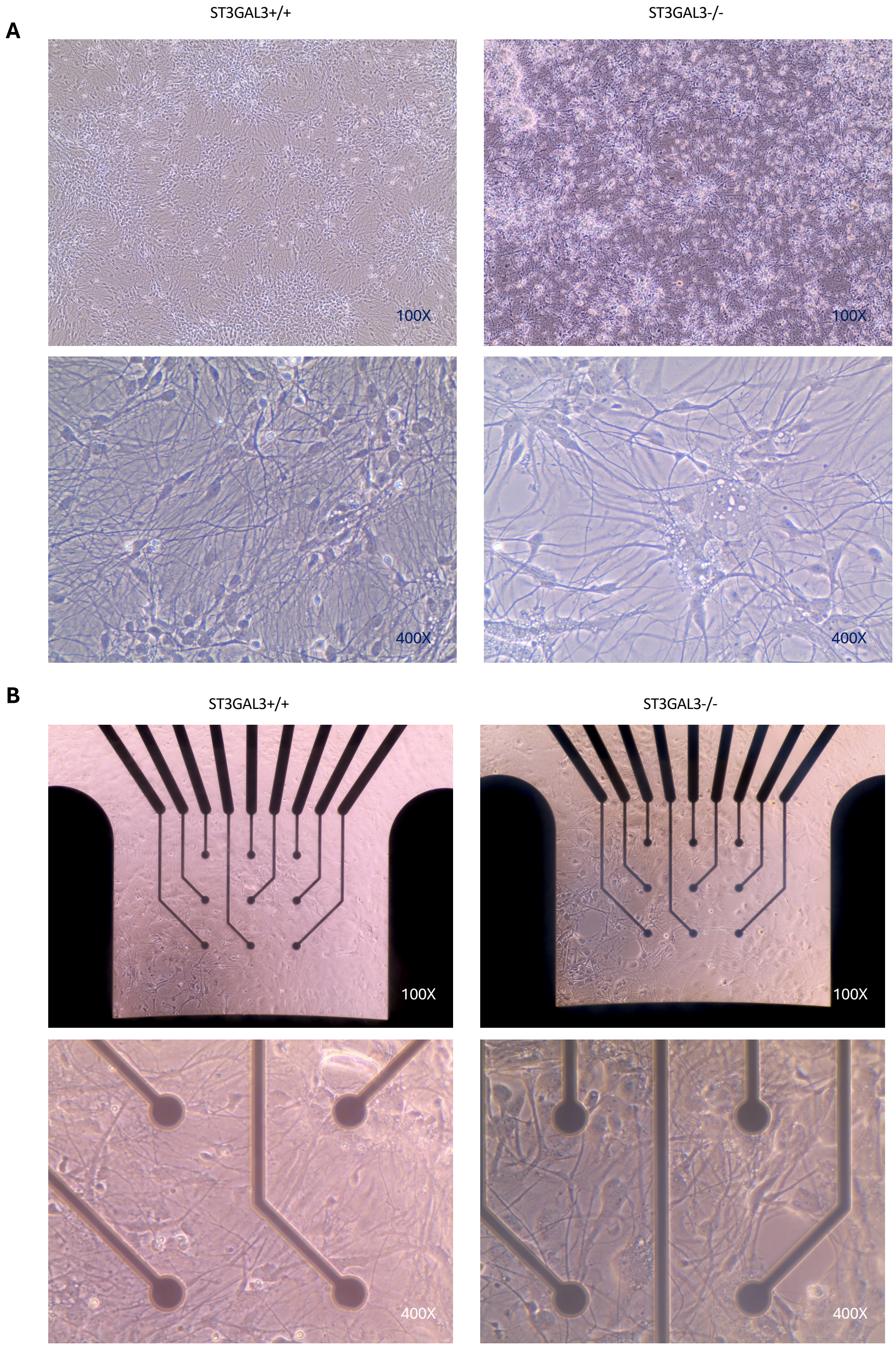


**Figure S5. Representative bright-field images of neurons derived from directed differentiation of iPSCs. (A)** *ST3GAL3*$^{-/-}$ (KO) and isogenic *ST3GAL3*$^{+/+}$ (WT) heterogeneous cortical neurons cultured on standard 24-well plates at 100× and 400× magnification, showing characteristic neuronal morphology with extended neurites. **(B)** *ST3GAL3*$^{-/-}$ (KO) and isogenic *ST3GAL3*$^{+/+}$ (WT) neurons cultured on 6wellMEA200/30iR-Ti-rcr microelectrode array (MEA) chips at 100× and 400× magnification, illustrating neuronal networks growing on microelectrode channels.

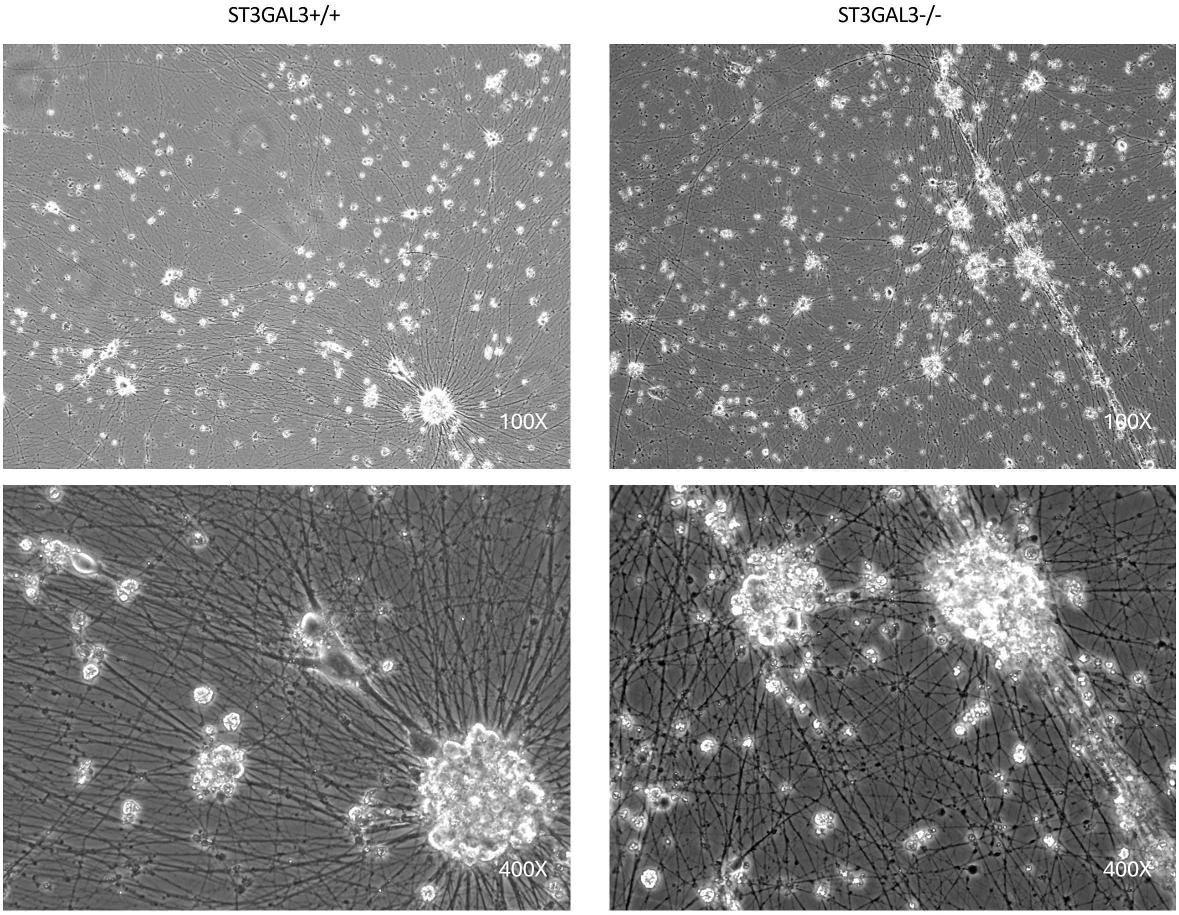


**Figure S6. Representative bright-field images of neurons derived from induced differentiation of iPSCs.** *ST3GAL3*$^{-/-}$ (KO) and isogenic *ST3GAL3*$^{+/+}$ (WT) Glutamatergic/GABAergic co-cultures on standard 24-well plates at 100× and 400× magnification, showing characteristic neuronal morphology with extended neurites.

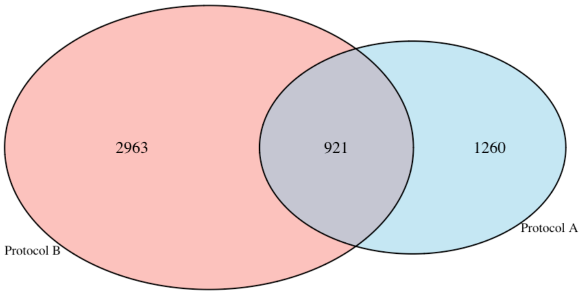


**Figure S7. Overlap of differentially expressed genes (DEGs) identified with DESeq2 between differentiation protocols.** Venn diagram showing the number of DESeq2-identified DEGs (padj < 0.01, | $\log_2$FC | > 2) in *ST3GAL3*$^{-/-}$ (KO) versus *ST3GAL3*$^{+/+}$ (WT) neurons generated using protocol A (blue) and protocol B (red). A total of 2,181 DEGs were identified in protocol A and 3,884 in protocol B, with 921 genes consistently differentially expressed across both protocols.

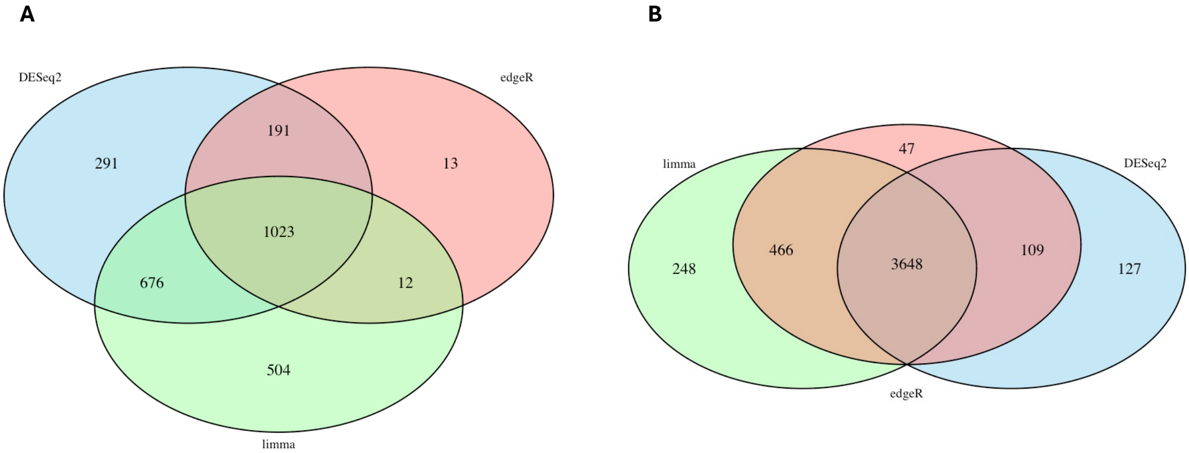


**Figure S8. Overlap of differentially expressed genes (DEGs) identified with DESeq2, limma-voom, and edgeR.** Venn diagrams showing the number of DEGs (adjusted P < 0.01, | $\log_2$FC | > 2) *ST3GAL3*$^{-/-}$ (KO) versus *ST3GAL3*$^{+/+}$ (WT) neurons identified via DESeq2, edgeR, and limma-Voom for **(A)** protocol A and **(B)** protocol B. A total of 1023 DEGs were identified in protocol A according to all three statistical methods, and 3,648 in protocol B.

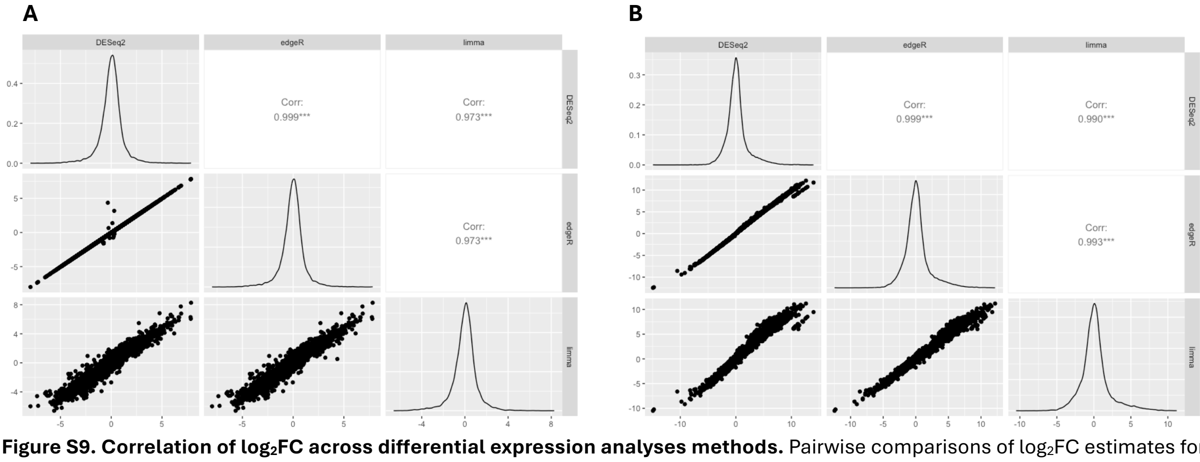


**Figure S9. Correlation of $\log_2$FC across differential expression analyses methods.** Pairwise comparisons of $\log_2$FC estimates for *ST3GAL3*$^{-/-}$ (KO) versus *ST3GAL3*$^{+/+}$ (WT) neurons obtained using DESeq2, edgeR, and limma-voom for **(A)** protocol A and **(B)** protocol B. Scatterplots in the lower panels show gene-wise correlations between methods, with density plots of $\log_2$FC distributions on the diagonal. Pearson correlation coefficients (*$p < 0.001$*) are reported in the upper panels, demonstrating very high concordance across methods (protocol A: r = 0.973–0.999; protocol B: r = 0.990–0.999).

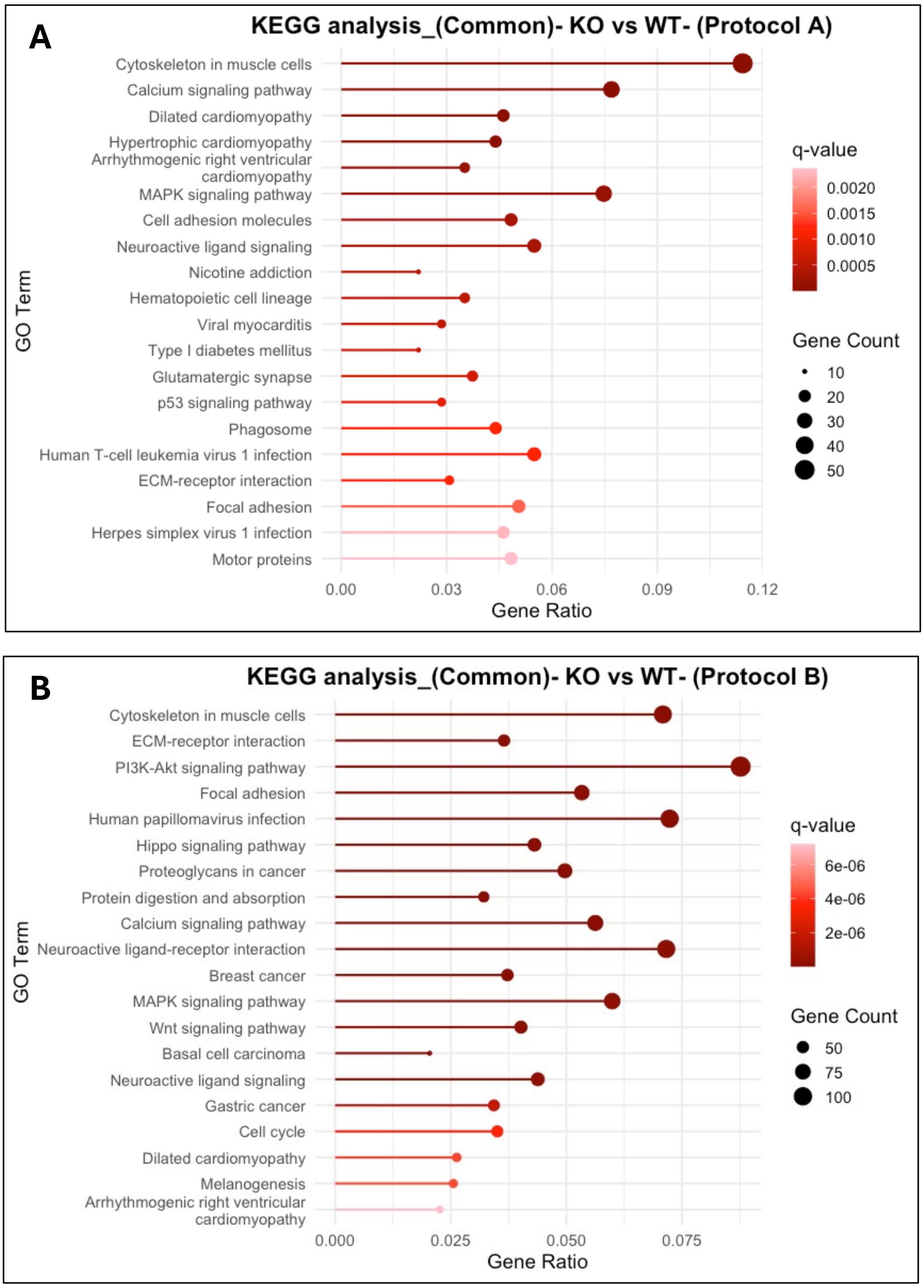


**Figure S10. KEGG pathway enrichment analysis of DEGs in *ST3GAL3*$^{-/-}$ (KO) versus *ST3GAL3*$^{+/+}$ (WT) neurons.** Bubble plots showing the top 20 enriched KEGG terms for DEGs identified with DESeq2, limma-voom, and edgeR in **(A)** protocol A and **(B)** protocol B. Pathways are ranked according to enrichment significance (q-value). The x-axis represents the gene ratio (proportion of DEGs annotated to the GO term), while bubble size corresponds to the number of DEGs and bubble colour reflects the q-value.

A

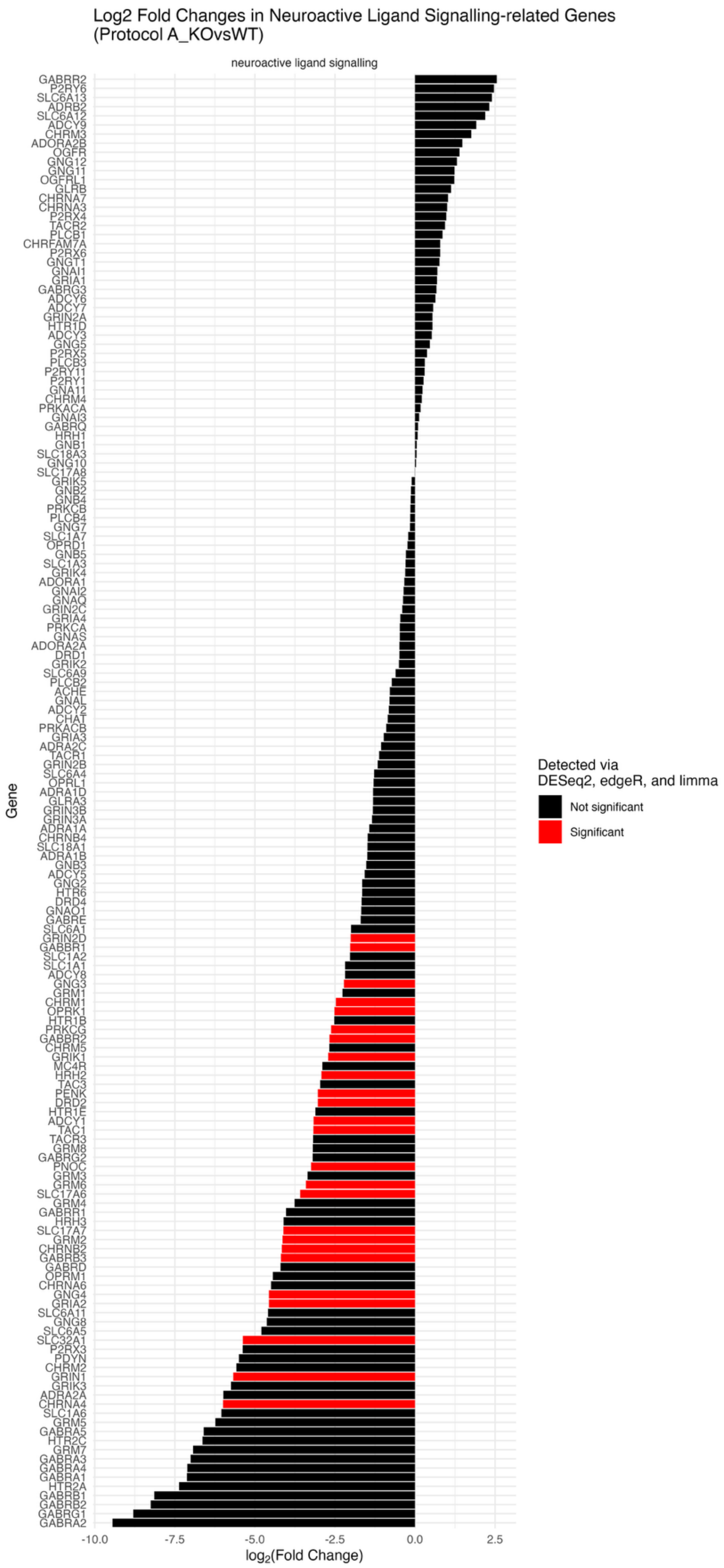
Log2 Fold Changes in Neuroactive Ligand Signalling-related Genes
(Protocol A_KOvsWT)
neuroactive ligand signalling
Gene
GABRR2
P2RY6
SLC6A13
ADRB2
SLC6A12
ADCY9
CHRM3
ADORA2B
OGFR
GNG12
GNG11
OGFRL1
GLRB
CHRNA7
CHRNA3
P2RX4
TACR2
PLCB1
CHRFAM7A
P2RX6
GNGT1
GNAI1
GRIA1
GABRG3
ADCY6
ADCY7
GRIN2A
HTR1D
ADCY3
GNG5
P2RX5
PLCB3
P2RY11
P2RY1
GNA11
CHRM4
PRKACA
GNAI3
GABRQ
HRH1
GNB1
SLC18A3
GNG10
SLC17A8
GRIK5
GNB2
GNB4
PRKCB
PLCB4
GNG7
SLC1A7
OPRD1
GNB5
SLC1A3
GRIK4
ADORA1
GNAI2
GNAQ
GRIN2C
GRIA4
PRKCA
GNAS
ADORA2A
DRD1
GRIK2
SLC6A9
PLCB2
ACHE
GNAL
ADCY2
CHAT
PRKACB
GRIA3
ADRA2C
TACR1
GRIN2B
SLC6A4
OPRL1
ADRA1D
GLRA3
GRIN3B
GRIN3A
ADRA1A
CHRNB4
SLC18A1
ADRA1B
GNB3
ADCY5
GNG2
HTR6
DRD4
GNAO1
GABRE
SLC6A1
GRIN2D
GABBR1
SLC1A2
SLC1A1
ADCY8
GNG3
GRM1
CHRM1
OPRK1
HTR1B
PRKCG
GABBR2
CHRM5
GRIK1
MC4R
HRH2
TAC3
PENK
DRD2
HTR1E
ADCY1
TAC1
TACR3
GRM8
GABRG2
PNOC
GRM3
GRM6
SLC17A6
GRM4
GABRR1
HRH3
SLC17A7
GRM2
CHRNB2
GABRB3
GABRD
OPRM1
CHRNA6
GNG4
GRIA2
SLC6A11
GNG8
SLC6A5
SLC32A1
P2RX3
PDYN
CHRM2
GRIN1
GRIK3
ADRA2A
CHRNA4
SLC1A6
GRM5
GABRA5
HTR2C
GRM7
GABRA3
GABRA4
GABRA1
HTR2A
GABRB1
GABRB2
GABRG1
GABRA2
-10.0
-7.5
-5.0
-2.5
0.0
2.5
log2(Fold Change)
Detected via
DESeq2, edgeR, and limma
Not significant
Significant

**B**

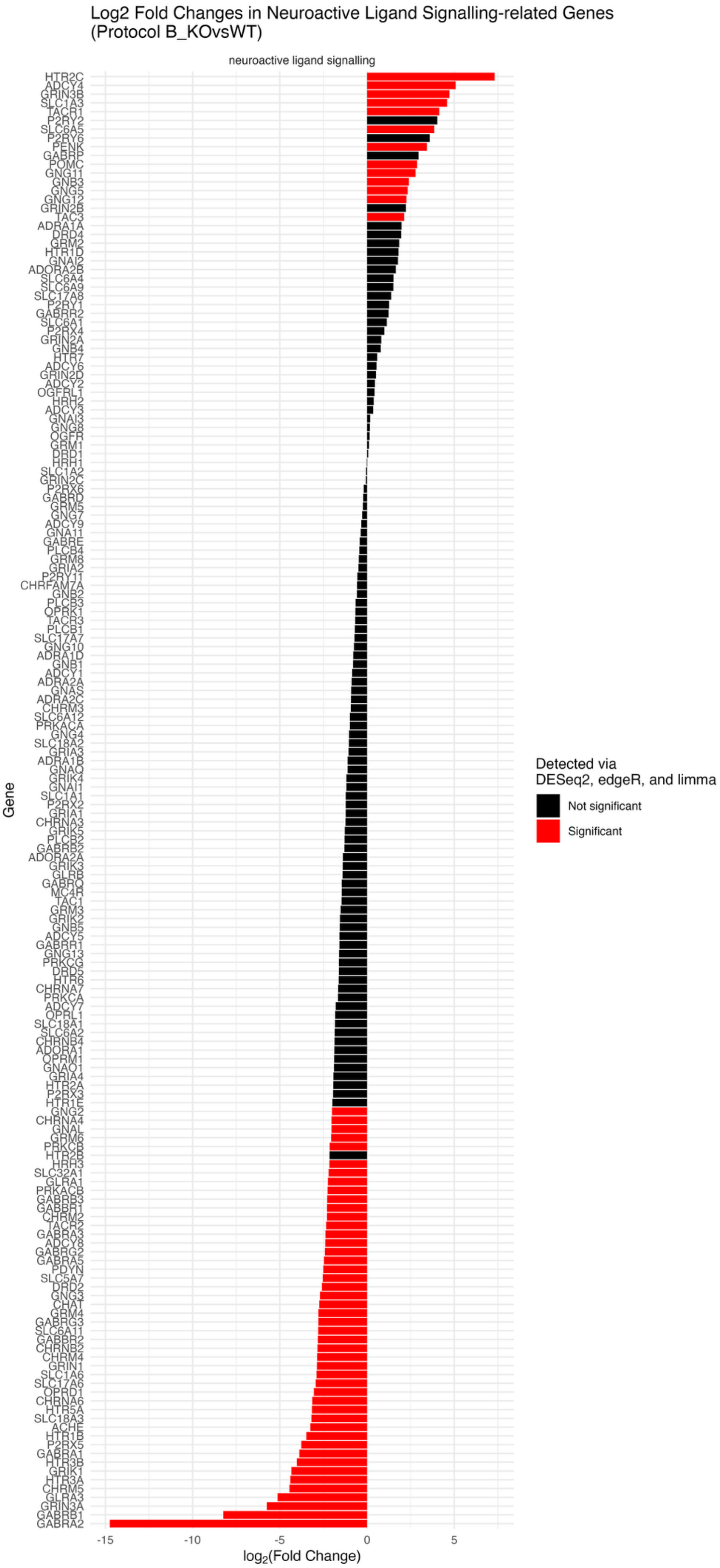

Log2 Fold Changes in Neuroactive Ligand Signalling-related Genes
(Protocol B_KOvsWT)
neuroactive ligand signalling
HTR2C
ADCY4
GRIN3B
SLC1A3
TACR1
P2RY12
SLC6A5
P2RY6
PENK
GABRP
POMC
GNG11
GNB3
GNG5
GNG12
GRIN2B
TAC3
ADRA1A
DRD4
GRM2
HTR1D
GNAI2
ADORA2B
SLC6A4
SLC6A9
SLC17A8
P2RY1
GABRR2
SLC6A1
P2RX4
GRIN2A
GNB4
HTR7
ADCY6
GRIN2D
ADCY2
OGFRL1
HRH2
ADCY3
GNAI3
GNG8
OGFR
GRM1
DRD1
HRH1
SLC1A2
GRIN2C
P2RX6
GABRD
GRM5
GNG7
ADCY9
GNA11
GABRE
PLCB4
GRM8
GRIA2
P2RY11
CHRFAM7A
GNB2
PLCB3
OPRK1
TACR3
PLCB1
SLC17A7
GNG10
ADRA1D
GNB1
ADCY1
ADRA2A
GNAS
ADRA2C
CHRM3
SLC6A12
PRKACA
GNG4
SLC18A2
GRIA3
ADRA1B
GNAQ
GRIK4
GNAI1
SLC1A1
P2RX2
GRIA1
CHRNA3
GRIK5
PLCB2
GABRB2
ADORA2A
GRIK3
GLRB
GABRQ
MC4R
TAC1
GRM3
GRIK2
GNB5
ADCY5
GABRR1
GNG13
PRKCG
DRD5
HTR6
CHRNA7
PRKCA
ADCY7
OPRL1
SLC18A1
SLC6A2
CHRNB4
ADORA1
OPRM1
GNAO1
GRIA4
HTR2A
P2RX3
HTR1E
GNG2
CHRNA4
GNAL
GRM6
PRKCB
HTR2B
HRH3
SLC32A1
GLRA1
PRKACB
GABRB3
GABBR1
CHRM2
TACR2
GABRA3
ADCY8
GABRG2
GABRA5
PDYN
SLC5A7
DRD2
GNG3
CHAT
GRM4
GABRG3
SLC6A11
GABBR2
CHRNB2
CHRM4
GRIN1
SLC1A6
SLC17A6
OPRD1
CHRNA6
HTR5A
SLC18A3
ACHE
HTR1B
P2RX5
GABRA1
HTR3B
GRIK1
HTR3A
CHRM5
GLRA3
GRIN3A
GABRB1
GABRA2
Gene
-15
-10
-5
0
5
log2(Fold Change)
Detected via
DESeq2, edgeR, and limma
Not significant
Significant

**Figure S11. Log$_2$FC of neuroactive ligand signalling pathway (KEGG hsa04080) genes in *ST3GAL3*$^{-/-}$ (KO) versus *ST3GAL3*$^{+/+}$ (WT) cultures. (A)** protocol A and **(B)** protocol B. Genes were considered significant when consistently detected by DESeq2, edgeR, and limma. with log$_2$FC values derived from DESeq2. Significant differential expression is highlighted in red, while black bars denote non-significant genes.

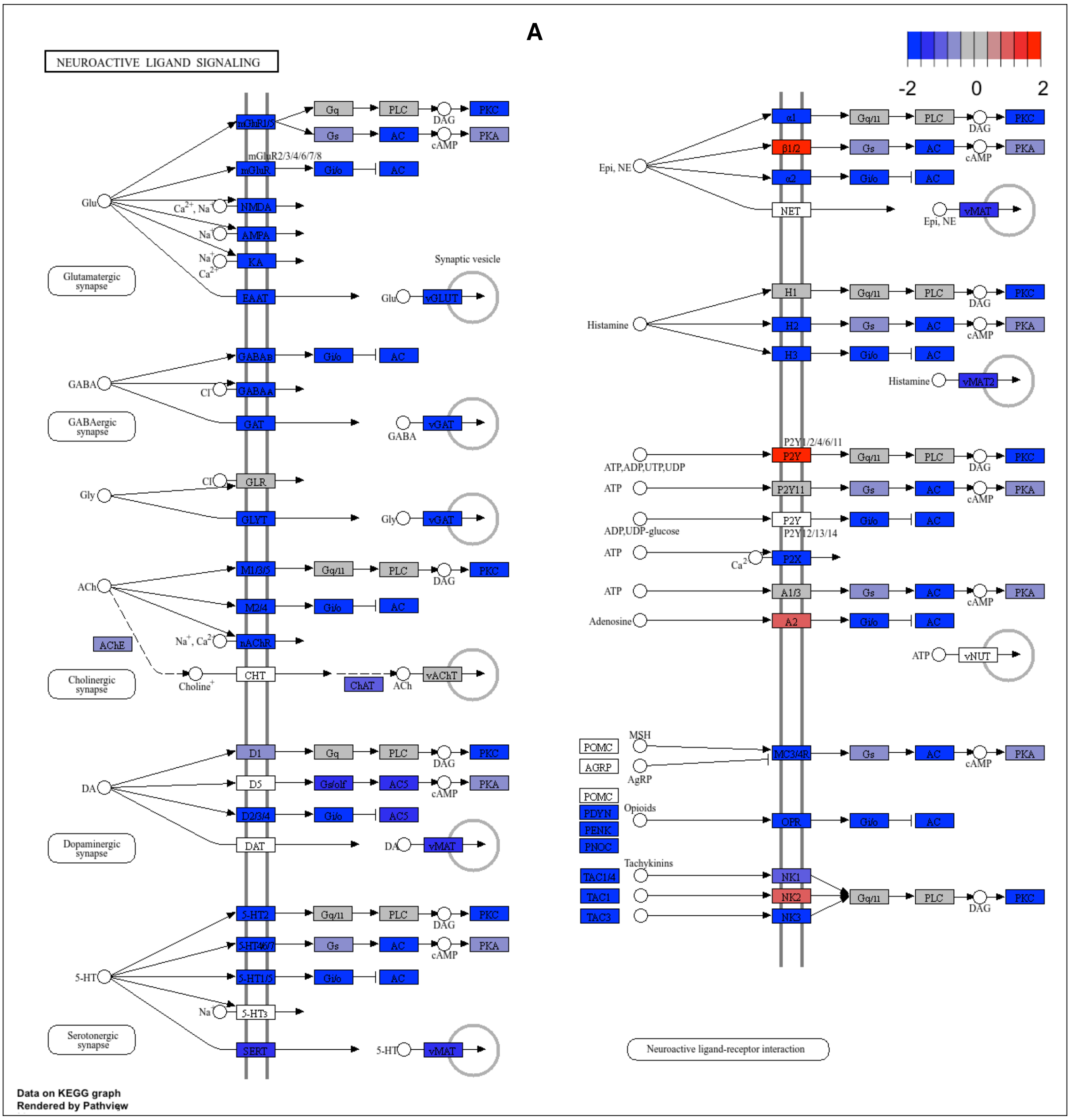

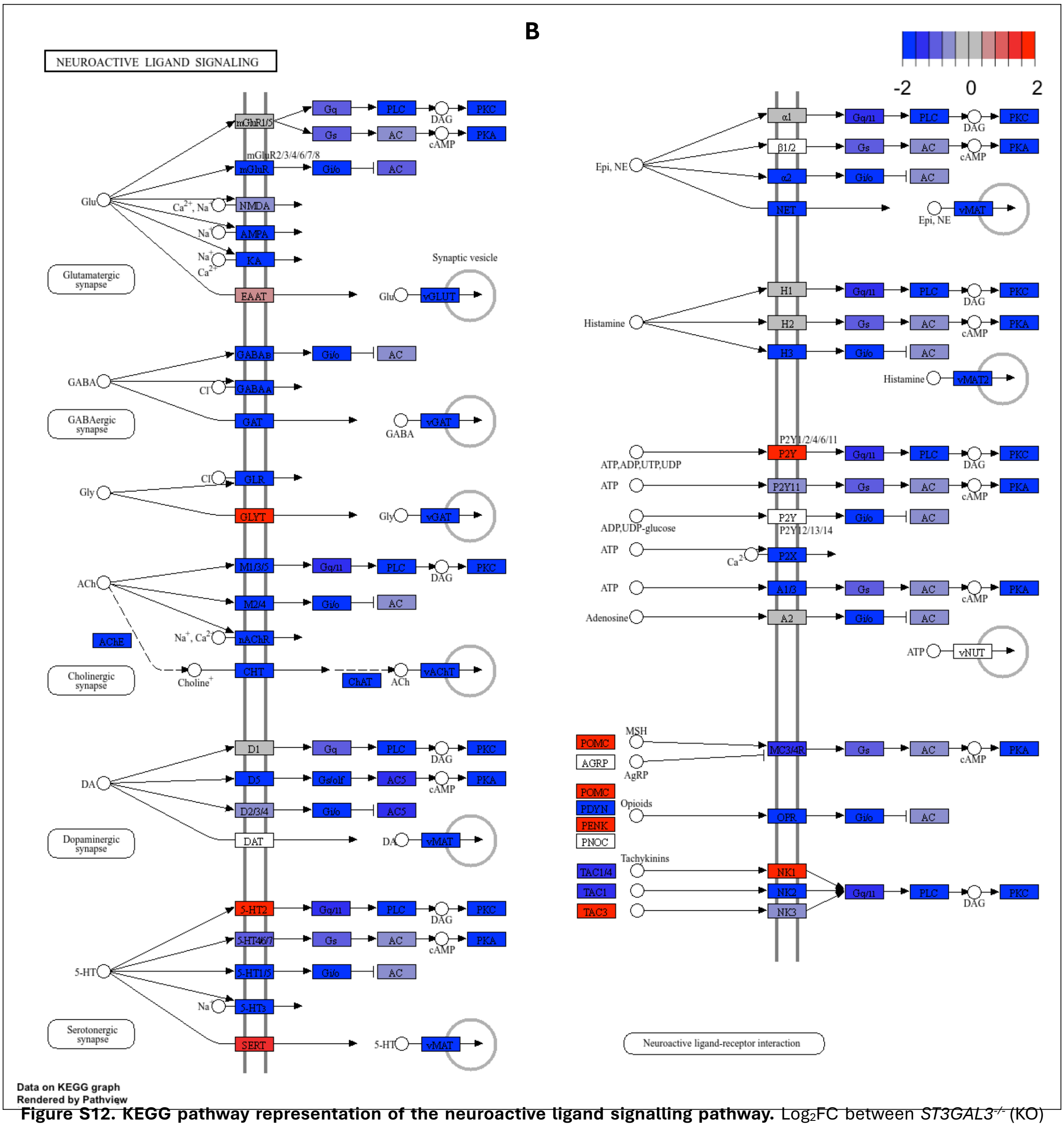


**Figure S12. KEGG pathway representation of the neuroactive ligand signalling pathway.** $Log_2FC$ between *ST3GAL3*$^{-/-}$ (KO) versus *ST3GAL3*$^{+/+}$ (WT) neurons differentiated via **(A)** protocol A or **(B)** protocol B were mapped onto their corresponding proteins in the KEGG *Neuroactive ligand signalling* pathway using Pathview. Boxes represent proteins encoded by the DEGs identified via DEseq2, coloured according to $log_2FC$ (red = upregulated, blue = downregulated in KO relative to WT).

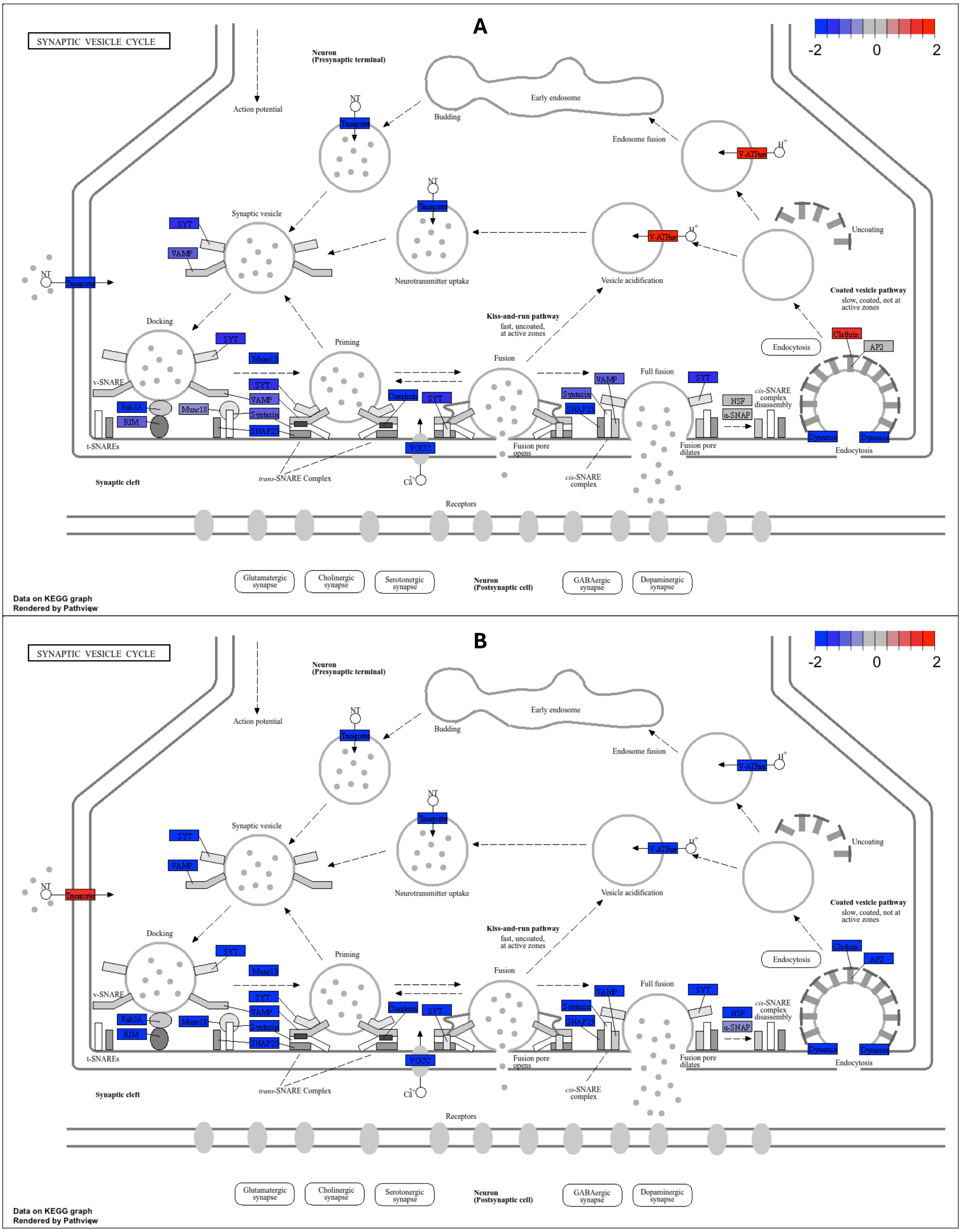


**Figure S13. KEGG pathway representation of Synaptic vesicle-cycle (hsa04721) genes.** $Log_2FC$ between *ST3GAL3*$^{-/-}$ (KO) versus *ST3GAL3*$^{+/+}$ (WT) neurons differentiated via **(A)** protocol A or **(B)** protocol B were mapped onto their corresponding proteins in the KEGG *Glutamatergic synapse* pathway using Pathview. Boxes represent proteins encoded by the DEGs identified via DEseq2, coloured according to $log_2FC$ (red = upregulated, blue = downregulated in KO relative to WT).

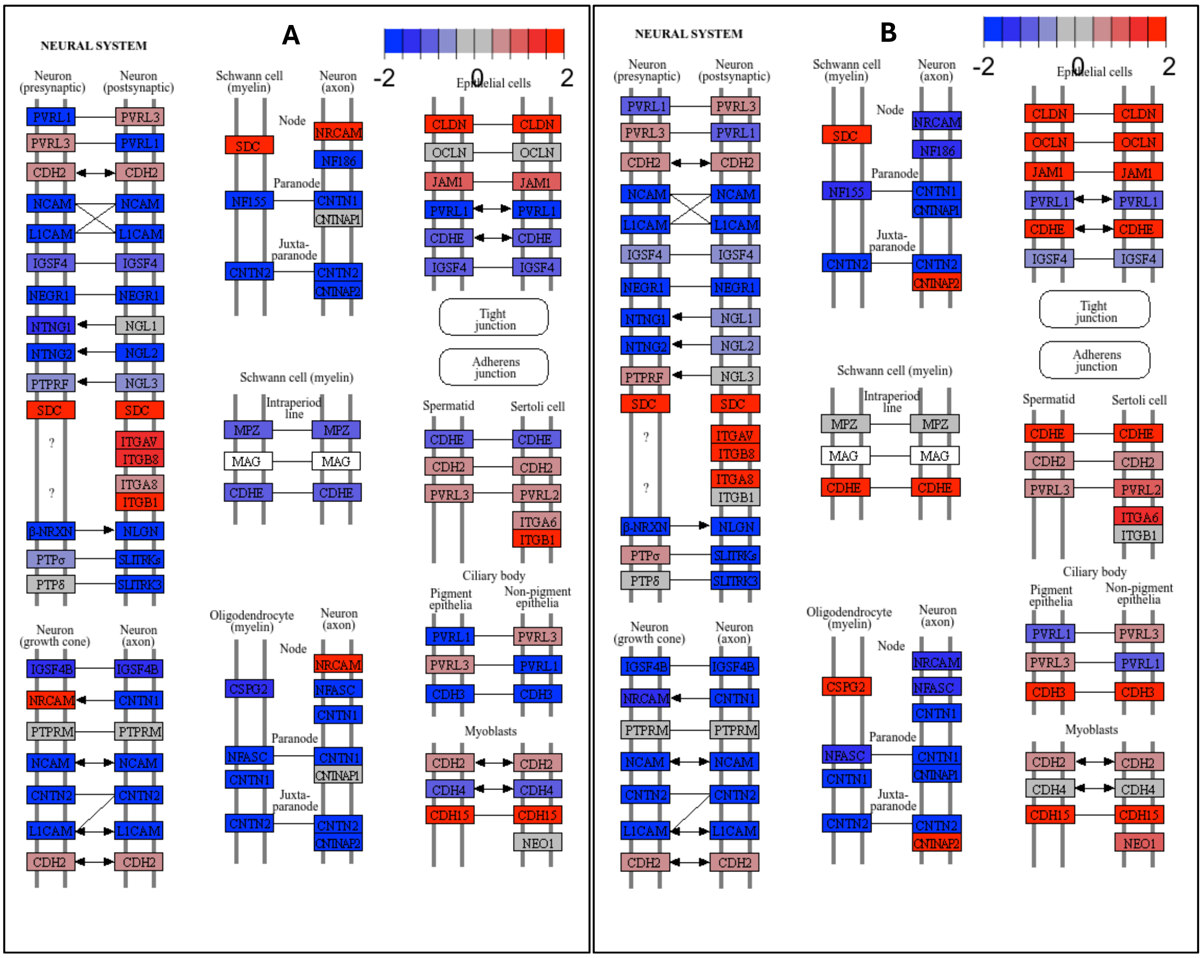


**Figure S14. KEGG pathway representation of Neral cell adhesion genes.** Gene set was extracted from cell adhesion molecules (hsa04514). $Log_2FC$ between *ST3GAL3*$^{-/-}$ (KO) versus *ST3GAL3*$^{+/+}$ (WT) neurons differentiated via **(A)** protocol A or **(B)** protocol B were mapped onto their corresponding proteins using Pathview. Boxes represent proteins encoded by the DEGs identified via DESeq2, coloured according to $log_2FC$ (red = upregulated, blue = downregulated in KO relative to WT).

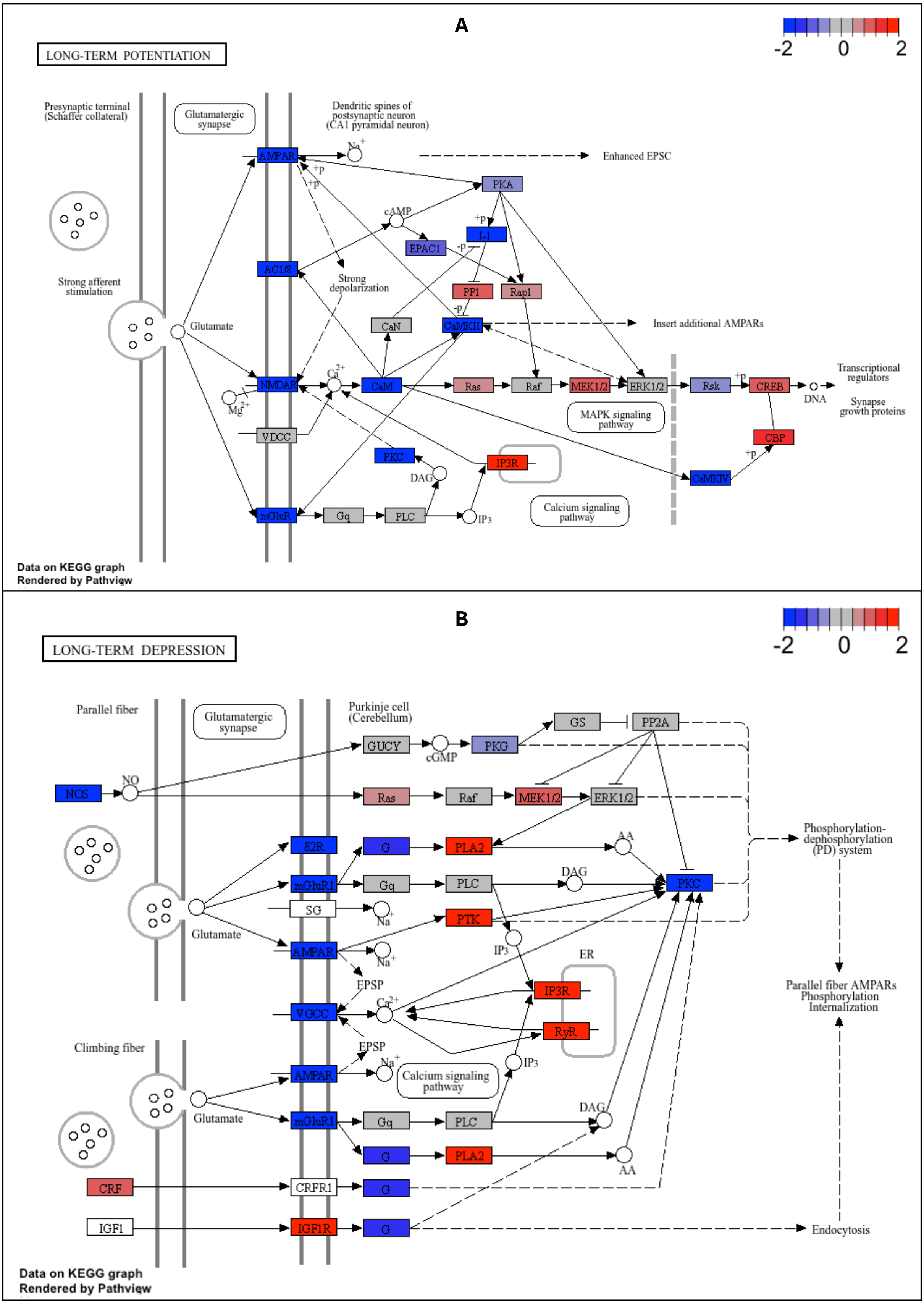


**Figure S15. KEGG pathway representation of LTD (hsa04730) and LTP (hsa04720) genes identified in protocol A.** Log$_2$FC between *ST3GAL3*$^{-/-}$ (KO) versus *ST3GAL3*$^{+/+}$ (WT) neurons differentiated via protocol A were mapped onto the **(A)** LTP (hsa04720) and **(B)** LTD (hsa04730) corresponding proteins using Pathview. Boxes represent proteins encoded by the DEGs identified via DEseq2, coloured according to log$_2$FC (red = upregulated, blue = downregulated in KO relative to WT).

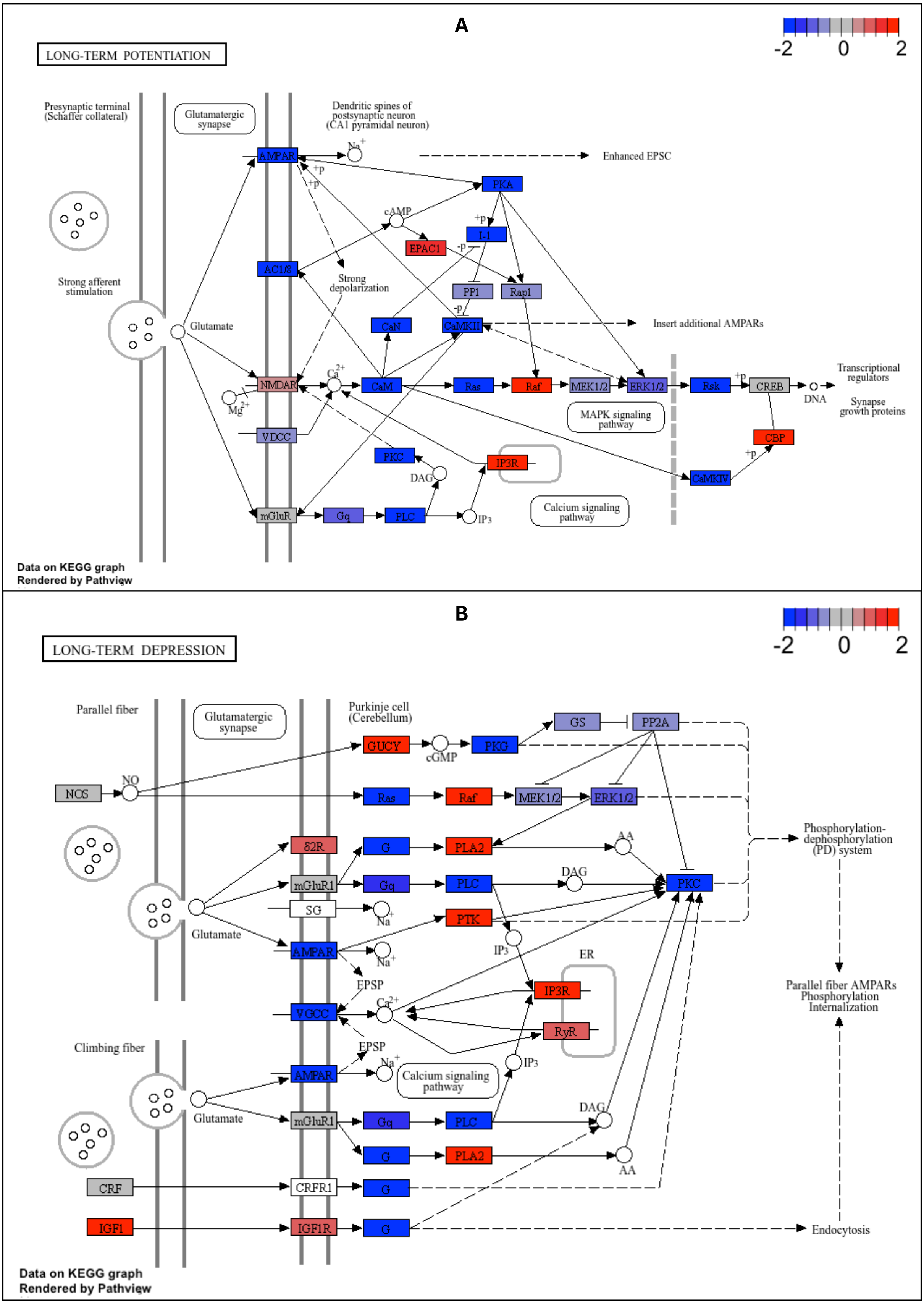


**Figure S16. KEGG pathway representation of LTD (hsa04730) and LTP (hsa04720) genes identified in protocol B.** $log_2$FC between *ST3GAL3*$^{-/-}$ (KO) versus *ST3GAL3*$^{+/+}$ (WT) neurons differentiated via protocol B were mapped onto the **(A)** LTP (hsa04720) and **(B)** LTD (hsa04730) corresponding proteins using Pathview. Boxes represent proteins encoded by the DEGs identified via DESeq2, coloured according to $log_2$FC (red = upregulated, blue = downregulated in KO relative to WT).

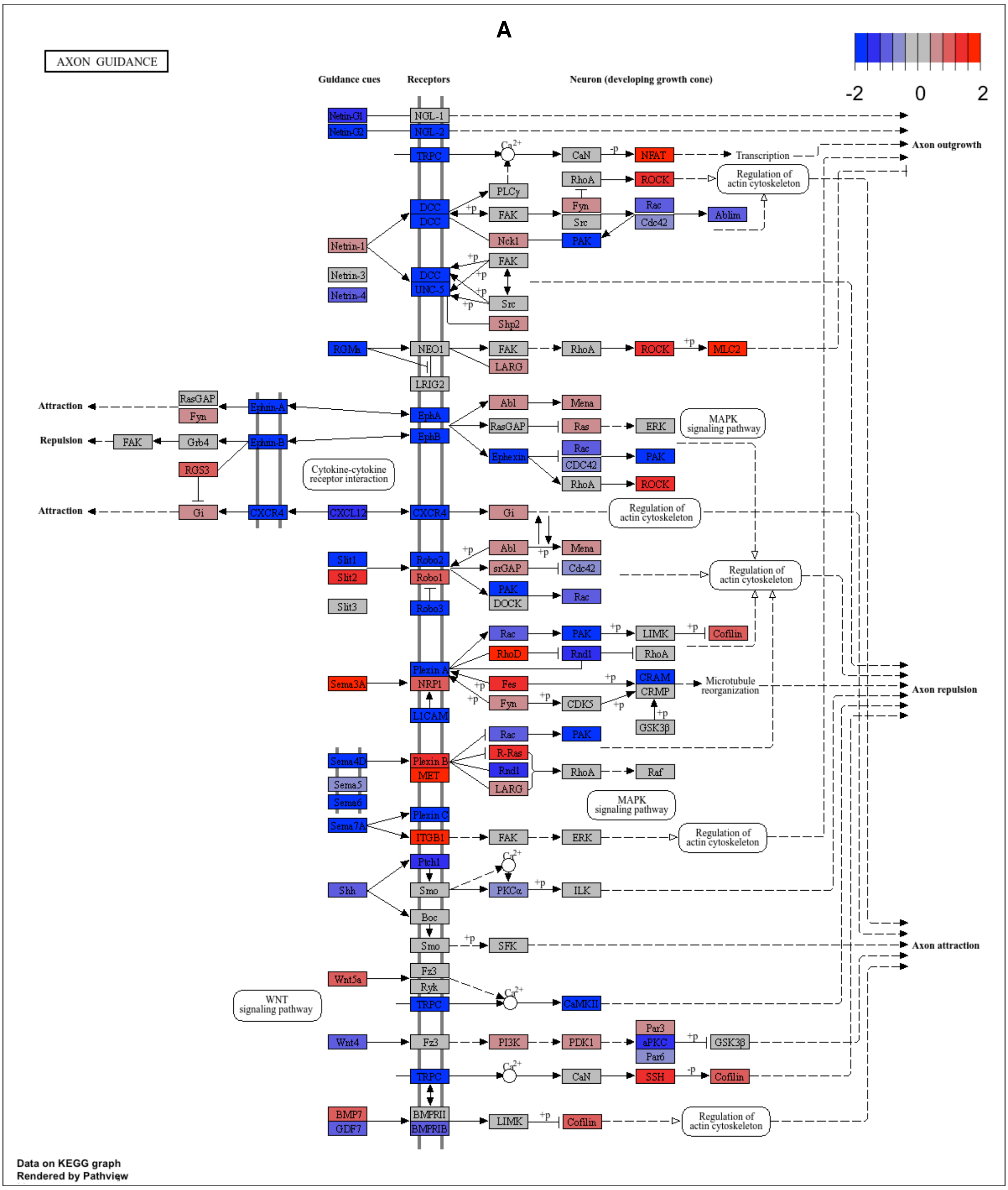
A
AXON GUIDANCE
-2
0
2
Guidance cues
Receptors
Neuron (developing growth cone)
Netrin-G1
Netrin-G2
NGL-1
NGL-2
TRPC
Ca2+
CaN
-p
NFAT
Transcription
Axon outgrowth
RhoA
ROCK
Regulation of actin cytoskeleton
PLCγ
Fyn
Rac
DCC
+p
FAK
Src
Cdc42
Ablim
Netrin-1
Nck1
PAK
UNC-5
Netrin-3
Netrin-4
Shp2
RGMa
NEO1
LARG
MLC2
LRIG2
RasGAP
Attraction
Ephrin-A
EphA
Abl
Mena
Repulsion
Grb4
Ephrin-B
EphB
Ras
ERK
MAPK signaling pathway
RGS3
Cytokine-cytokine receptor interaction
Ephexin
CDC42
Gi
CXCR4
CXCL12
Slit1
Slit2
Slit3
Robo2
Robo1
Robo3
srGAP
DOCK
LIMK
Cofilin
RhoD
Rnd1
Plexin A
NRP1
L1CAM
Sema3A
Fes
CRAM
CRMP
Microtubule reorganization
Axon repulsion
CDK5
GSK3β
Sema4D
Sema5
Sema6
Sema7A
Plexin B
MET
R-Ras
Raf
Plexin C
ITGB1
Ptch1
Shh
Smo
PKCα
ILK
Boc
SFK
Axon attraction
Wnt5a
Fz3
Ryk
CaMKII
WNT signaling pathway
Wnt4
PI3K
PDK1
Par3
aPKC
Par6
SSH
BMP7
GDF7
BMPRII
BMPRIB
Data on KEGG graph
Rendered by Pathview

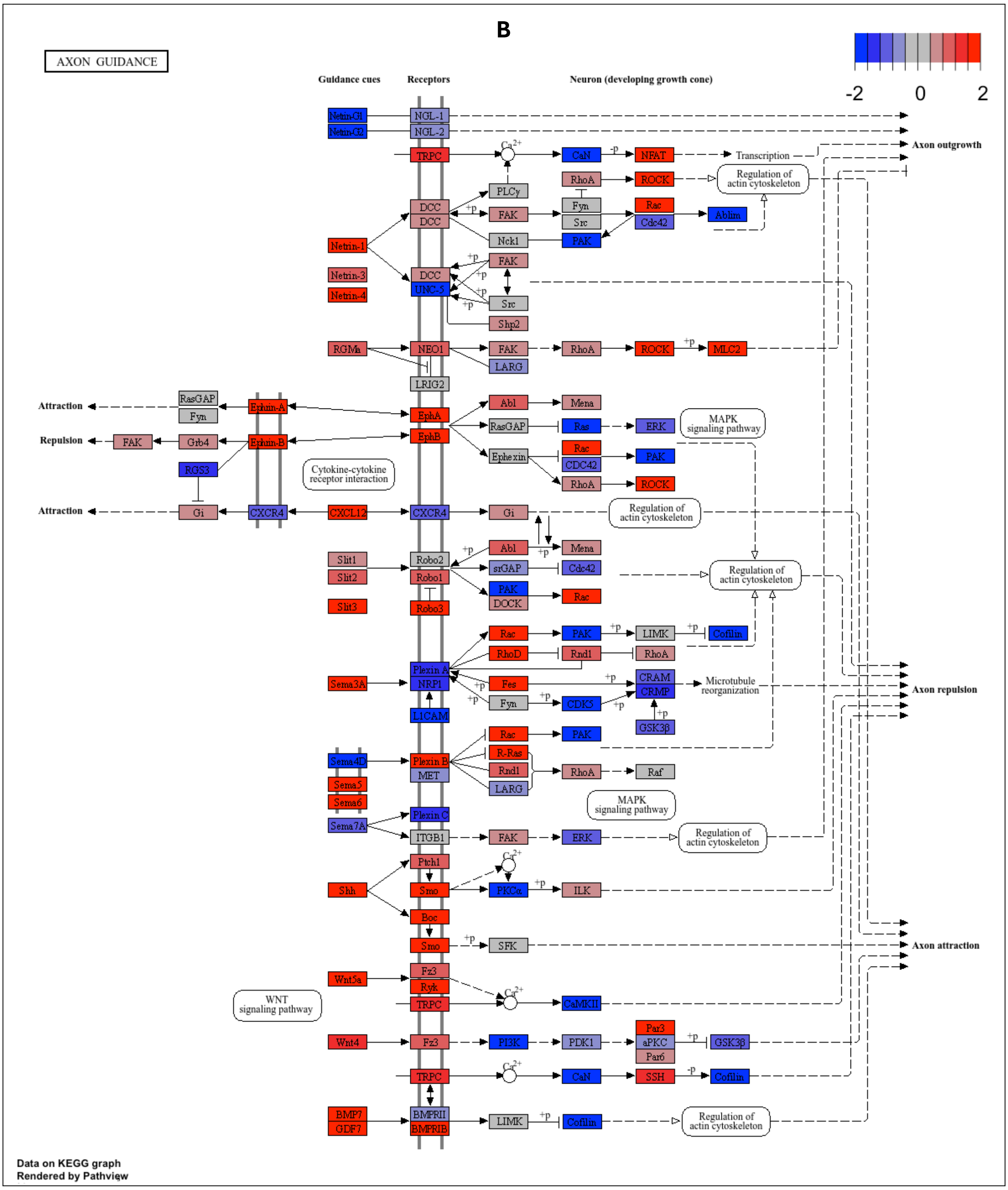


**Figure S17. KEGG pathway representation of axon guidance (hsa04360) genes.** Log$_2$FC between *ST3GAL3*$^{-/-}$ (KO) versus *ST3GAL3*$^{+/+}$ (WT) neurons differentiated via **(A)** protocol A or **(B)** protocol B were mapped onto their corresponding proteins using Pathview. Boxes represent proteins encoded by the DEGs identified via DESeq2, coloured according to Log$_2$FC (red = upregulated, blue = downregulated in KO relative to WT).

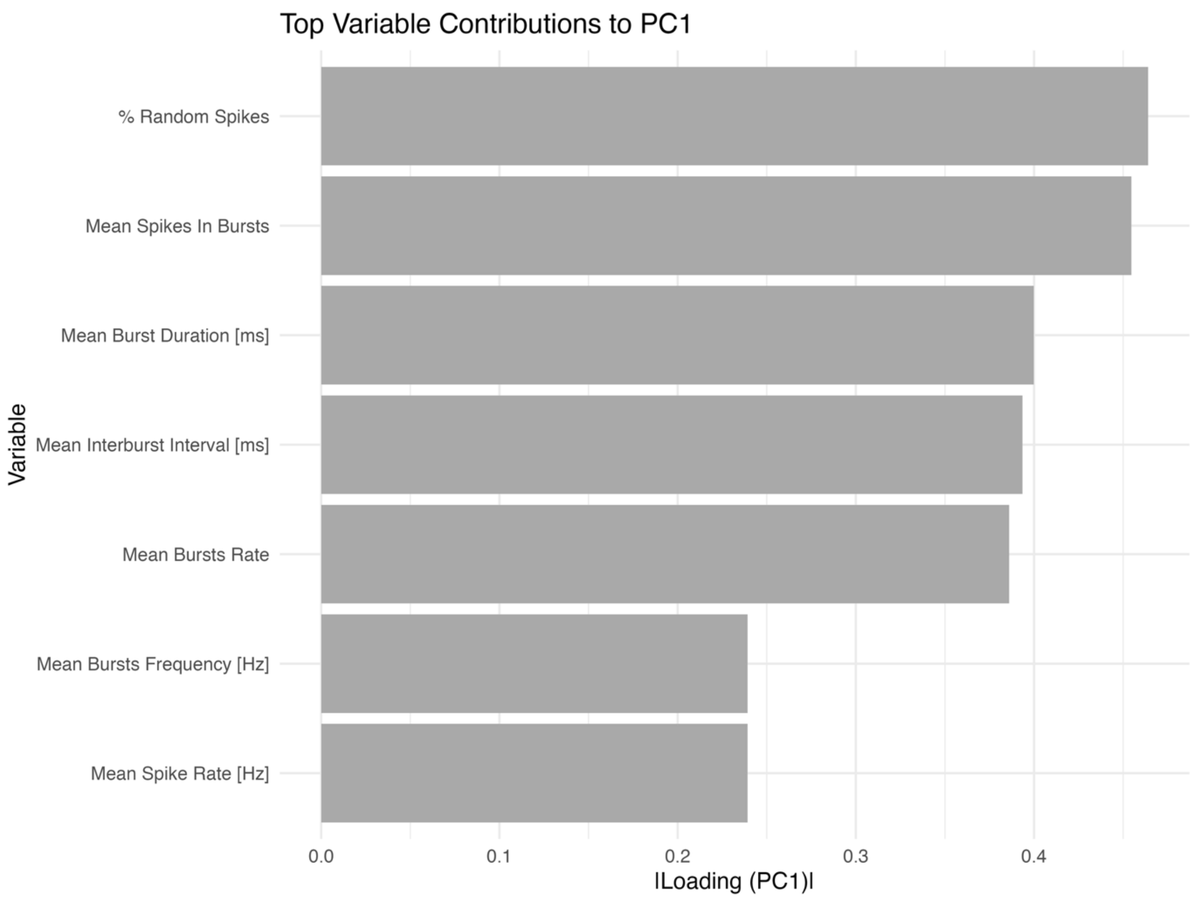


**Figure S18. Variable loadings on PC1 from the PCA of MEA-derived mean network parameters (Fig.21).** Horizontal bars show the absolute loading of each electrophysiological parameter on the first principal component (PC1); larger magnitudes indicate a stronger contribution to variance captured by PC1 (signs are not displayed). Parameters were averaged per biological replicate prior to PCA. ms milliseconds. Hz hertz.

### Table S7. Shapiro-Wilk Normality Test (per Genotype)

| Variable | ST3GAL3 KO | WT |
|---|---|---|
| **MBR** | 0.04655 | 0.11597 |
| **%RS** | 0.01767 | 0.18606 |
| **MBD** | 0.00106 | 0.00114 |
| **MSiB** | 0.00065 | 0.01001 |
| **MSR** | 0.00020 | 0.01865 |
| **MIBI** | 0.00702 | 0.00945 |
| **MBF** | 0.00020 | 0.01858 |

### Table S8. Wilcoxon Rank-Sum Test on mean MEA parameters (Fig. 21A-G)

| Variable | Cell line | Median | P-Value |
|---|---|---|---|
| **MBR** | ST3GAL3 KO | 16.666 | 0.15194 |
| | WT | 7.417 | |
| **%RS** | ST3GAL3 KO | 52.10 | 0.65563 |
| | WT | 57.55 | |
| **MBD** | ST3GAL3 KO | 340.850 | 0.33116 |
| | WT | 244.625 | |
| **MSiB** | ST3GAL3 KO | 6.35 | 0.65563 |
| | WT | 5.14 | |
| **MSR** | ST3GAL3 KO | 18.17 | 1 |
| | WT | 20.06 | |
| **MIBI** | ST3GAL3 KO | 19867.28 | 0.82376 |
| | WT | 22048.41 | |
| **MBF** | ST3GAL3 KO | 18.168 | 1 |
| | WT | 20.056 | |

**Table S8.** To compare mean electrophysiological parameters between *ST3GAL3*$^{-/-}$ (KO) versus *ST3GAL3*$^{+/+}$ (WT) cortical neuronal cultures, the Wilcoxon rank-sum test (also known as the Mann–Whitney U test) was applied. This non-parametric test was non-normal distribution of the data (See Table S7). For each parameter, the distributions between groups were assessed independently. Median values were compared across genotypes, and exact p-values were computed. No statistically significant differences were observed across all assessed metrics (al $p > 0.05$).

**A**

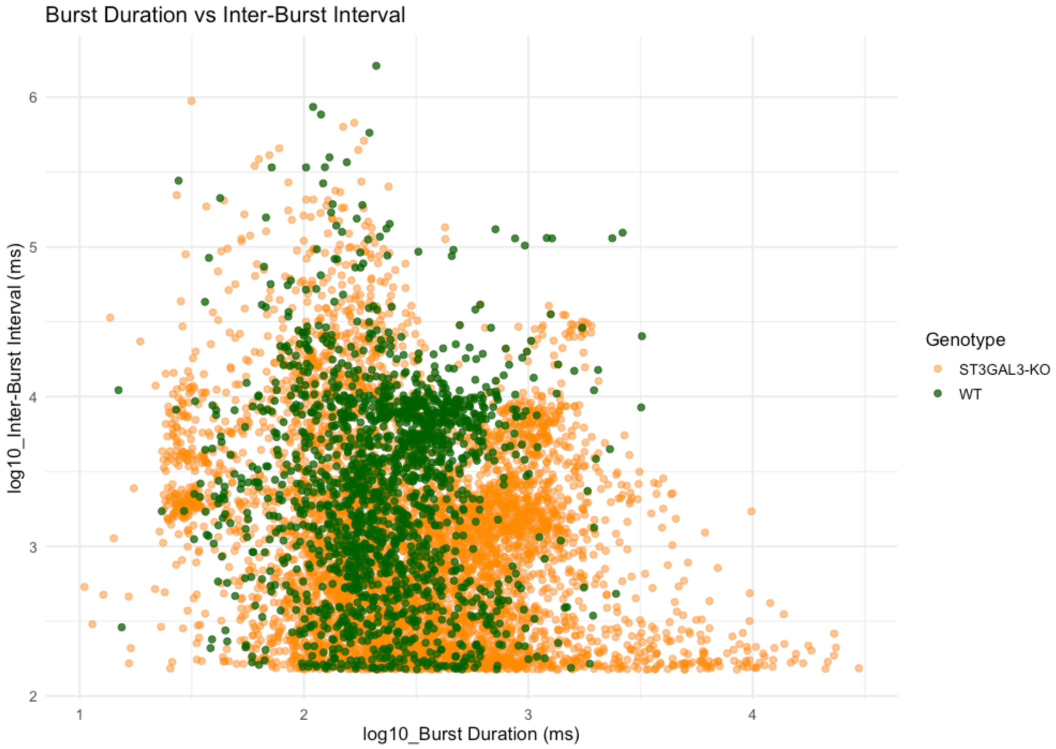


**B**

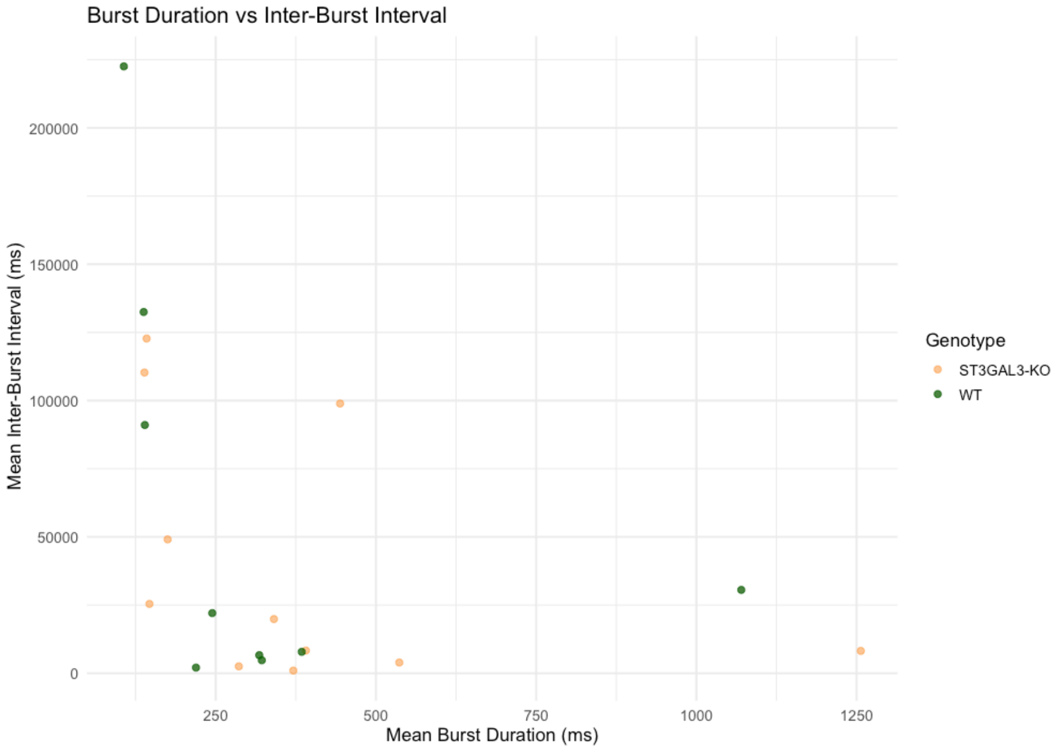


**Figure S19. Relationship between burst duration and inter-burst interval in ST3GAL3 KO and WT iPSC-derived cortical neurons.** **(A)** Scatterplot of individual burst events depicting the relationship between burst duration (BD; $log_{10}$-transformed, x-axis) and the subsequent inter-burst interval (IBI; $log_{10}$-transformed, y-axis), coloured by genotype. Each point represents a single burst recorded. **(B)** Same relationship examined at the level of mean values per recording well. Each point reflects the mean BD and mean IBI calculated across all burst events for a single replicate. Statistical testing using Pearson's correlation revealed no significant linear association between the two variables ($r$ = –0.388, $p$= 0.091; 95% CI: [–0.709, 0.066]). These results collectively indicate that, despite increased burst duration heterogeneity in KO cultures, this variation does not translate into systematic changes in network recovery time. ms milliseconds.

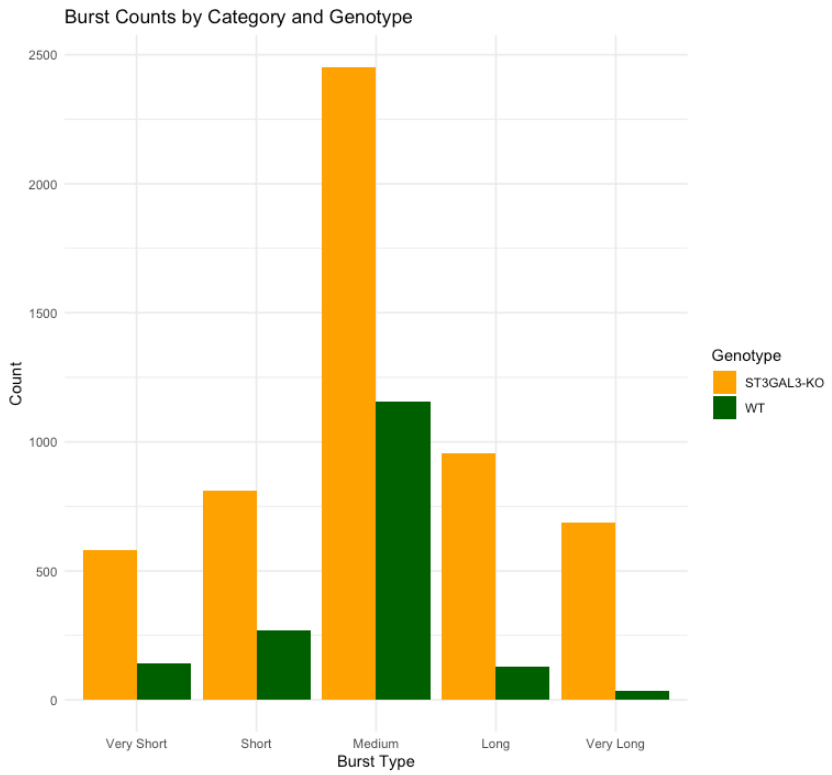


**Figure S20. Total burst counts per duration category across genotypes in iPSC-derived cortical neuronal networks.** Burst events from MEA recordings were classified into five duration categories based on percentile thresholds: Very Short (<10th percentile), Short (10th–25th), Medium (25th–75th), Long (75th–90th), and Very Long (>90th percentile). Bars represent the absolute count of bursts falling into each category for ST3GAL3 KO (orange) and WT (green) cultures.

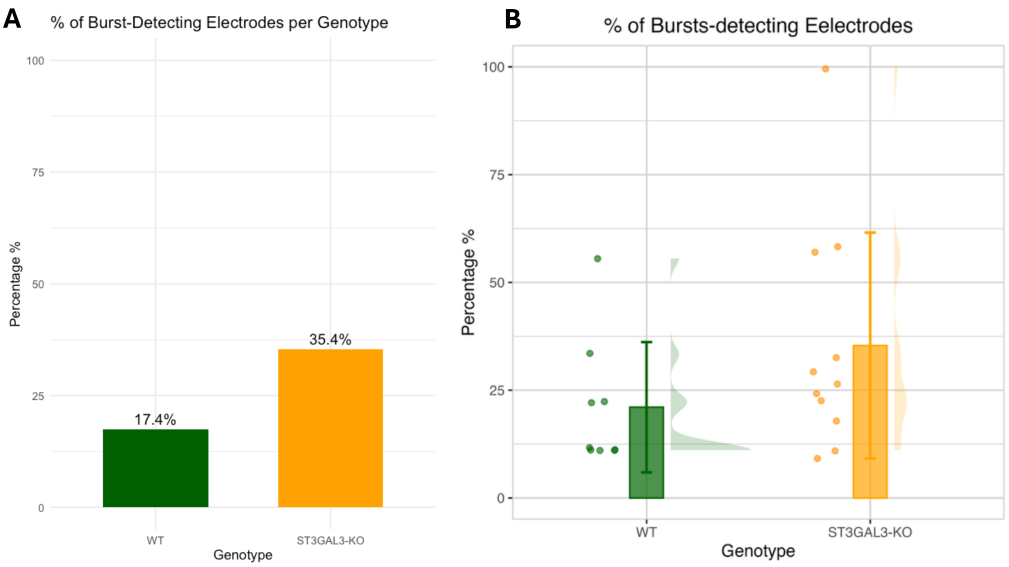


**Figure S21. Proportion of burst-detecting electrodes in ST3GAL3 KO and wild-type WT iPSC-derived cortical neuronal cultures. (A)**Bar plot showing the total percentage of burst-detecting electrodes across all wells per genotype. In aggregate. **(B)** Bar and jitter plot showing the percentage of burst-detecting electrodes per well across genotypes. Each point represents an individual well. While ST3GAL3 KO cultures exhibited a numerically higher percentage of burst-detecting electrodes per well, statistical comparison using the Wilcoxon rank-sum test did not reveal a significant difference between groups ($W$= 30, $p$= 0.137). Normality was assessed using the Shapiro-Wilk test and indicated non-normal distributions in both WT ($W$= 0.734, $p$= 0.0035) and KO ($W$= 0.806, $p$= 0.011), justifying the use of a non-parametric test.

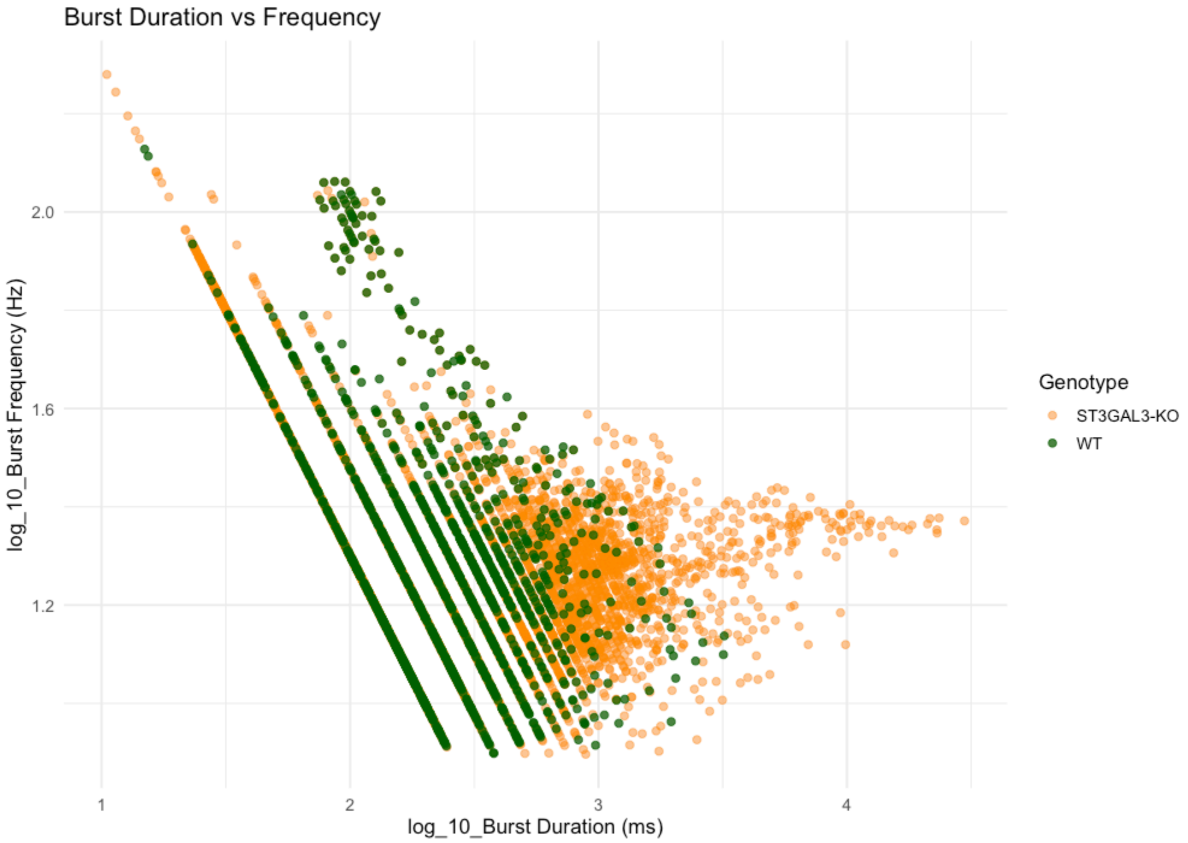


**Fig S22. Relationship between burst duration and burst frequency in ST3GAL3 KO and WT iPSC-derived cortical neurons. (A)** Scatterplot of individual burst events depicting the relationship between burst duration (BD; $log_{10}$-transformed, x-axis) and burst frequency (BF; $log_{10}$-transformed, y-axis), coloured by genotype. Each point represents a single burst recorded. ms milliseconds. Hz hertz.

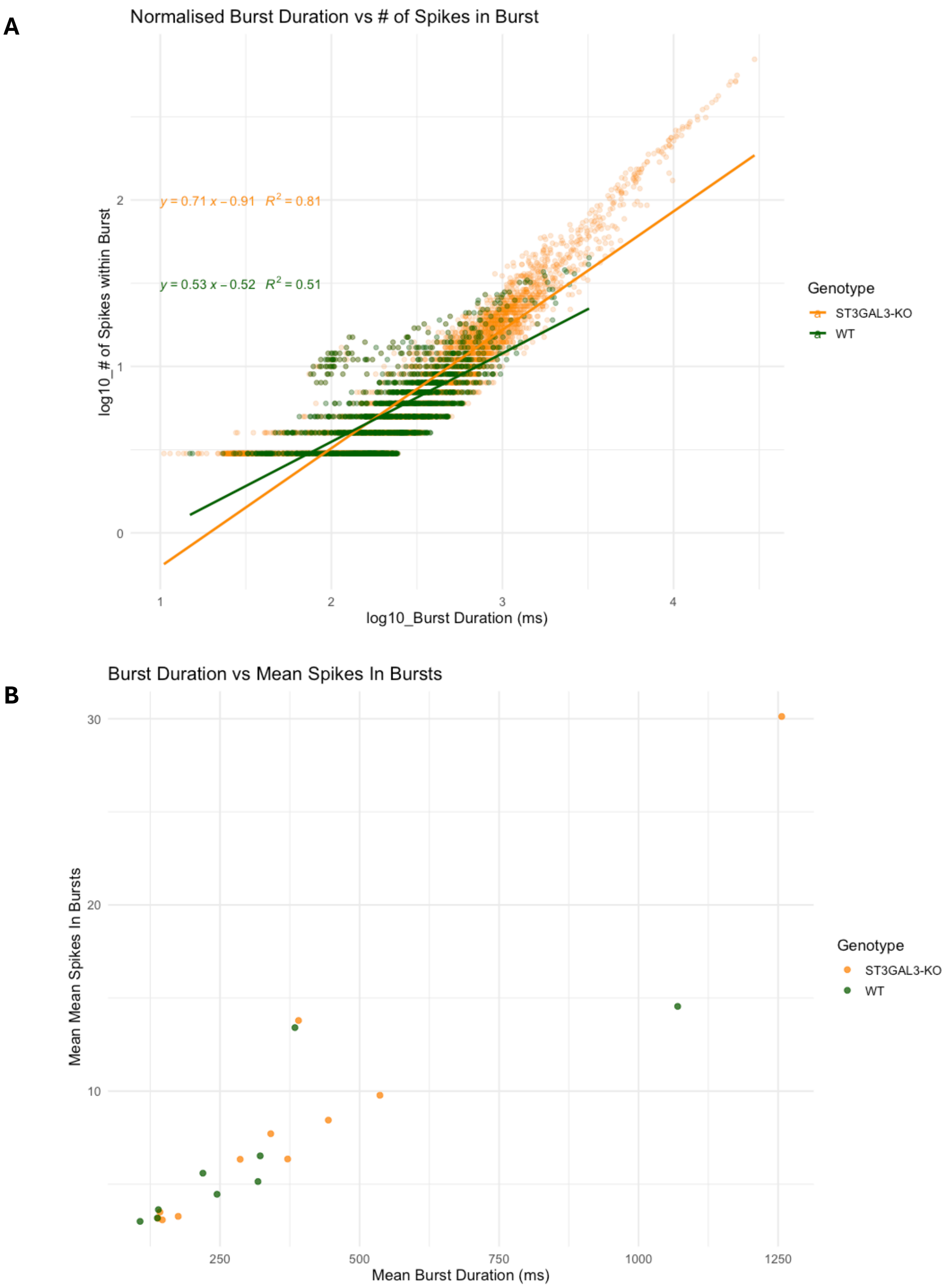


**Figure S23. Relationship between burst duration and number spikes in burst in ST3GAL3 KO and WT iPSC-derived cortical neurons. (A)** Scatterplot of individual burst events depicting the relationship between burst duration (BD; $log_{10}$-transformed, x-axis) and the number of spikes within each burst event (IBI; $log_{10}$-transformed, y-axis), coloured by genotype. Each point represents a single burst recorded. **(B)** Same relationship examined at the level of mean values per recording well. Each point reflects the mean BD and mean SIB calculated across all burst events for a single replicate. Statistical testing using Pearson's correlation revealed a strong significant linear association between the two variables (r = 0.9057, p = < 3.953e-08, 95% confidence interval of [-0.773, 0.963]). # number.
